# What Can We Count On? Performance of Microplate Cell Counting Assays in 2D Monolayer and 3D ECM-based In Vitro Tumour Models

**DOI:** 10.64898/2026.04.27.720021

**Authors:** Mahsa Vaezzadeh, Annemarie Nadort, Aleksandra Igrunkova, Vivienne S. Lee, Antonio Di Ieva, Benjamin Heng, Anna Guller

## Abstract

Accurate cell counting is essential in tissue engineering and cancer research. The ongoing transition towards advanced 3D in vitro tumour models raises a question about the validity of the standard cell counting protocols, particularly in the systems containing extracellular matrix-based scaffolds. Here, we provide a quantitative analysis of the performance of three popular plate reader-based cell counting/viability assays, such as the Alamar Blue, MTT, CellTiter Glo 3D assays, in 2D monolayer and 3D scaffold-based cultures of U251 human glioblastoma cells, including cell-laden Matrigel plugs, and original tissue engineering constructs based on the decellularised sheep brain scaffolds. We quantitatively characterized the assays’ linearity, precision, biological and technical reproducibility, proportionality, and inter-assay agreement. The study revealed that assays’ performance is highly platform-dependent, with 2D cultures allowing significantly more precise and reliable measurements than in 3D ECM scaffold-based cultures. The numerical results provided in this study can help researchers make informed decisions when working with 3D scaffold-based in vitro tumour models and for other tissue engineering purposes where precise cell counting is essential.

**ToC:** 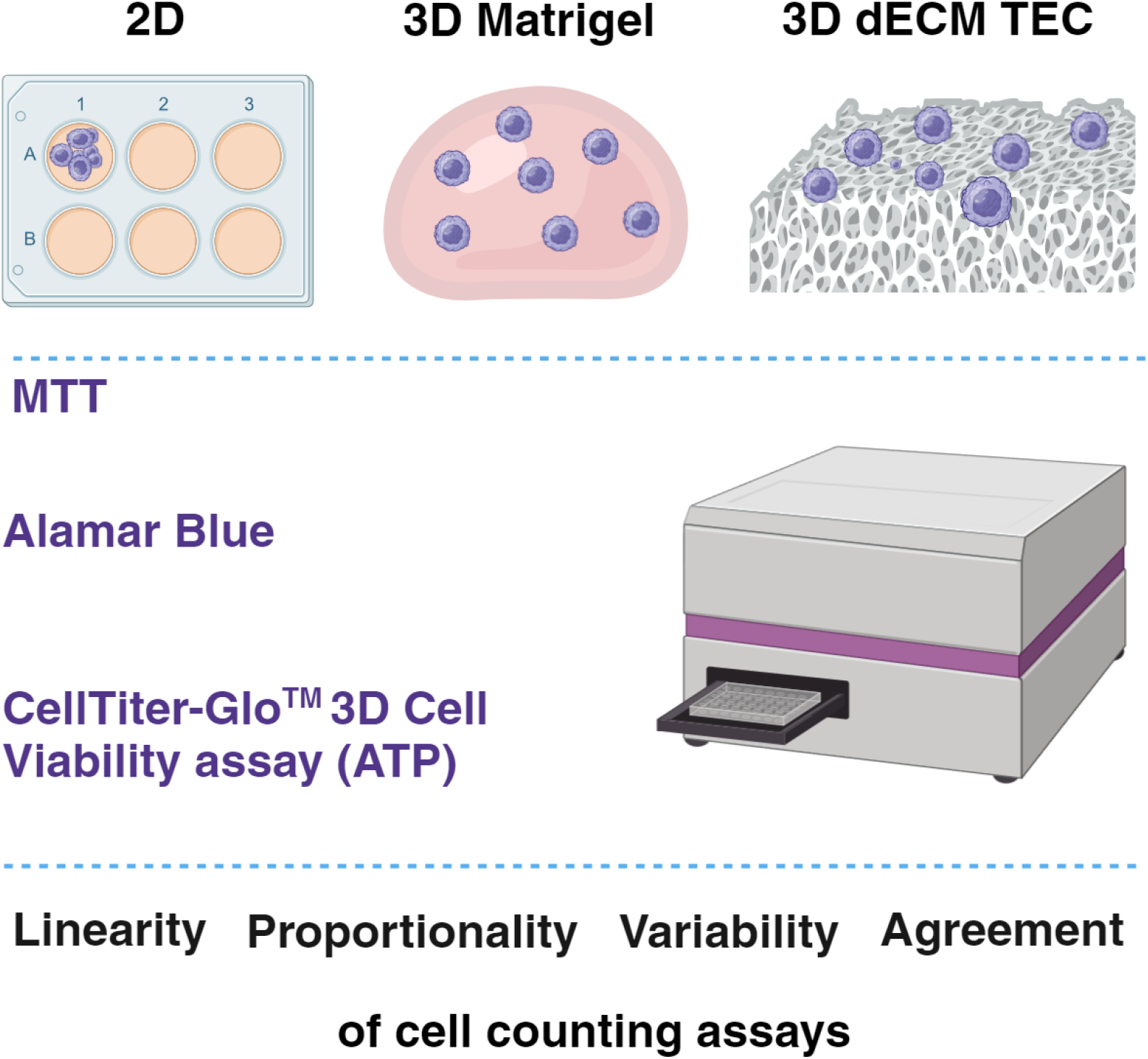

## 1. INTRODUCTION

The sustained growth of the tumour volume is associated with the abnormal proliferation of malignant cells and represents one of the key hallmarks of cancer ^[1]^. The changes in the tumour volume are associated with the cancer stage, progression rate, and prognosis and can reflect treatment responses ^[2]^. While the relationship between the cell number and the tumour’s size is complex and still not fully described ^[3]^, the quantitative analysis of cell numbers and viability is particularly important for cancer biology and oncology research. In vitro tumour models are the essential instrument in such research. The changes in cancer cell numbers in the tumours modelled in vitro reflect natural tumour growth as well as the effects of experimental treatments. Therefore, precise cell counting is one of the crucial requirements for the application of in vitro cultures as testbeds in oncology. Surprisingly, this critical demand is not well methodologically addressed yet.

There are several types of in vitro models of solid malignant tumours. In contrast to blood cancer, in such tumours cells’ growth is dependent on their attachment to the extracellular matrix (ECM) and on the cell-cell contacts. Therefore, the relevant in vitro models include the traditional two-dimensional (2D) monolayers of tumour cells cultured on the surfaces of specialized laboratory plastic vehicles and three-dimensional (3D) in vitro models. The last ones can be broadly classified as scaffold-free and scaffold-based systems. In the scaffold-free models, cell-cell attachment is encouraged by the absence of an adherent substrates. This results in the formation of multicellular aggregates, which are commonly termed spheroids, tumoroids, or tumour organoids. In contrast, in the scaffold-based models, cancer cells are provided with a 3D substrate (a scaffold) that to a certain extent mimics the native ECM and provides the adhesion cues. The scaffolds are usually presented by gel-like and solid biomaterials. The gel-like substrates allow mixing cells with the gel and immediately embedding them inside the simulated ECM or seeding cells on the gel surface. At the same time, solid scaffolds most commonly have fibro-porous structure, preserve their shape and the volume and allow seeding of the cells on the surface of the scaffold that may be flowed by the further migration of the cells inside the scaffold and its colonization.

Traditionally, the “gold standard” cell counting in vitro employs the trypan blue exclusion method combined with the detection of the live cells under microscope using a hemocytometer. Similarly, it can also be performed using automatic cell counters. However, the protocol includes the detachment of the cells from the substrate, which is often not acceptable for the purposes of experimental cancer research where the cell counting needs to be done without destroying the cell culture and/or in a high-throughput workflows. An alternative approach involves cell counting using plate readers via detection of light absorbance (optical density) or light emission (fluorescence or luminescence) of the sample. These optical signals are obtained by adding the tested cell culture with the reagents that react with the cellular components or metabolic products in a chemically defined way, while the reaction product has specific optical characteristics that are changing proportionally to the cell numbers. Most commonly, the plate reader-based cell counting methods rely on a few parameters such as amount of double-stranded DNA (dsDNA) or various measures of metabolic activity of the cells (Table 1).

**Table 1.** Comparison of the commonly used plate reader-based cell counting assays explored in the current study.

| Assay<br>(Manufacturer) | Biological principle | Substrate | Protocol | Detection<br>principle* | Sensitivity, Ref.** |
| --- | --- | --- | --- | --- | --- |
| MTT*<br>(Sigma-Aldrich) | Activity of dehydrogenases mostly in mitochondria and cytosol | Cell lysate | End-point | Abs. | 5,000 cell/well <sup>[4]</sup> - 25,000 cells per well, <sup>[5]</sup> |
| Alamar Blue<br>(Sigma Aldrich) | Activity of reductases in various cell compartments | Cell culture supernatant | End-point or continuous | Fl. or Abs. | 1,500 cells/well <sup>[5]</sup> – 2,000 cells/well <sup>[6]</sup> |
| CellTiter-Glo® 3D<br>(Promega) | Amount of ATP produced in mitochondria | Cell lysate | End-point | Lum. | 10 nM ATP, or 400 cells per well, <sup>[7]</sup> |

\*Abbreviations: MTT - Thiazolyl Blue Tetrazolium Bromide (#M5655, Sigma-Aldrich); Alamar Blue - Resazurin sodium salt (#199303, Sigma-Aldrich); CellTiter-Glo® 3D - CellTiter-Glo® 3D Cell Viability Assay (#G9681, G9682 and G9683, Promega); Fl. – Fluorescence, Abs. – Absorbance, Lum. – Luminescence. \*\*References are given for the low limits of the reliable detection observed experimentally.

It must be noted that in the literature the results of cell counting assays are also often interpreted in terms of cells’ viability, proliferation, or cytotoxicity of experimental treatments, depending on the study design. To make the message of our work more straightforward, here, we limited the interpretation of the assays as counting of viable cell, assuming that all assays reflect the changes in viable cells only, while may not be specifically revealing the live: dead cells’ ratios.

The methodology of cell counting in 3D cultures is under development ^[8]^. It is much more advanced for the spheroid-like models, which allow the direct measurements of the growing experimental microtumours by imaging and various plate-reader-based assays, as demonstrated elsewhere ^[9]^. However, for the scaffold-based models, such as the ones based on Matrigel, the most popular commercially available product mimicking the ECM of a highly vascularised tumour ^[10]^, and especially for the novel 3D in vitro tumour models that employ ECM-mimicking biomaterials, the protocols of cell counting and viability assays still need to be adapted and validated ^[8, 11]^. In contrast to multicellular spheroids or organoids, the scaffold-based tumour models do not significantly change in size, while the number of cells goes up or down. Then, the direct observation of the experimental tumour growth dynamics becomes challenging. In addition, the scaffolds are mostly non-transparent and light-scattering materials, implying that the cells’ counting and viability analysis may require confocal optical setups, increasing the time- and resource cost for such tests ^[11a]^. Only a few relatively high-throughput approaches for the analysis of cell viability in scaffold-based in vitro cultures have been demonstrated ^[11e, 12]^. This emphasises the importance of the development and validation of plate-reader-compatible cell counting and viability assays for the scaffold-based 3D tumour models. Recent publications have contributed to a better understanding this problem ^[13]^. Among the common findings of some of these studies, it was noted that reliable cell counting in scaffold-based 3D in vitro cell cultures may require a combination of several assays and adaptation for the specific tumour/cell type.

In 2018 and 2019, two new ISO standards on cell counting for biomedical research applications were published ^[14]^. These standards addressed the growing demand for reliable biomanufacturing, regenerative medicine and immunotherapy and recommended evaluation of cell counting methods by several quantitative performance measures. These measures included the testing of the assays’ precision by coefficients of variation (CVs), the examination of the assays’ linearity by comparing the coefficients of determination (R-squared, or R^2^) obtained through the linear regression analysis, as well as the studying of the proportionality indexes (PIs) that reflect to what extent the assays readings follow the changes in cell numbers. Bland-Altman analysis represented another recommended approach, specifically applicable for the comparison of the performance of two different assays in the same cell culture conditions. However, these standards are adapted for the work with suspension cell cultures or dispersions of cells. To the best of our knowledge, there have not been any reports on the application of such systematic analysis of the performance of several cell counting assays in 3D scaffold-based in vitro systems.

The current study quantitatively explores the performance of four established cell counting assays (see Table 1) in 2D conventional monolayer and in 3D ECM scaffold-based in vitro tumour models. Two ECM-mimicking materials were applied as scaffolds for cell culturing, including Matrigel and the fibro-porous scaffolds representing the ECM of decellularised sheep brain (dECM) ^[15]^. The selection of the brain-derived ECM as a scaffolding material was motivated by our focus ^[16]^ on glioblastoma research. This deadliest brain cancer still has very limited treatment options ^[17]^. Thus, reliable methodologies for cell counting for in vitro models of glioblastoma are essential for accelerating the improvement of clinical outcomes.

To address this challenge, here, we aimed to reveal whether and to what extent the cell counting approaches that employ plate readers are reliable in 3D ECM scaffold-based in vitro models of glioblastoma, in comparison with the matching 2D monolayer cell cultures.

## 2. MATERIALS AND METHODS

### 2.1. Study Design

The study design is schematically shown in Figure 1. We employed human glioblastoma cell line U251 cultured in three platforms, including conventional 2D monolayers and 3D models, where cells were embedded in Matrigel or seeded on the top of brain-derived dECM fibro-porous scaffolds. To examine the cell counting assays’ performance, two types of experiments were performed. In 1-day (static) experiments, the assays’ readings were comparatively analysed for the predefined numbers of U251 cells that were seeded and grown on the conventional tissue culture plastic or mixed with and embedded in Matrigel droplets. In dynamic assays, equal numbers of cells were mixed with Matrigel or seeded on the top of brain dECM scaffolds and cultured for 1, 3, 7, 14, or 21 days. During this time, cells seeded on brain dECM scaffolds gradually colonised the volume of the scaffolds, forming 3D tissue engineering constructs (TECs). The readings of the tested assays were compared at the same time points. We compared the performance of different assays within different cell culture platforms and conditions, as well as the performance of the same assays in different platforms, under static and dynamic conditions. The measures of the assays’ performance are explained in Section 2.2.

**Figure 1.**
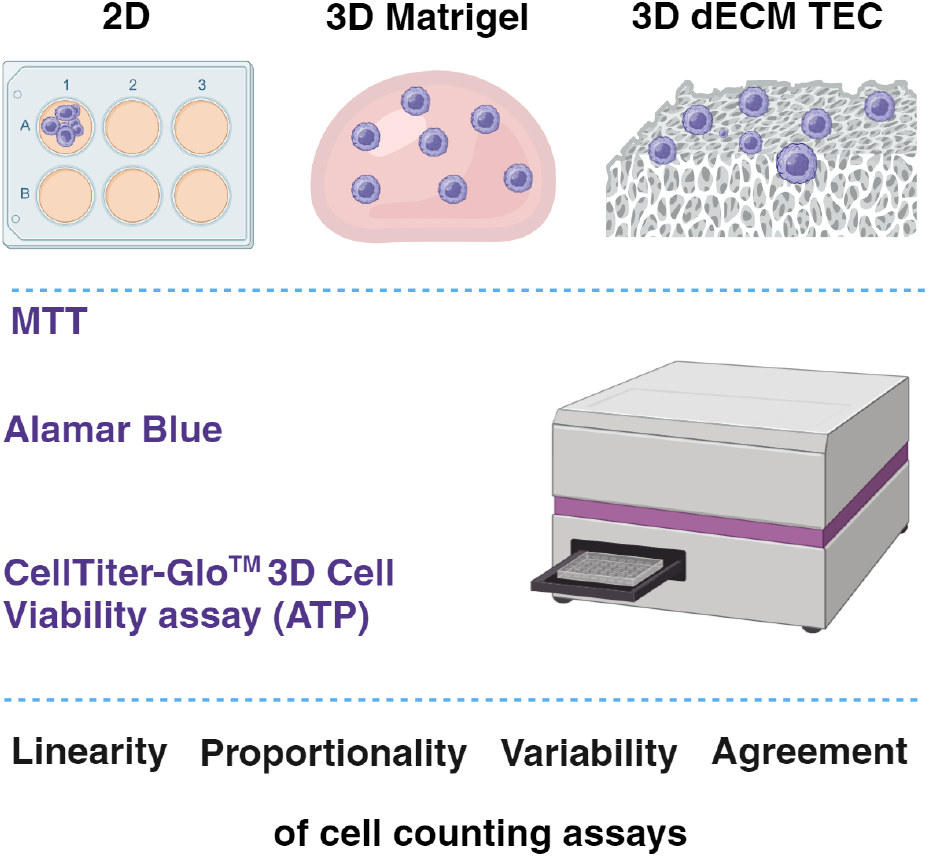
The schematic view of the current study design. Abbreviations: 2D – two-dimensional in vitro culture (monolayer); 3D – three-dimensional in vitro culture; TEC – tissue engineering construct. Image created with BioRender.com.

### 2.2. Quantification of the assays’ performance

In order to quantitatively evaluate the performance of different cell counting assays in 2D and 3D in vitro models of glioblastoma, we adapted several approaches suggested by Huang et al., who have demonstrated a practical implementation of recent ISO standards for cell counting and examined two cytometry instruments and two fluorescence staining methods in the suspension cultures ^[14]^. The cited study provides a detailed description of the calculation methods, advantages, and limitations of each used parameter of the assay quality.

Briefly, in the current study, we employed and adapted the following measures of the assays’ performance such as: 1) the measures of linearity; 2) the measures of variation, including precision, technical repeatability, and biological reproducibility; 3) the proportionality / correlation; and 4) the agreement between assays (by Bland-Altman plots).

#### 2.2.1. Measures of the linearity of the assays (R2 and aR2)

To assess the linearity of our assays, we analyzed the data using the linear regression algorithm of SPSS 29.0.0 software (IBM, Armonk, NY, USA). Specifically, we focused on the coefficient of determination (R^2^) and the adjusted R^2^ (aR^2^) as measures of how well our assays’ outcomes could be predicted from the independent variables, such as predetermined cell numbers or time in culture. The closer these coefficients are to 1, the more accurately our linear model reflects the experimental data, indicating strong assay linearity. Conversely, coefficients significantly lower than 1 would indicate poor linearity, showing that the model does not adequately explain the variance in cell counts.

#### 2.2.2. The proportionality index

The proportionality index (PI) measures how well a cell counting method adheres to the proportionality principle along the tested range, which expects that cell counts should steadily increase or decrease in direct relation to the changes of the independent variable (such as the pre-determined cell numbers in the current study). Deviations from proportionality suggest the presence of systematic or random errors, leading to reduced accuracy in the measurements.

In the current study, a PI shows the ratio of the fold changes observed through the assays’ readings to the fold changes of the cell numbers predetermined by the direct cell counting using Countess® II FL Automated Cell Counter (Thermo Fisher Scientific). The PIs were obtained for the experiments with 2D and static (one-day) Matrigel-based 3D cultures of U251 cells.

As in the longitudinal (dynamic) experiments with the 3D Matrigel constructs and TECs the exact cell number contained within these cultures was not known after seeding, the PIs were not employed for these experiments.

#### 2.2.3. Measures of the assays’ variation

The variation of the assay readings vs the independent variable characterizes the *<u>precision</u>* of the given assay. The variation was measured by the overall coefficient of variation (CV). The higher the variation (and the CV), the lower the precision is. The overall CVs were calculated as ratios of the standard deviations and means for the assay’s readings obtained across all the biological and technical replicates for the same predefined cell number (in static experiments) or days in vitro (in dynamic experiments) and multiplied by 100. So, the units of CVs are percentages, reflecting the deviation of the assay’s readings around their mean value.

The precision of the assay can be affected by the variation across the technical replicates (*<u>the technical repeatability</u>*) tested on the same day and the variation across the biological replicates of the same sample (*<u>the biological reproducibility</u>*), that are usually examined on different days and/or represent different passages of the cells.

Technical coefficients of variation (CVt), which characterize technical repeatability, were calculated as the ratios of the standard deviations to the means across technical replicates within each biological replicate. These were then averaged across the technical replicates of the same sample and multiplied by 100 to present the value as a percentage.

Biological coefficients of variation (CVb), indicators of biological reproducibility, were determined as the ratios of the standard deviations to the means between biological replicates of the sample, converted to a percentage value as previously described. Both CVt and CVb are measures of variation; therefore, their higher values indicate lower repeatability and reproducibility of the assay, respectively.

### 2.2.4. Estimation of the agreement between two assays by the Bland-Altman plots

To estimate the agreement between the assays, we used the Bland-Altman plots ^[18]^. The Bland-Altman plots are scatter plots where the mean of the measurements (in this study, it is the estimated cell number) from the two assays is shown on X-axis, and the difference between these measurements is reflected in axis Y. Additionally, three reference lines are depicted on these plots. The first line denotes the mean difference between the assays (average bias), while the other two lines delineate the upper and lower limits of agreement (LoA). These limits are determined by calculating the mean difference ± 1.96 × the standard deviation of the differences, thereby estimating the range within which 95% of the differences between the two methods are expected to fall. This facilitates a visual assessment of the assays’ agreement.

In interpreting Bland-Altman plots, good agreement between the two assays is suggested if most data points reside within the LoA and are evenly distributed across the measurement range. Conversely, a significant number of points outside these limits, or a discernible pattern in the distribution of points (e.g., a trend or funnel shape), may indicate systematic bias or a variability that changes with the measurement’s magnitude. This suggests that the assays might not agree throughout their entire measurement range.

### 2.3. Cell Line

Human glioblastoma U251 MG (formerly known as U-373 MG) was supplied by the European Collection of Cell Cultures (ECACC; Salisbury, United Kingdom) under catalogue number #09063001, and was purchased from CellBank Australia (Westmead, NSW, Australia). Cells were cultured at 37°C with 5% CO_2_ in a humidified atmosphere in a complete cell culture media, containing Dulbecco’s modified eagle’s medium (DMEM) (Gibco) supplemented with 1% (v/v) penicillin-streptomycin (PS) and 10% (v/v) fetal bovine serum (FBS). Cells were routinely passaged at 2 to 3 days intervals. This cell line was selected following the preliminary experiments that showed its efficient attachment to all used cell culture substrates, which was not affected by multiple washings required for some cell counting protocols. This feature was essential for the reliable comparative quantitative assessment of the tested assays.

### 2.4. 2D cell culture

For 2D culture platform, U251 cells at passages’ number 10-15 were stained with Trypan Blue and counted with Countess® II FL Automated Cell Counter (Thermo Fisher Scientific), by a standard procedure and seeded at 0.5 × 10^4^, 1.5 × 10^4^, 3.1 × 10^4^, 6.2 × 10^4^, 12.5 × 10^4^, 1.8 × 10^5^, 2.5 × 10^5^, 3.75 × 10^5^, and 5 × 10^5^ cells per well in 6-well-plates (9.5 cm^2^; Corning®, Lifesciences, United States). The examined cell counting assays were conducted in 12-15 hours after plating the cells, after cell attachment was confirmed. For each tested experimental condition, we examined at least three biological replicates with three technical replicates.

#### 2.5. 3D Matrigel constructs

#### 2.5.1. Matrigel preparation

Phenol-red free high protein concentration Matrigel (Corning® Matrigel®) was used for preparation of the 3D dome-shaped 3D cell cultures of U251 cells. Matrigel was removed from the freezer (−20 °C) and thawed overnight on ice in the fridge. We used pre-chilled pipette tips to prevent premature gelation of Matrigel. High protein concentration Matrigel (21 mg/ml) was diluted with ice-cold phenol red free and serum-free DMEM to a final concentration of 6 mg/mL.

#### 2.5.2. Static 3D Matrigel-based cultures

The static Matrigel-based 3D culture experiments were performed to examine the performance of different assays under conditions when the predetermined (cell counter measured) number of cells were embedded in the Matrigel plugs or the Matrigel plugs were left cell-free. The assays were performed without using a tissue-dissociating solution for cell recovery, in a few hours after the preparation of the Matrigel constructs, to avoid the variation associated with proliferation of the cells. The aim of these experiments was to determine the effect of the Matrigel matrix on the assays’ readings.

The cells were counted using Countess® II FL Automated Cell Counter (Thermo Fisher Scientific) and resuspended in 50 µL of cold Matrigel (6 mg/mL) with cell densities of 0.5 × 10^4^, 1.5 × 10^4^, 3.1 × 10^4^, 6.2 × 10^4^, 12.5 × 10^4^, 1.8 × 10^5^, 2.5 × 10^5^, 3.75 × 10^5^ and 5 × 10^5^ cells per 50 µL droplet. The droplets were deposited onto a pre-incubated cover slips placed individually in the wells of 24-well plate and allowed to polymerise at 37°C. After one hour, 1 mL of DMEM was added to each well. Then, the resulting 3D Matrigel constructs were incubated at 37°C for 12-15 hours before the assays’ performance was tested. At least three biological replicates with three technical replicates was used for each tested experimental condition.

#### 2.5.3. Dynamic 3D Matrigel-based cultures

In the following dynamic experiments, the Matrigel-based 3D cultures of U251 cells were seeded with 10,000 cells (enumerated using Countess® II FL Automated Cell Counter (Thermo Fisher Scientific)) and maintained in vitro for up to 21 days. Consequently, in this series, the direct measurement of cell numbers was only available on the day of cell seeding, while afterwards it was estimated via the assay’s readings normalized as fold changes against the average reading obtained on the first day of culture and corresponding to 10,000 cells.

The cells were counted using Countess® II FL Automated Cell Counter (Thermo Fisher Scientific) and subsequently, they were resuspended in cold Matrigel (6 mg/mL) to achieve a final concentration of 200,000 cells/mL. Following this, 50 µL droplets of Matrigel-based constructs containing 10,000 cells each were dispensed onto pre-incubated glass cover slips, placed within individual wells of a 24-well plate to prevent cells’ attachment to the plastic, and allowed to solidify at 37°C. Following a one-hour incubation, 1 mL of complete cell culture medium was introduced into each well. The Matrigel constructs were then cultured at 37°C in vitro for 21 days. The culture medium was replaced with the fresh one every 2 days. Assays were performed at different time points during the culture on 1, 3, 7, 10, 14, 17 and 21^st^ days of in vitro culture. Each experimental condition was replicated at least three times biologically, with each biological replicate having three technical replicates.

### 2.6. 3D tissue engineering constructs (TECs)

Glioblastoma 3D tissue engineering constructs (TECs) were obtained by seeding and culturing of U251 cells on decellularized sheep brain scaffolds. First, the fresh sheep (lamb) brain tissue was decellularized (DCL) as described by us elsewhere ^[19]^ with minor modifications ^[15]^. A detailed protocol of the DCL and associated procedures is presented in Section A1 (Appendix A). Next, the fragments of the decellularized brain tissue were sterilized chemically, cut into smaller pieces and again sterilized by UV light and pre-conditioned in a complete cell culture media as described in Section A2 (Appendix A).

To form TECs, the media from the wells containing sterilized and pre-conditioned scaffolds was removed, and the scaffolds were left to semi-dry for 20 minutes for the better attachment of the cells from suspension. A 20 μl drop of U251 cells’ suspension containing (5 × 10^6^ cells/mL, or 100,000 viable cells per scaffold) counted using Countess® II FL Automated Cell Counter (Thermo Fisher Scientific) was pipetted onto the centre of each scaffold. The cells were allowed to settle and attach to the scaffolds for 2 h at 37°C. Next, 2 mL of complete cell culture media was added slowly to each well. After overnight incubation, the obtained 3D TECs were aseptically transferred to the new 24-well plates in order to avoid the outgrowth of the non-scaffold bound cells on the plastic bottoms of the wells. The 3D cultures of U251 cells in TECs were grown at 37 °C under a humidified atmosphere of 5% CO_2._ The cell culture media was changed every 2 days.

The U251 3D TECs were cultured in vitro for three weeks. The number of viable cells was measured at various time points, including days 3, 7, 10, 14, 17, and 21 after seeding on scaffolds, using DNA, MTT, Alamar Blue, and ATP quantification assays (see Section 2.6.1 – 2.6.4 for details). At each time point, TECs were transferred into new multiwell plates prior starting of the cell counting assays’ reagents to ensure that the signal being measured was from cells attached and growing on the scaffold and not from the cells that had attached to the surface or sides of the culture wells. Repeated measures (three biological replicates with three technical replicates in each of them, N = 9) were used for each time point.

In preliminary experiments, different types of cells, including neuron-like PC12 cells of rat origin, human cerebral microvascular endothelial cells HCMEC-SV40, human and mouse microglial cells HMC3 and BV2, mouse macrophages RAW), as well as human glioblastoma U87 and U251 cells, were successfully seeded and cultured on the sheep brain dECM scaffolds. However, for the purposes of the current study, we selected the U251 cell line for a simplified 3D TEC model of glioblastoma. In particular, in contrast to U87 cell line, U251 cells do not form spheroid-like cell aggregates at high cell densities in 2D culture. Such aggregates are easily washed out during cell counting assays, which may significantly affect the quality of the data.

The scaffolds and TECs were morphologically characterized using histology, immunohistochemistry and scanning electron microscopy as described in Section A3 (Appendix A). The results of the morphological characterization of the scaffolds and 3D TECs are presented in Section A4 (Appendix A).

### 2.7. Cell counting and viability measurements

#### 2.7.2. MTT assay

MTT (#M5655, Sigma Aldrich) was prepared by dissolving 0.5 mg/mL MTT in phenol red-free DMEM/F12 (#D6434, Sigma-Aldrich, North Ryde, NSW, Australia), with 1% (v/v) PS and 10% (v/v) FBS and filtered through a 0.22 μm filter. Cell culture medium was removed from the wells with the tested cultures and 1 mL of the stock MTT solution was added to the wells. Then, the plates were incubated at 37°C for 2 hours. After the incubation, MTT reagent was removed, and the formazan crystals were dissolved in 1 mL DMSO per well.

Triplicate aliquots of the resulting solutions were transferred in 96-well plates, and the light absorbance was recorded at 560 nm using a PHERAstar FS plate reader (BMG Labtech).

#### 2.7.3. Alamar blue assay

Culture medium was replaced with 1 mL of phenol red-free DMEM/F12 (#D6434, Sigma-Aldrich, North Ryde, NSW, Australia), with 1% (v/v) PS and 10% (v/v) FBS, containing 0.1 mg/mL of reassuring sodium salt (#199303, Sigma-Aldrich, North Ryde, NSW, Australia) filtered through a 0.22 μm filter. After 2 h of incubation, the fluorescence of triplicate samples (100 µL) from each well measured in a black-walled 96-well plates, at the respective excitation and emission wavelengths of 540 and 620 nm, respectively, using PHERAstar FS plate reader (BMG Labtech).

For each well, the fluorescence readings were done in triplicate (values obtained from 3 different wells were averaged). For each experiment, wells containing only the Alamar Blue solution without cells were also prepared and incubated for 2 h. The fluorescence measured in these samples was used as a background and subtracted from the readings obtained in cell-containing samples.

#### 2.7.4. CellTiter-Glo® 3D Cell Viability Assay (ATP assay)

ATP was quantified using the CellTiter-Glo® luminescent 3D Cell Viability Assay (Promega), according to the manufacturer’s protocol. Briefly, the cell culture medium in 2D cultures and Matrigel-based 3D constructs was replaced with 250 µL of phenol red-free Dulbecco’s modified Eagle’s medium (Sigma), and an equal volume of the assay reagent was added to each well and mixed well. The same procedure was applied to the 3D TECs, at each time point. The cell cultures were incubated for 30 minutes with the assay mixture at room temperature. After that, luminescence was quantified using a PHERAstar FS plate reader. The reading obtained from the samples were compared to a known concentrations of ATP standards within the 96-well microtiter plate format.

### 2.9. Statistical analysis

Data analysis was carried out using several the standard software packages, including Microsoft® Excel® for Microsoft 365 MSO (Version 2310 Build 16.0.16924.20054), SPSS 29.0.0. (241) (IBM Statistics), and GraphPad Prism version 9.00 for Windows (GraphPad Software, Inc.). For the descriptive purposes, the data is presented as mean ± standard deviation if not overwise specified. The level of statistical significance *p* was accepted at the value <0.05.

The deviation of the quantitative data from normal distribution was checked with one sample Kolmogorov-Smirnov test and Shapiro–Wilk’s test. The intergroup differences for the normally distributed data were evaluated using a one-way ANOVA with Tukey’s post-hoc test. For the data deviated from the normal distribution or if the ANOVA or other assumptions of the parametric approaches were not met, which was the most common in the current study, we applied nonparametric methods of statistical analysis.

For the intergroup comparisons, Kruskal-Wallis’s test was employed as a nonparametric analogue of ANOVA for three or more independent samples. The Mann-Whitney test served as an alternative to the independent samples t-test. For two related samples, we used Wilcoxon signed-ranks test as a nonparametric alternative to a paired-samples t-test. The regression analysis, curve fitting procedures and bivariate Spearman’s correlations were applied to examine the relationship between the assays’ readings and cell numbers or the time of the dynamic 3D cultures growth. The bivariate correlation coefficients were calculated together with the asymptotic (Fisher) confidence intervals for the cross-comparison of the performance of the various studied assays in different cell culture conditions.

### 2.10. Application of artificial intelligence tools

In the preliminary stages of this research, ChatGPT-4, developed by OpenAI, was utilized for initial brainstorming for the possible approaches for the comparative analysis of the performance of various cell viability assays. In the preparation of this manuscript, Grammarly (Grammarly Inc., Ukraine, USA) was employed for grammar, spelling, and plagiarism checks to ensure the clarity, accuracy, and originality of the written content. The use of both ChatGPT-4 and Grammarly contributed to refining the final text, complementing our manual editing processes, and aligning with our commitment to high academic standards.

## 3. RESULTS

### 3.1. Estimation of cell number by cell counting assays in static (1-day) 2D and Matrigel-based 3D cell cultures of U251 cells

#### 3.1.1. Overview of the results of all examined assays

The overview of the results of the cell number estimation by different assays in static 2D and 3D cultures of U251 cells is presented in Figure 2. As the full range (100,000 – 2,000,000 cells) of predefined cell numbers examined using the DNA assay in 2D culture was wider than the range for the metabolic assays (5,200 – 500,000 cells), it is displayed separately in Appendix B (Figure B1 (b)). To make the comparisons between the assays more accessible, we employed the following procedure of normalization. The estimated cell number (ECN) for each data point was calculated based on the fold change (FC) in assay readings relative to a reference sample. For each assay, the FC was determined by dividing the assay reading of a given sample by the assay reading obtained for the reference sample with the smallest number of cells. Specifically, the reference sample had 100,000 cells for the DNA assay and 5,200 cells for the Alamar Blue, MTT, and ATP assays. The resulting FC was then multiplied by the known cell number of the reference sample to estimate the cell number for each respective sample. While all assays revealed an increase in ECNs along with increasing directly counted (predefined) cell numbers, there were notable differences in their performance and precision across 2D and 3D (Matrigel-based) cultures.

**Figure 2.**
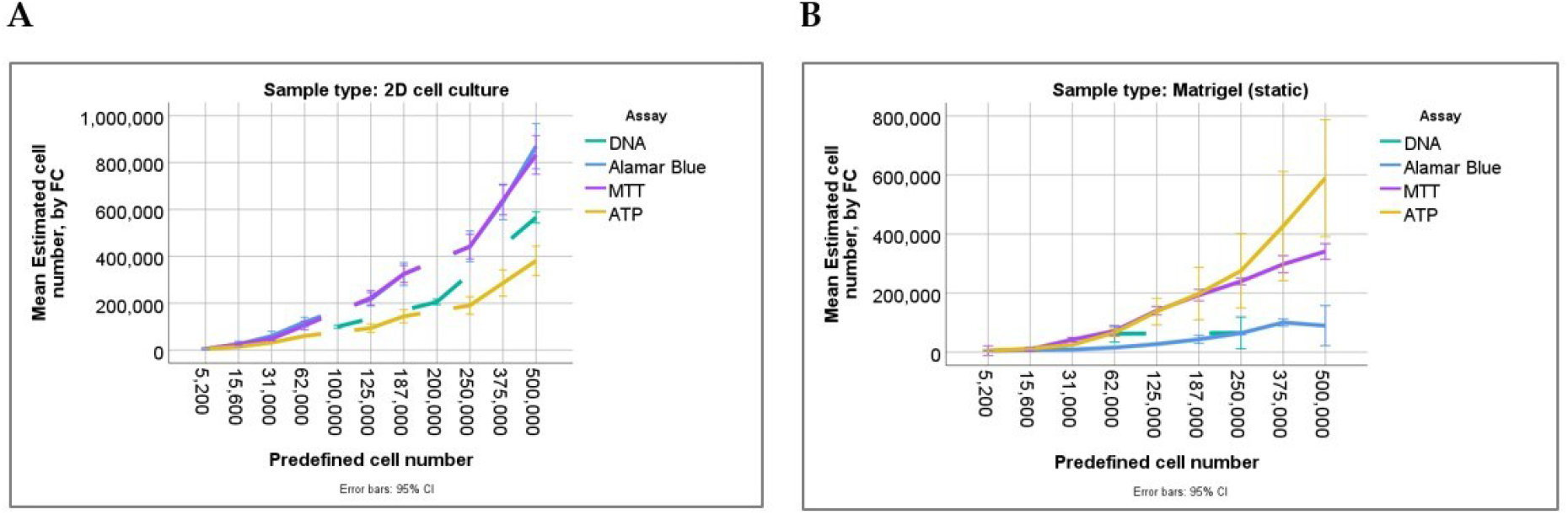
Overview of the relationship between the cell numbers estimated by different assays and the cell numbers counted directly (predefined) in static (1-day) experiments in 2D and 3D Matrigel-based in vitro cultures of U251 cells. (A) 2D cell cultures: note the similarity of different assays’ results in the lower cell density samples (with the directly counted cell number between 5,200 and 31,000). At the higher cell densities, the relatively higher cell numbers were estimated by Alamar Blue and MTT assays and relatively lower ones by ATP assay, compared with DNA assay, which mostly shows the close similarity between the estimated and predefined cell numbers. (B) 3D Matrigel-based constructs: note the limited range of DNA assay and minimal change of the assay readings in the tested range of cell numbers. The results of the metabolic assays are similar at the lower cell densities and differ from each other at higher predefined cell numbers. Alamar Blue and MTT assays tend to underestimate cell numbers in higher density 3D Matrigel-based cultures, while ATP demonstrates some overestimation, compared with the predefined cell number in the sample.

The detailed results of the cell counting assays in static 2D and 3D cultures of U251 cells are presented below.

#### 3.1.3. Alamar Blue, MTT and ATP assays

The overview of the results of Alamar Blue, MTT and ATP assays in static 2D and 3D cultures of U251 cells is presented in Table 3. The ECN was calculated by multiplying the FC in assay readings—using the smallest sample as the reference, which contained 5.2×10^3^ cells—by the number of cells in that reference sample. The values of the FCs of the assays’ readings are presented in Table B2 (Appendix B) with a comment following it.

**Table 3.** Cell number estimation by the Alamar Blue, ATP and MTT assays in static (1-day) 2D and 3D Matrigel-based cell cultures of U251 cells.

| Number of cells,<br>counted directly, $\times 10^3$ | ECN*, $\times 10^3$ | | | | | |
| --- | --- | --- | --- | --- | --- | --- |
|  | Alamar Blue |  | MTT |  | ATP |  |
|  | 2D | 3D | 2D | 3D | 2D | 3D |
| 5.20 | 5.20±2.23 | 5.20±993 | 5.20±1.60 | 5.20±6.29 | 5.20±310 | 5.20±2.97 |
| 15.60 | 24.97±11.75 | 7.54±3.17 | 25.07±6.57 | 9.53±6.74 | 14.39±980 | 11.61±1.89 |
| 31.00 | 60.00±19.04 | 8.72±2.11 | 48.02±12.24 | 42.26±6.80 | 31.98±966 | 24.93±557 |
| 62.00 | 119.61±19.65 | 15.33±2.92 | 104.96±28.98 | 71.86±14.36 | 60.50±1.96 | 65.79±10.32 |
| 125.00 | 216.71±27.01 | 27.22±3.40 | 222.74±48.97 | 140.44±13.63 | 93.76±28.77 | 137.27±36.15 |
| 187.00 | 324.20±46.19 | 43.17±13.08 | 324.97±57.38 | 193.02±18.46 | 144.23±43.99 | 198.23±71.36 |
| 250.00 | 442.50±62.27 | 64.36±5.35 | 441.36±82.65 | 239.70±10.75 | 190.70±58.26 | 275.43±101.34 |
| 375.00 | 631.97±71.85 | 100.34±11.38 | 640.36±100.02 | 297.69±27.33 | 285.99±87.80 | 426.57±148.92 |
| 500.00 | 869.49±91.68 | 89.68±64.43 | 832.52±129.38 | 341.01±24.98 | 381.01±98.52 | 589.26±159.50 |

\* The ECN was calculated by multiplying the fold change of the assay readings relative to the sample with the smallest cell count ( $5.2 \times 10^3$ ) by this baseline number of cells ( $5.2 \times 10^3$ ). Abbreviations: ECN – estimated cell number; 2D – two-dimensional cell culture; 3D - three-dimensional Matrigel-based constructs. The data presented as Mean ± St. Dev.

As shown in Table 3, in 2D cell culture conditions, the Alamar Blue, MTT, and ATP assays all detected increases in estimated cell numbers (ECNs) in parallel with the rise in directly counted cell numbers. The ECNs reported by Alamar Blue ranged from 5.20×10^3^ to 869.49×10^3^ cells. The MTT assay’s ECNs showed a progression from 5.20×10^3^ to 832.52×10^3^ cells, while the ATP assay’s ECNs started at 5.20×10^3^ and extended up to 381.01×10^3^ cells, compared to the variation of the directly counted cell number between 5.20×10^3^ and 500×10^3^ cells. There was notable similarity between the ECNs obtained by the Alamar Blue and MTT assays in 2D cultures, both overestimating the cell contents vs the predefined cell number, starting in a low cell density (15,600 -31,000 cells) and continuing till the end of the tested range. At the same time, the ATP assay’s results were very close to the predefined cell numbers in the range between 5,200 and 62,000 cells, and after that they became rather underestimating, compared to the directly counted cell numbers.

In 3D Matrigel-based cell cultures, the Alamar Blue, MTT, and ATP assays similarly exhibited increases in ECNs as the directly counted cell numbers escalated. For Alamar Blue, the ECNs varied from 5.20×10^3^ to 89.68×10^3^ cells at the highest. The MTT assay showed ECNs beginning at 5.20×10^3^ and reaching up to 341.01×10^3^ cells. The ATP assay’s ECNs also rose with increasing cell counts, starting from 5.20×10^3^ and peaking at 589.26×10^3^. Considering the changes of ECNs in static 3D Matrigel-based culture, the Alamar Blue assay demonstrated severe underestimation of the cell count vs the predefined cell numbers, while the ATP assay’s results provided a slightly overestimated values of the cell densities. The MTT assay showed the intermediate outcomes, with a moderate underestimation of the cell counts in 3D Matrigel constructs cultured for 1 day.

The ECNs obtained in static 3D cultures using the metabolic assays demonstrated the reversal of over- and under-estimation trends, compared with the 2D cultures. In particular, the Alamar Blue and MTT assays that overestimated the cell numbers in 2D conditions, were underestimating in 3D. At the same time, the ATP assay that underestimated cell contents in 2D cultures showed some overestimation in 3D static Matrigel-based cultures. Overall, Alamar Blue and MTT assays showed lower ECNs in 3D vs 2D, while the ATP assay showed lower ECNs in 3D vs 2D culture are low cell densities and higher ECNs, respectively, at the higher cell densities (≥62,000 cells).

To examine whether these deviations of ECN vs the predefined cell numbers are statistically significant, we employed the non-parametric Wilcoxon signed ranks test. It was found that all metabolic assays in static 2D cell culture provide ECNs that were statistically significantly different from the cell numbers counted directly with (p<0.001 for all the assays). In 3D Matrigel-based cultures, the Alamar Blue assay had statistically significant deviation of the ECN from the predefined cell numbers (mostly underestimation), with p<0.001. Interestingly, there was no statistical significance of the differences between the ECNs obtained using MTT and ATP assays and the predetermined cell numbers (p=0.120 and p=0.433, respectively) in static Matrigel constructs.

The Mann-Whitney U-test results for estimated cell numbers (ECNs) obtained by FC in static 2D versus 3D Matrigel-based cell cultures reveal distinct patterns of the metabolic assays’ responsiveness to the cell culture environment. For the Alamar Blue assay, a significant difference in ECNs between 2D and 3D cultures was observed (p < 0.001), with ECNs being notably higher in 2D cultures. The MTT assay also demonstrated a significant disparity in ECNs between the two culture conditions (p = 0.014). Contrastingly, the ATP assay showed no statistically significant difference in ECNs between 2D and 3D cultures (p = 0.442).

#### 3.1.4. Quantification of the cell counting assays’ performance in static (1-day) 2D and 3D Matrigel-based cultures of U251 cells

##### Analysis of the assays’ linearity (linear regression models)

To examine the linearity of the assays, we applied linear regression models. Table 4 summarizes the results of fitting the ECNs obtained by the metabolic assays in 2D culture of U251 cells (see Table 3) to the linear regression curves, with using the directly counted cell number as an independent variable. The R value reflects the Pearson’s correlations coefficient (R) between the ECN and the directly counted cell number. The R^2^ and aR^2^ are the coefficients of determination of the model and the main measures of the assays’ linearity (the closer their values to 1, the more linear the model is). The standard error of estimation suggests the average distance of the data points from the fitted line. The intercept shows the expected ECN for the samples containing zero cells. It is interpreted as the background signal as well as an indication of the assay’s sensitivity threshold above which the cell counting becomes reliable if the model is statistically significant. The coefficient for the predefined cell number, B, shows the slope of the model curve. Finally, the statistical significance of the model is presented via p-values. The visual representation of the linear regression models fitting to the experimental data is shown in Figure B2 (Appendix B). The outcomes of regression analysis are4 presented in Table 4.

**Table 4.** Summary of linear regression model outcomes for the estimated cell number via FC according to the metabolic assays’ readings in static 2D and 3D (Matrigel-based) cultures of U251 cells.

| Assay and sample type | R Value | R<sup>2</sup> | aR<sup>2</sup> | Std. Error of the Estimate | Intercept | B* | Sign. (p) |
| --- | --- | --- | --- | --- | --- | --- | --- |
| Alamar Blue (2D) | 0.987 | 0.973 | 0.973 | 47,223.05848 | 3,959.485 | 1.717 | <0.001 |
| Alamar Blue (3D) | 0.825 | 0.680 | 0.674 | 23,262.24399 | 5,225.376 | 0.203 | <0.001 |
| MTT (2D) | 0.973 | 0.947 | 0.946 | 66,242.59862 | 3,315.865 | 1.688 | <0.001 |
| MTT (3D) | 0.967 | 0.934 | 0.933 | 30,757.62155 | 30,596.412 | 0.696 | <0.001 |
| ATP (2D) | 0.928 | 0.862 | 0.860 | 49,628.22733 | 5,090.825 | 0.749 | <0.001 |
| ATP (3D) | 0.930 | 0.865 | 0.862 | 77,747.47436 | -9,804.739 | 1.175 | <0.001 |

\*B is the Coefficient (B) for the Predefined Cell Number.

Table 4 reveals distinct patterns in assays’ linearity across static 2D and 3D environments. Within 2D cultures, all the assays exhibit strong predictive capabilities in linear manner: DNA (R^2^ = 0.991), Alamar Blue (R^2^ = 0.973), MTT (R^2^ = 0.947), and ATP (R^2^ = 0.862) combined with high values for the slope. The intercepts are positive, suggesting the baseline levels of detection above zero and reflecting either inherent assay sensitivity or background signals. In 3D Matrigel-based static cultures, the linearity of the DNA, Alamar Blue and ATP assays is reduced, compared to the monolayer conditions. The DNA assay’s predictive power plummets (R^2^ = 0.000, aR^2^ = -0.033; p=0.912), suggesting its limited utility in 3D due to negligible linearity. The slope near zero (0.017) combined with a high intercept (60,952.537) reflects the high background noise. The Alamar Blue linearity is the lowest one among the metabolic assays in 3D Matrigel cultures (R^2^ = 0.680) and the slope value is also low, while the model remains statistically significant. The ATP assay maintain substantial linearity in 3D culture (R^2^ = 0.865), albeit with some adjustments from its 2D performance. Notably, the ATP assay demonstrates remarkable consistency, with its R^2^ value slightly improving in 3D (aR^2^ = 0.860 to 0.862), highlighted by a significant slope (1.175) that suggests a robust correlation even in complex 3D matrices. However, the negative intercept of the ATP assay (-9,804.739) in 3D Matrigel cultures is intriguing, potentially indicating initial underestimation of cell counts at lower cell densities before the assay readings align more closely with actual cell numbers. Finally, the MTT assay in the static Matrigel cultures shows high R^2^ value (0.934), with intermediate slope (0.696) and high statistical significance of the model. However, the intercept value for this assay in 3D culture is roughly ten times higher than in 2D environments, indicating that the assay becomes efficient in detecting of the changes in cell counts at the level above >30,000 cells per sample.

##### Analysis of the assay’s proportionality

Figure 3 presents the distribution of proportionality indexes (PIs) for various assays in 2D and 3D (Matrigel-based) static cell cultures. The DNA assay in 2D culture closely aligns with the expected cell count (PI = 1.02±0.06), suggesting a highly proportional response. In contrast, Alamar Blue (PI = 1.73±0.11) and MTT (PI = 1.61±0.24) assays show elevated PIs, reflecting a tendency to overestimate cell numbers. The ATP assay (PI = 0.86±0.12) typically yields lower ECNs than the predefined cell numbers. In the 3D context, the Alamar Blue and DNA assays’ PIs reveal a substantial deviation from the expected proportionality and drop sharply (PI = 0.35±0.26 and PI = 0.77±0.33, respectively). The MTT assay maintains a closer-to-ideal PI (0.97±0.24). The ATP assay, surprisingly, gets closer to the ideal range, compared to the value obtained in 2D culture and demonstrates just a slight overestimation trend (PI = 1.02±0.15).

**Figure 3.**
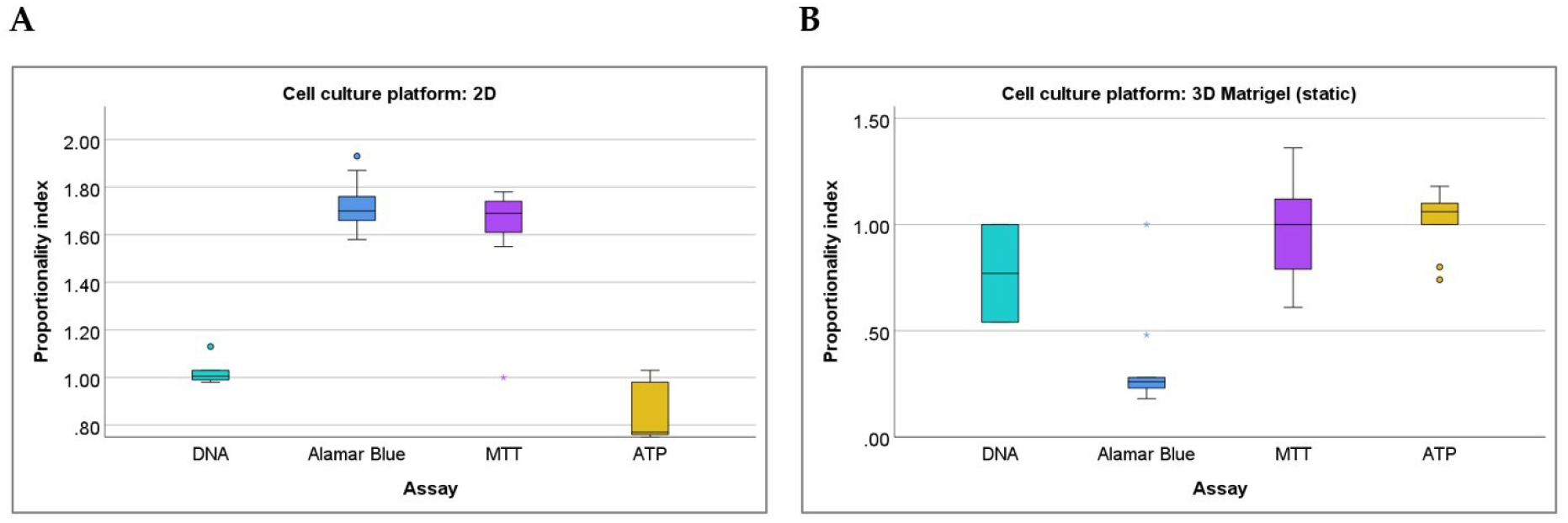
The proportionality indexes obtained for different cell counting assays in static (1-day) 2D and 3D (Matrigel-based) cell cultures of U251 cells. **(A)** 2D culture; **(B)** 3D Matrigel-based constructs.

Statistical analysis using the Kruskal-Wallis Test detected pronounced differences in the distribution of PIs among assays within each culture condition (Table B3 in Appendix B), indicating variability in their relative estimation capacity. In 2D cultures, the statistically significant contrasts are notable between ATP and MTT (p < 0.001) assays, ATP and Alamar Blue (p < 0.001) assays and ATP and DNA (p=0.044) assays. At the same time, there are no statistically significant differences in the PI values between the ATP and DNA assays, DNA and MTT assays, and Alamar Blue and MTT assays. In the 3D setting, significant variations are observed, particularly for the Alamar Blue assay when compared to MTT (p = 0.002) and ATP (p < 0.001) assays.

Next, we also examined the Spearman’s correlation coefficients for the PI values and the predefined cell numbers. It was found that in static 2D cultures, the PIs for the DNA, Alamar Blue and MTT assays do not show statistically significant associations with the cell numbers. This indicates that these assays remain proportional across the tested cell number ranges. In contrast, the PI of the ATP assay in 2D culture shows a statistically significant strong negative correlation with the directly counted cell number (Rs=-0.729, p=0.026, N=9), reflecting the increase of the cell number underestimation with the predefined cell number getting higher. In the static 3D culture, the statistically significant correlations with the predefined cell number were detected for the PIs of the Alamar Blue assay (Rs=-0.717, p=0.030, N=9) and the ATP assay (Rs=0.908, p<0.001, N=9), showing the increasing under- and severe overestimation of cell counts with the increase of the predefined cell numbers.

##### Analysis of the assays’ variation

To explore the technical replicability, biological reproducibility and the overall precision of the cell counting assays, we calculated the values of the technical, biological and overall coefficients of variation, CVt, CVb, and CV, respectively. The results are shown in Figure 4. Detailed information on the values of the CVt, CVb, and CV for different assays and cell culture conditions is reported Table B4 (in Appendix B).

**Figure 4.**
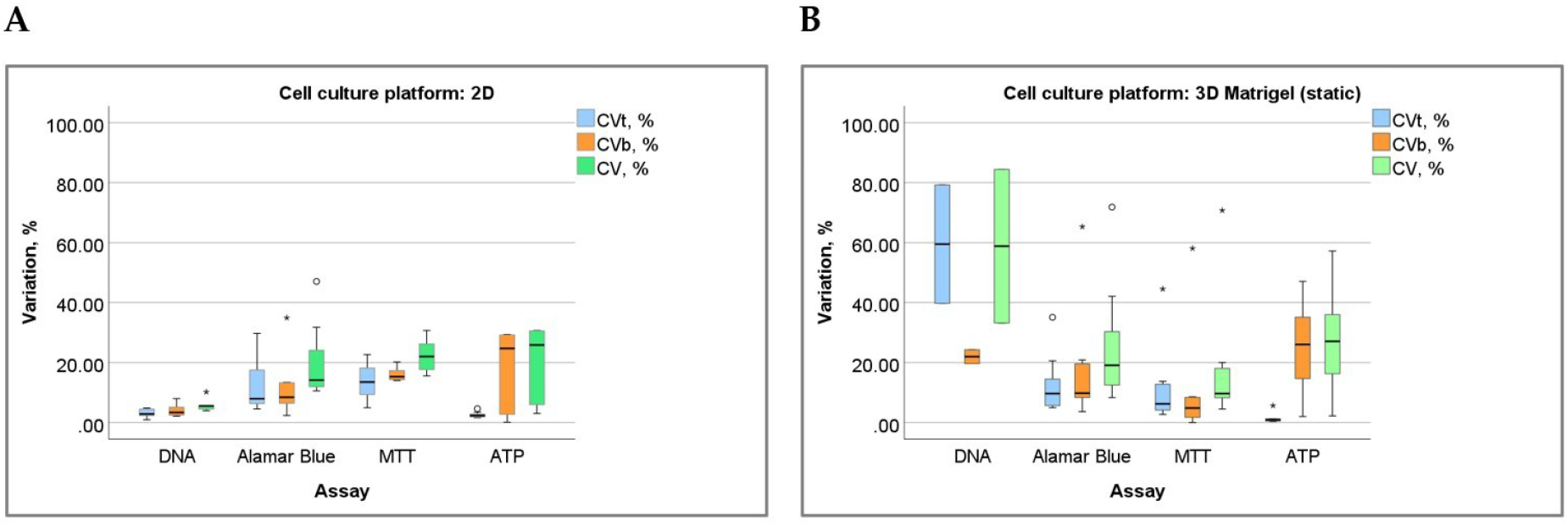
The coefficients of variation (CVt, CVb and CV) obtained for different cell counting assays in static (1-day) 2D and 3D (Matrigel-based) cell cultures of U251 cells. **(A)** 2D culture; **(B)** 3D Matrigel-based constructs.

The differences between the assay’s coefficients of variation within the same cell cultures platform (Tables B5, B6, and B7 in Appendix B) were examined using the Kruskal-Wallis Test with pairwise comparisons. It has revealed significant differences in technical variability (CVt) across the assays in both 2D and 3D (Matrigel-based) cultures (p < 0.001). Specifically, the ATP assay exhibited significantly lower CVt compared to Alamar Blue (p = 0.004) and MTT (p = 0.001). At the same time, there was no significant difference between the ATP and DNA assays (p = 1.000). In 3D Matrigel cultures, the ATP assay maintained lower CVt relative to DNA (p = 0.008), Alamar Blue (p = 0.007), and MTT (p = 0.010).

Statistically significant differences were also observed in biological variability (CVb) values between the DNA and MTT assays (p = 0.020) in 2D cultures (CVb for DNA assay was lower than that of MTT assay) and between the MTT and ATP assays (p = 0.038) in 3D Matrigel constructs, due to the lower CVb in the MTT assay than in the ATP test.

Overall variability (CV) also varied significantly across assays within 2D cultures (p = 0.014), revealing distinct total variability profiles for each assay. It was linked to the fact that DNA assay had CV value lower than CV of the Alamar Blue assay (p=0.035) and MTT (p=0.014) assays. However, no significant differences were observed in 3D cultures (p = 0.260).

Interestingly, across the cell culture platforms, we detected statistically significant differences only in three cases: for all coefficients of variation of the DNA assay (the vaues of CVt, CVb, and CV were lower in 2D vs 3D cultures, p=0.046); this is explainable by the very high noise in DNA assay in Matrigel; b) for CVb of the MTT assay (p=0.009, with a lower value in 3D vs 2D conditions); and c) for CVt of the ATP assay (p=0.04, with a lower value in 2D culture).

To determine whether the changes in the technical, biological and overall variation are associated with each other, or with the predefined cell number in the culture, we employed analysis of correlations (Spearman’s correlation coefficient) (details are provided in Table B9, Appendix B). A summary on the detected statistically significant links is show in Table 5.

**Table 5.** Summary of statistically significant non-parametrical correlations (Spearman’s coefficient, Rs) for the measures or variation in static 2D and 3D (Matrigel-based) cultures of U251 cells.

| Culture Plat-<br>form | Assay | Correlation with Cell Number |  | Correlation between<br>CV, CVt and CVb |  |
| --- | --- | --- | --- | --- | --- |
|  |  | Detected links | Rs | Detected<br>links | Rs |
|  | Alamar<br>Blue | CVt, CVb, CV ↓ with Cell<br>Number | CVt: -0.786, CVb: -0.786,<br>CV: -0.933 | CVt, CVb -><br>CV | CVt and CVb with<br>CV: 0.786 |
|  | MTT | CVt, CV ↓ with Cell<br>Number | CVt: -0.950, CV: -0.933 | CVt -> CV | CVt with CV: 0.967 |
|  | ATP | CVt ↓ with Cell<br>Number | CVt: -0.717 | CVb -> CV | CVb and CV: 1.000 |
|  | Alamar<br>Blue | CVt ↓ with Cell<br>Number | CVt: -0.733 | CVb -> CV | CVb and CV: 0.800 |
|  | MTT | CVt, CV ↓ with Cell<br>Number | CVt: -0.717, CV: -0.933 | CVt -> CV | CVt with CV: 0.850 |
|  | ATP | - | - | CVb -> CV | CVb and CV: 0.983 |

Table 5 demonstrated a series of interesting relationships. In particular, it reveals the potential influence of cell density (number of cells) on the assays’ variation. There changes of CV, CVt and CVb along the cell numbers’ ranges are ilustrated by Figure B3 (Appendix B). For example, in 2D cultures, all metabolic assays showed a decrease in CVt and overall CV as the cell number increased, indicating that higher cell densities led to more consistent results. In 3D Matrigel comstructs, the Alamar Blue and MTT assays showed a significant decrease in technical variation with an increase in cell number, while the MTT assay in parallel also demonstrated strong decrease in overall variation (CV). Surprisingly, there were no statistically significant correlations between the coefficients of variation and cell numbers for the ATP assay in 3D cultures.

Moreover, it becomes visible that the overall variation (CV) in many cases was specifically dependednt on technical or biological variation, or on both of them. Biological variation was statistically significantly positively correlated with CV for DNA, Alamar Blue, and ATP assays in 2D cultures and for Alamar Blue and ATP assays in 3D conditions. Technical variation positively correlated with the CV for the Alamar Blue in 2D culture and for the MTT assay in both 2D and 3D cultures.

##### Analysis of the inter-assay agreement (Bland-Altman plots)

As a next step, we examined the extent of the agreement in cell numbers estimation between the metabolic assays applied in the static 2D and 3D cultures of U251 cells using Bland-Altman plots (Figure 5). The comparisons with DNA assay were not considered due to the differences in cell seeding protocols (and the ranges of the tested cell numbers, respectively) between the DNA assay and the metabolic assays.

**Figure 5.**
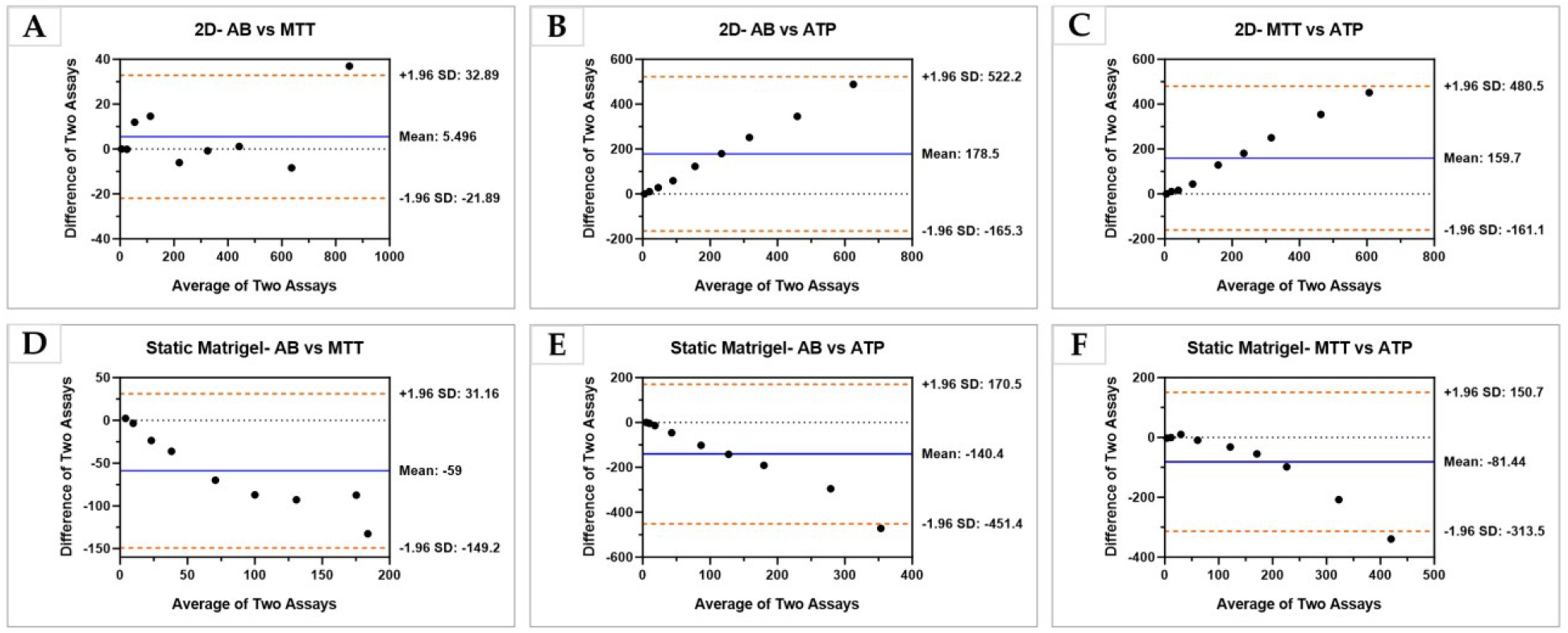
Assessment of the agreement between the metabolic assays regarding the estimated cell number in the static (1-day) 2D and 3D (Matrigel-based) cultures of U251 cells by Bland-Altman plots. The X-axis represents the average estimated cell numbers (ECN) (in thousands) obtained by different assays for the same number of the cells counted directly. The Y-axis shows the difference between the ECNs (in thousands) by the two assays under comparison. The “Mean” indicates the difference between the assays’ ECNs (in thousands) is depicted as a horizontal blue line. The dashed orange lines demonstrate the positions of the limits of agreement (LoA) that correspond to the boundaries of the CI95% for the mean difference. (A, B, C) Bland-Altman plots for the 2D static cell cultures and the following assay pairs: (A) Alamar Blue vs MTT assays; (B) Alamar Blue vs ATP assays; (C) MTT vs ATP assays. (D, E, F) Bland-Altman plots for the static 3D Matrigel-based cell cultures and the following assay pairs: (D) Alamar Blue and MTT assays; (E) Alamar Blue and ATP assays; (F) MTT and ATP assays.

Figure 5 shows the results of the assessment of the agreement in three pairs of assays (Alamar Blue and MTT, Alamar Blue and ATP, and MTT and ATP assays). In the 2D setup, Alamar Blue and MTT showed a mean discrepancy of 5.496 × 10^3^ cells with limits of agreement from -21.89 × 10^3^ to +32.89 × 10^3^, indicating a moderate level of agreement. A larger bias was observed between Alamar Blue and ATP, as well as MTT and ATP, with mean differences of 178.5 × 10^3^ and 159.7 × 10^3^ cells, respectively, and wider limits of agreement, reflecting a considerable variation in estimated cell counts between these methods.

The agreement between assays in the 3D GB models showed different characteristics. For Alamar Blue versus MTT, the average difference was -59 × 10^3^ cells, suggesting that Alamar Blue generally reported fewer cells than MTT. This pair had narrower limits of agreement ranging from -149.2 × 10^3^ to +31.16 × 10^3^. Similarly, the comparison between Alamar Blue and ATP assays indicated a mean difference of -140.4 × 10^3^ cells, with Alamar Blue also tending to estimate fewer cells. The limits here were substantial, from -451.4 × 10^3^ to +170.5 × 10^3^. The MTT versus ATP comparison in 3D showed an average difference of -81.44 × 10^3^ cells, with limits of agreement from -313.5 × 10^3^ to +150.7 ×10^3^.

### 3.2. Estimation of cell number by cell counting assays in static (1-21 day) 3D ECM scaffold-based cultures of U251 cells

#### 3.2.1. Overview of the results of all examined assays

In the other setup, we examined the performance of the cell counting and viability assays in the dynamic 3D Matrigel-based cultures of U251 cells. The number of available data points for DNA assay was 176, for the Alamar Blue assay it was 171, for MTT it was 252 and for ATP assay we had 186 data points, respectively. The DNA assay in Matrigel-based cultures was performed after recovering the cells from the gels to reduce the noise from the Matrigel matrix.

The overview of the results of the cell number estimation by different assays in dynamic 3D cultures (Matrigel-based and d-ECM based constructs containing U251 cells) is shown in Figure 6. The ECN for each time point was calculated based on the FC in assay readings relative to the average readings obtained on day 1 of in vitro culture. At this time point, approximately 12-18 h after cell seeding, the cells are well attached to the surface of the brain-derived dECM scaffolds in 3D TECs (see Figure A6, in Appendix A) or embedded in Matrigel. The effective cell seeding density for both, Matrigel constructs and TECs was 10,000 per construct. For each assay, the FC was determined by dividing the assay reading of a given sample by the assay reading obtained for the reference sample (1-day old matching type cell culture) containing approximately 10,000 live cells. According to the results presented in Figure 6, all assays in both types of examined 3D cultures detected gradual increase of cell numbers. In Matrigel constructs, the ECN on the 21^st^ day in vitro (DIV21) varied between 2 groups of assays. According to the DNA and Alamar Blue assays, it was about 500,000 cells per construct, while the estimation by the MTT and Alamar Blue assay was close to 225,000 cells. In the brain-specific TECs, the ECNs of MTT assay were similar with the DNA assay results, and showed about 100,000 cells per construct on DIV21. At the same time, the Alamar Blue and ATP assays estimated the cell contents in TECs on DIV21 as approximately 230,000 cells per construct. Interestingly, in 3D TECs, two groups of assays (DNA and MTT) and (Alamar Blue and ATP) showed different growth rates: while the first pair of assays demonstrated a relatively slow and steady growth along all the studied culture period, the second pair of assays showed an accelerated growth rate that started on DIV1. The detailed results of the ECNs obtained via different cell counting assays in dynamic scaffold-based 3D cultures of U251 cells are reported in Tables B10 (Matrigel constructs) and B11 (TECs).

**Figure 6.**
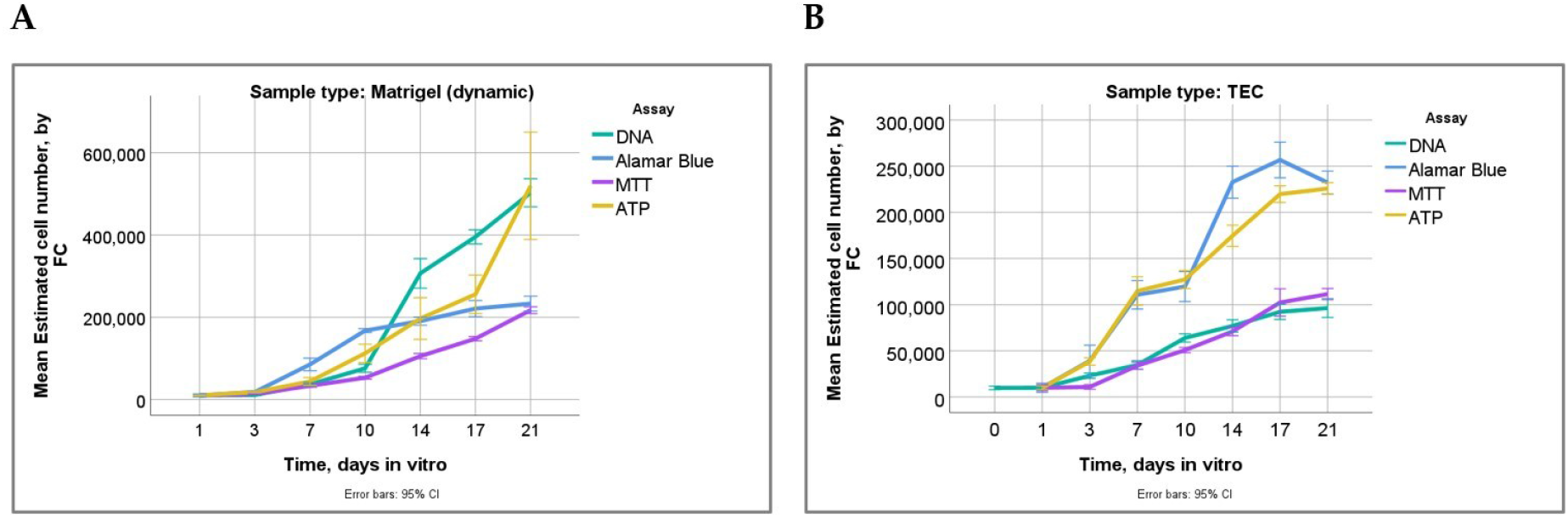
Overview of the dynamics of the estimated cell numbers (ECN) calculated using fold change (FC) of different assays’ readings in dynamic (1-21-days) experiments in 3D Matrigel-based constructs and TECs containing U251 cells. (A) 3D Matrigel-based constructs: note that the ECNs of all assays demonstrate increase with time, while providing different values on the same time points, expecting a few time points where the lines representing different assays overlap. At the latest time point (21 days), the ECNs detected by the DNA and ATP assay are relatively higher than that obtained by MTT and Alamar Blue assays. (B) 3D TECs: note the similarity between the ECNs obtained by DNA and MTT assays, which show overall lower ECNs than another group of assays with similar readings (Alamar Blue and ATP assays).

#### 3.2.3. Alamar Blue, MTT and ATP assays

Table 8 summarizes the results of Alamar Blue, MTT and ATP assays in the dynamic scaffold-based cultures of U251 cells.

**Table 8.** Cell number estimation by the Alamar Blue, ATP and MTT assays in in dynamic 3D Matrigel-based constructs and brain dECM-based TECs containing U251 cells.

| Time in vitro, days | ECN*, ×10<sup>3</sup> |  |  |  |  |  |
| --- | --- | --- | --- | --- | --- | --- |
|  | Matrigel |  |  | TECs |  |  |
|  | Alamar Blue | MTT | ATP | Alamar Blue | MTT | ATP |
| 1 | 10.00±4.80 | 10.00±1.23 | 10.01±2.41 | 10.00±5.83 | 10.00±8.39 | 10.000±4.801 |
| 3 | 18.21±3.93 | 13.18±3.39 | 17.67±2.90 | 23.10±6.78 | 10.92±6.37 | 38.453±10.162 |
| 7 | 85.17±38.94 | 33.26±10.24 | 43.83±23.69 | 34.74±10.52 | 33.95±9.96 | 114.836±38.913 |
| 10 | 167.58±11.87 | 52.90±10.04 | 112.29±52.86 | 63.95±10.70 | 50.83±7.15 | 127.270±24.574 |
| 14 | 190.68±25.19 | 105.38±18.80 | 196.76±128.13 | 76.97±15.77 | 71.00±11.40 | 174.638±29.209 |
| 17 | 221.10±49.65 | 147.86±15.53 | 255.89±118.26 | 92.27±9.84 | 102.35±37.62 | 219.763±22.906 |
| 21 | 233.10±45.52 | 217.22±24.26 | 519.53±329.97 | 96.8±24.03 | 111.48±15.24 | 225.845±15.984 |

\* The ECN was calculated by the FC of the assay readings relative to the DIV1-sample with the smallest cell count ( $\approx 1.0 \times 10^4$ ). Abbreviation: ECN – estimated cell number. The data presented as Mean ± St. Dev.

Several important observations can be done following the data presented in Table 8. The most obvious difference in the cell density achieved during the culture period between the Matrigel constructs and TECs that was noted from the DNA assay, emerges again from the outcomes of the metabolic assays. The TECs consistently demonstrate a lower cell number than the Matrigel constructs. The results of the statistical analysis of the ECN differences between two types of the scaffold-based 3D cultures are presented in Table B12 (Appendix B). In particular, statistically significant differences between the models was detected by Alamar Blue assay on all time points, excepting DIV1 and DIV21. The MTT assay revealed the different ECNs in Matrigel- and TEC systems on DIV14, 17 and 21. The ATP assay showed the statistically significant differences of ECN, depending on the culture platform, on all time points, excepting DIV1.

Next, we explored the differences in ECNs obtained by different assays within the same type of ECM scaffold-based 3D cultures of U251 cells using the independent sample Kruskal-Wallis. It was found that in both Matrigel constructs and TECs, on DIV1, the ECNs obtained by different assays were not statistically significantly different. However, from DIV3 onwards, significant differences emerged (p<0.001). There was a clear trend of bigger differences between the assays’ ECNs with longer cell culture time. The details on the ranks of ECNs and pairwise comparisons between the assays at different time points are presented in Tables B13, B14 and B15 in Appendix B, with the statistically significant differences highlighted.

The following results were obtained by correlation analysis. In Matrigel constructs, the Alamar Blue and MTT assays showed very strong and highly statistically significant (p<0.001) Spearman’s correlation coefficients (Rs=0,922 and Rs = 0.975, respectively) comparable with the coefficient detected for the DNA assay (Rs=0.956), reported above, with the number of days in culture. The Rs for ATP assay also showed moderate strong positive and highly statistically significant link (Rs=0.776) between the assay’s readings and the time in culture. The same trend was notable in TECs, with the strongest correlation between the time in vitro and MTT assay’s ECNs (Rs=0.950), followed by the ATP assay (Rs=0.939) and Alamar Blue (Rs=0.856), all with p<0.001. The respective Rs for the DNA assay in TECs was 0.927 and p<0.001. These findings indicate that time in culture may serve as a potential predictor of ECNs in dynamic ECM scaffold-based assays, which is essential for the following quantification of the assay performance.

#### 3.2.4. Quantification of the cell counting assays’ performance in dynamic 3D Matrigel-based cultures and TECs

##### Analysis of the assays’ linearity (linear regression models)

To quantify the linearity of the assays we employed linear regression modelling, exploring the relationship between the ECNs detected by the assays and the time in culture, measured in days. The interpretation of the different parameters and statistics of such modelling approach was given in section 3.1.4 above. Here, a few modifications of this interpretation include the meaning of the intercept as a number of cells expected on the zero time point, and the coefficient B signifying the approximate increase of cell numbers per day. The results of the linear regression analysis for the ECNs vs time in culture in dynamic 3D Matrigel and TEC GB models are shown in Table 9. The visual representation of the linear regression models fitting to the experimental data is shown in Figure B4 (Appendix B).

**Table 9.** Summary of linear regression model outcomes for the estimated cell number via FC according to the assays’ readings in dynamic (1-21 days) 3D (Matrigel-based) cultures of U251 cells.

| Assay and sample type | R Value | R<sup>2</sup> | aR<sup>2</sup> | Std. Error of the Estimate | Intercept | B* | Sign. (p) |
| --- | --- | --- | --- | --- | --- | --- | --- |
| Alamar Blue (Matrigel) | 0.892 | 0.795 | 0.794 | 39,806.77153 | 5,566.175 | 12,220.207 | <0.001 |
| Alamar Blue (TEC) | 0.858 | 0.736 | 0.734 | 48,361.11726 | 14,626.921 | 12,497.421 | <0.001 |
| MTT (Matrigel) | 0.949 | 0.901 | 0.901 | 23,122.44319 | -23,980.411 | 10,241.923 | <0.001 |
| MTT (TEC) | 0.906 | 0.821 | 0.820 | 17,682.34214 | -19,09.419 | 5,532.823 | <0.001 |
| ATP (Matrigel) | 0.716 | 0.512 | 0.509 | 154,862.69181 | -74,180.140 | 23,014.909 | <0.001 |
| ATP (TEC) | 0.940 | 0.884 | 0.883 | 27,784.64195 | 13,264.739 | 11,204.864 | <0.001 |

*\*B is the Coefficient (B) for the Time in culture (measured in days).*

The main implications of the data presented in Table 9 are the following. Firstly, for all the tested assays and for both 3D ECM scaffold-based cell cultures, the linear regression models are highly statistically significant (p<0.001), indicating the experimental data, in general fits the linear trends. This also means that the duration of culture is a reliable predictor of the cell numbers estimated by FC of assays’ readings. The models show a high positive correlation between the ECN and the time in culture, with the only exception of the ATP in Matrigel that had R=0.716 (moderate correlation strength). The linearity of the assays is characterized by R^2^ and adjusted R^2^ values, which vary between approximately 51% (for Alamar Blue assay in Matrigel) up to 90% (for MTT assay in both 3D scaffold-based culture types). In Matrigel-based constructs, the highest linearity was achieved for MTT assay (aR^2^=0.901), followed by the DNA assay (aR^2^=0.850), Alamar Blue (aR^2^=0.794) and ATP (aR^2^=0.509). assays. In 3D TECs, the ATP assay showed the highest linearity metric (aR^2^=0.883), followed by DNA assay (aR^2^ = 0.844), MTT (aR^2^=0.820) and Alamar Blue assay (aR^2^=0.734).

Interestingly, R^2^ and aR^2^ values are quite close between the Matrigel constructs and TECs for DNA assay, Alamar Blue and MTT assays (with the TECs’ ones a little bit lower), while for the ATP assay the trend is clearly reversed as it is visible from the higher R^2^ and aR^2^ in TECs compared to 3D Matrigel-based models.

Another remarkable observation is that the absolute values of the intercepts for the models reflecting the behavior of the assays in Matrigel-based 3D cultures of U251 cells are large, excepting the Alamar Blue assay. This indicates a relatively high noise vs signal ratio in cell-free Matrigel. At the same time, some intercepts are negative (DNA and ATP assays in Matrigel, and MTT in both 3D scaffold-based platforms) also pointing at a potential influence of noise on the detection in the cultures with low cell numbers.

The cell density growth rates estimated by the coefficient B are demonstrating certain disagreement between the results of different assays withing the same culture type. For the Matrigel platform, MTT assay indicates the lowest cell growth rate with a B=10,241.923 (cells per day per construct), followed by Alamar Blue assay (B=12,220.207), ATP assay (B=23,014.909) and the DNA assay (B=26,600.613). In 3D brain dECM-based TECs of U251 cells, DNA assay reflected the lowest cell number increase per day (B=4,482.135), followed by MTT assay (B=5,532.823), ATP assay (B=11,204.864), and Alamar Blue assay (B=12,497.421). Taken together, these results also reveal that the cell culture platform affects the estimation of cell numbers by different assays in different manner.

##### Analysis of the assays’ variation

The result of the analysis of the technical replicability, biological reproducibility and the overall precision of the cell counting assays, presented via the values of the technical, biological and overall coefficients of variation, CVt, CVb, and CV, respectively, are shown in Figure 7. Detailed data on the values of the CVt, CVb, and CV for different assays and across two 3D culture platforms is listed in Table B16 (in Appendix B).

**Figure 7.**
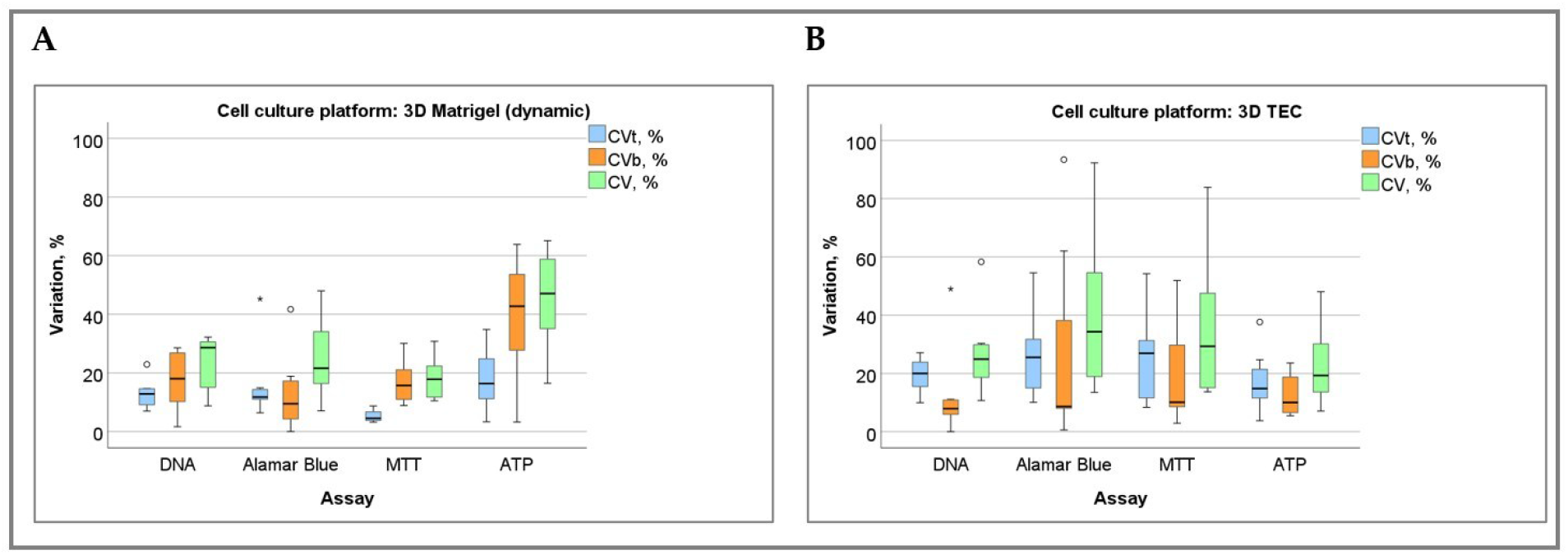
The coefficients of variation (CVt, CVb and CV) obtained for different cell counting assays in in in dynamic (1-21-day) 3D Matrigel-based constructs and brain dECM-based TECs of U251 cells. (A) 3D Matrigel-based constructs; (B) 3D TECs.

In dynamic 3D Matrigel-based GB models, the MTT assay demonstrated the lowest CVt of 5.37%±2.19%, suggesting the best technical repeatability in measurements. The DNA assay showed a moderate technical variation with a CVt of 12.89%±5.38%, while the Alamar Blue and ATP assays exhibited a higher technical variability, with CVts of 16.31%±13.047% and 18.11%±10.84%, respectively. Biological variation was the highest for the ATP assay (CVb=38.94%±21.37%). DNA and Alamar Blue assays had lower CVbs at 17.48%±10.48% and 13.48%±14.157%, respectively. The overall variation reflected by CVs followed the same pattern, with the ATP assay being the highest at 45.22%±18.66% and the MTT assay being the lowest at 18.18%±7.73%. The DNA and Alamar Blue assays and had CVs at 23.01%±9.77% and 25.37%±15.626%, respectively.

In TECs, technical variability (CVt) tended to be higher across assays than in Matrigel-based cultures. It was the high for the Alamar Blue assay (26.21%±15.22%). For MTT and Alamar Blue assays assay it was not only high, but also varied a lot itself (CVb=20.21%±20.56% and 27.86%±35.47%, respectively). In contrast, ATP assay demonstrated relatively good biological reproducibility with CVb=12.82%±7.40%. The overall CV for TECs followed the pattern of technical variation: the Alamar Blue and MTT assays showed the highest overall variations at 41.02%±30.35% and 36.01%±26.50%, respectively. The overall CVs of the ATP and DNA assay were 23.12%±14.26% and 27.26%±15.35%, respectively.

To compare the variation of the assays, we employed the independent samples Kruskal-Wallis’ test with pairwise comparisons (Tables B17, B18, and B19 in Appendix B). It was found that all the assays across each of the examined cell culture platforms show comparable technical variability, excepting the pair of MTT and ATP assays in the dynamic Matrigel-based models, where the CVt of MTT was statistically significantly lower than CVt of the ATP assay (p=0.015). The biological reproducibility also did not significantly differ between the assays within the same cell culture platform with again a single borderline exception: in Matrigel constructs, the Alamar Blue assay demonstrated lower CVb assay vs ATP assay (p=0.046). Finally, the overall coefficients of variation were not statistically different among the examined assays, excepting the pair of MTT and ATP in Matrigel-based cultures, where the CV of ATP assay was statistically significantly higher than that of the MTT assay (p=0.042).

The comparisons of the coefficients of variation for the same assays across the cell culture platforms (Matrigel constructs and TECs) showed no statistically significant differences in the technical replicability, biological reproducibility and overall variation of the DNA, Alamar Blue, and ATP assays. MTT assay showed a statistically significantly lower technical variation (CVt) in Matrigel constructs than in TECs (p=0.003), but other metrics of the variation of this assay were similar across two cell culture platforms.

##### The differences between Analysis of the inter-assay agreement (Bland-Altman plots)

Figure 8 demonstrates the Bland-Altman plots reflecting the extent of agreement between two selected assays. In panels A-F, the results obtained for the assays in the dynamic 3D Matrigel-based cultures are shown. Panels G-L depict the plots for the assays performed in 3DTECs. As it is visible from the figure, in Matrigel constructs, the closest agreement (and the lowest mean difference) was observed between the DNA and ATP assays (26.01×10^3^ cells in average for the measurements of 2 assays on the same time point). The Alamar Blue assay showed similar levels of agreement with ATP assay (mean difference -32.87×10^3^ cells). The most notable disagreement was observed between the DNA assay and MTT (mean difference 108.3×10^3^ cells), while other pairs of assays demonstrated intermediate results in this comparison. In 3D TECs, the exceptionally high agreement was revealed between the DNA and MTT assays (with mean difference of 0.52×10^3^ cells, and the limits of agreement of approximately ±20×10^3^ cells). Another pair with comparatively good agreement was Alamar Blue and ATP assays (mean difference 12.97×10^3^ cells). Other tested pairs of assays showed the absolute values of mean differences varying between 73.77×10^3^ cells and 7.27×10^3^ cells.

**Figure 8.**
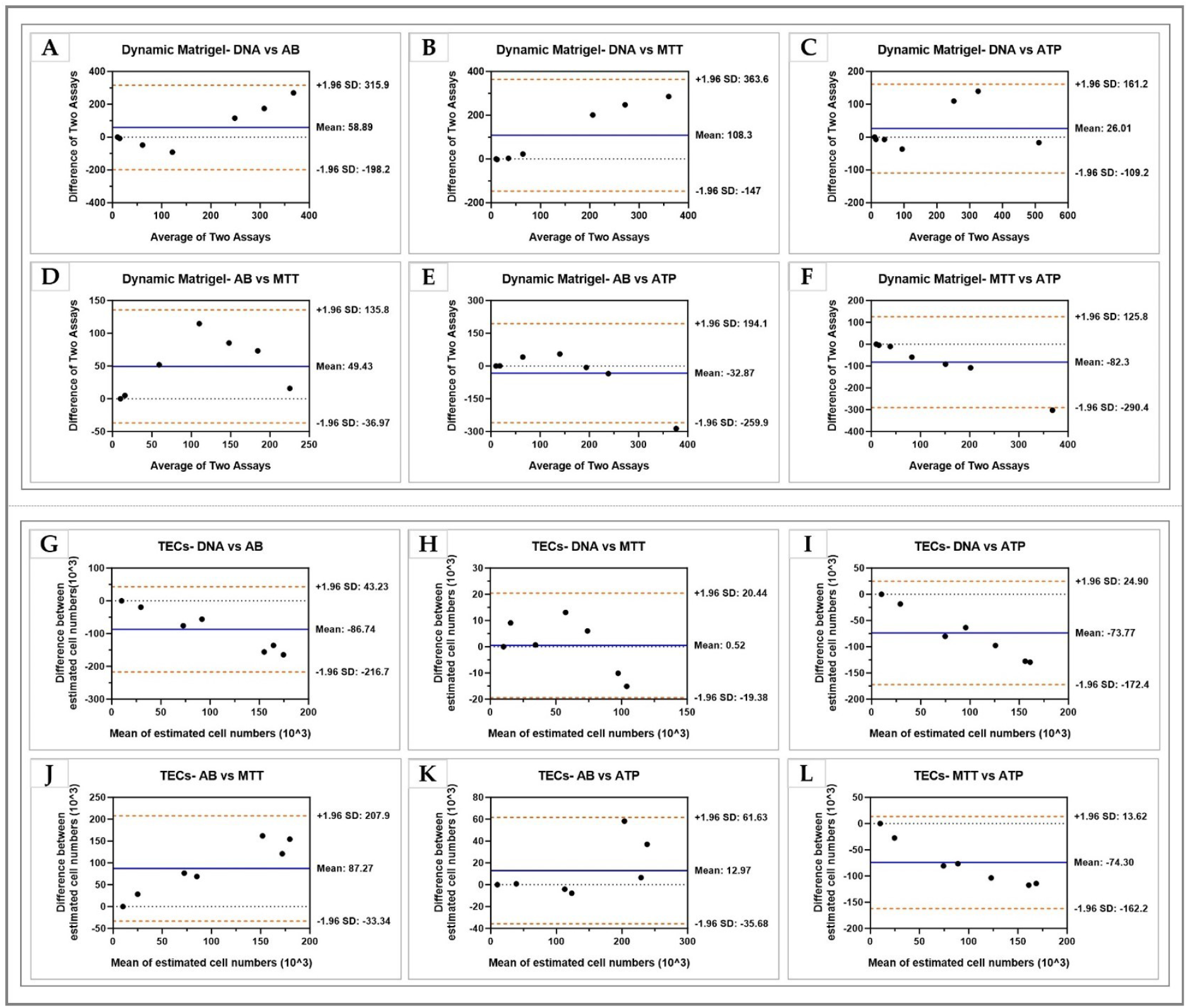
Assessment of the agreement between the metabolic assays regarding the estimated cell number in dynamic 3D Matrigel-based constructs and brain dECM-based TECs of U251 cells by Bland-Altman plots. The X-axis represents the average estimated cell numbers (ECN) (in thousands) obtained by the pairs of different assays at the same time point. The Y-axis shows the difference between the ECNs (in thousands) by the two assays under comparison. The “Mean” indicates the difference between the assays’ ECNs (in thousands) is depicted as a horizontal blue line. The dashed orange lines demonstrate the positions of the limits of agreement (LoA) that correspond to the boundaries of the CI_95%_ for the mean difference. **(A-F)** Bland-Altman plots for the 3D Matrigel-based constructs; **(G-L)** Bland-Altman plots for 3D TECs. The following assay pairs are shown: **(A, G)** DNA vs Alamar Blue assays; **(B, H)** DNA vs MTT assays; **(C, I)** DNA vs ATP assays; **(D, J)** Alamar Blue vs MTT assays; (E, K) Alamar Blue vs ATP assays; **(F, L)** MTT vs ATP assays.

## 4. DISCUSSION

Our study addresses an important aspect of tumour modelling in vitro and tissue engineering in general such as reliable counting of live cells within the growing cell cultures. The standard approaches in this area were developed and adapted for the use in conventional monolayer and suspension in vitro culture conditions. One of the main advantages that such methodology allows is the application of plate reader-based protocols. By employing plate readers, the researchers are armed with a tool for performing multiple data points acquisition at a short time, eventually providing the access to the high-throughput cell counting. This opportunity is exceptionally important for the testing of new treatment protocols in experimental pharmacology and cancer research. The plate reader-based cell counting relies on the detection of different types of optical signals emerging via the interaction of the sample with light and detected as transmitted or reflected signal of certain intensity. The main detection modalities in plate readers are absorbance (corresponding to the measurement of the optical density of the sample via the attenuation of the transmitted light) and fluorescence / luminescence (based on the detection of the intensity of the certain wavelength light excited within the sample and emitted under exposure to a light of the optimal range). The plate reader-based cell counting methods are presented, accordingly, by a series of mostly commercially available assays and reagents with an established relationship between the intensity of the emitted or transmitted light signal and the number of cells contained in the sample. As detailed in the Introduction, plate reader-based cell counting methods rely on the detection of several types of biological compounds or processes, such as DNA, mitochondrial activity, or redox activity, for which the linear relationship with cell number or viability were justified ^[5, 20]^.

While been well adapted and validated for the monolayer and cell suspension in vitro cultures, these conventional approaches in cell counting are challenged in 3D in vitro culture conditions, where the new light penetration, absorption and scattering conditions emerge due to the presence of the different spatial distribution of the cells, for example, the clustering, or because of the presence of the highly scattering or absorbing materials in the cellular microenvironment. The most notable example of such materials is extracellular matrix (ECM) or ECM-mimicking biomaterials. The ECM-light interaction is currently under study; however, these findings are just starting to enter the translation into the field of accessible research tools. As a result, plate reader-based assays in the ECM-containing scaffold-based 3D in vitro tissue models present an important modern methodological challenge ^[21]^.

At the same time, the development of tissue engineering and 3D culture have resulted in the appreciation of the importance of ECM in cell and tumour biology, including the cell viability, growth and metabolic rates, differentiation, and treatment responsiveness and resistance ^[22]^. A recent literature analysis ^[16]^ has revealed that despite a high interest to the 3D in vitro models of GB as drug development testbeds, the majority of the GB studies that employed 3D in vitro models of this tumour and where the clinically relevant results were achieved (for example, in terms of the doses, schedules and concentrations of the administered drugs), were performed in multicellular spheroids or similar scaffold-free systems. At the same time, 3D scaffold-based models of GB that contained CM or ECM-mimicking materials mostly demonstrated low or lack of the treatment effect of the drugs when the clinically reasonable and not very high concentrations or doses of the drugs were used. One of the common assumptions and observations related to this type of findings is that the 3D in vitro environment or ECM, or their combination affect the tumour cells’ drug sensitivity and responsiveness. Indeed, there are multiple examples in the literature that, seemingly, justify this ^[8, 23]^. However, we found this surprising that it was not a common idea in the current cancer research that not or not only cell’s responses are changing in 3D in vitro tumour models, but also the assays that we employ (in particular, for cell counting, viability, or cytotoxicity testing) may have different performance in the more complex environments that are created in advanced in vitro tissue models. This idea has motivated our current study.

To the best of our knowledge, the current study presents the first an in-depth quantitative and comparative analysis of the performance of the four commercially available plate reader-based cell counting assays in 2D (monolayer) and 3D ECM scaffold-based cell culture environments.

We observed that the assays’ performance varied notably across different platforms (static and dynamic 3D Matrigel-based cultures, and 3D organ-specific TECs). The assays demonstrated linear relationships with cell numbers and culture duration, but with varying degrees of coefficients of variation (CVs) and precision indices (PIs). The assays’ performance in cross-assays and cross-platform comparisons revealed significant differences, underscoring the complexity of 3D cellular environments. The summary of the observations is shown in Table 10.

**Table 10.** Overview of the metrics of the cell counting assays’ performance in the studied cell culture conditions.

| Assay | Culture Model | | R (Pearson's correlation coefficient) | Linearity ($R^2$) | Linearity adjusted ($aR^2$) | Overall CV, % | CVt, % | CVb, % | PI |
| --- | --- | --- | --- | --- | --- | --- | --- | --- | --- |
| Alamar Blue | 2D |  | 0.987 | 0.973 | 0.973 | 22.31±14.35 | 12.21±9.20 | 11.67±10.08 | 1.73±0.11 |
|  | 3D | MG stat. | 0.825 | 0.680 | 0.674 | 26.51±19.95 | 12.59±9.84 | 17.25±19.01 | 0.35±0.26 |
|  |  | MG dyn. | 0.892 | 0.795 | 0.794 | 25.37±15.63 | 16.31±13.05 | 13.48±14.16 | N.A. |
|  |  | TECs | 0.858 | 0.736 | 0.734 | 41.02±30.35 | 26.21±15.22 | 27.86±35.47 | N.A. |
| MTT | 2D |  | 0.973 | 0.947 | 0.946 | 22.17±5.57 | 13.14±6.29 | 16.02±2.19 | 1.61±0.24 |
|  | 3D | MG stat. | 0.967 | 0.934 | 0.933 | 29.79±39.79 | 21.36±31.80 | 10.98±19.31 | 0.97±0.24 |
|  |  | MG dyn. | 0.949 | 0.901 | 0.901 | 18.18±7.73 | 5.37±2.19 | 16.98±7.78 | N.A. |
|  |  | TECs | 0.906 | 0.821 | 0.820 | 36.01±26.50 | 25.04±16.20 | 20.21±20.56 | N.A. |
| ATP | 2D |  | 0.928 | 0.862 | 0.86 | 18.59±13.26 | 2.57±0.96 | 16.92±13.68 | 0.86±0.12 |
|  | 3D | MG stat. | 0.93 | 0.865 | 0.862 | 28.05±15.80 | 1.33±1.64 | 26.04±13.82 | 1.02±0.15 |
|  |  | MG dyn. | 0.716 | 0.512 | 0.509 | 45.22±18.66 | 18.11±10.84 | 38.94±21.37 | N.A. |
|  |  | TECs | 0.94 | 0.884 | 0.883 | 23.12±14.26 | 17.45±11.13 | 12.82±7.40 | N.A. |

As shown in Table 10, all the examined assays demonstrated high linearity in 2D culture (aR^2^ varied between 0.860 and 0.991). In static 3D Matrigel, there was notable loss of the performance quality of DNA assay due to the high noise from the DNA contained in the gel, while other assays showed aR^2^ in the range from 0.674 (Alamar Blue) to 0.933 (MTT). In dynamic 3D Matrigel constructs, aR^2^ values varied across assays from 0.509 (ATP assay) and 0.901 (MTT). In 3D TECs, linearity of the assays changed between aR^2^ equal to 0.734 (Alamar Blue) and 0.833 (ATP assay). These findings demonstrate that there is no universal choice of the most linear assay that would be suitable for all types of cell culture platforms. Therefore, other metrics should be involved. One of them, which is available for all the tested conditions, is the value of the overall coefficient of variation. As we have revealed, it is influenced by technical and biological variation to different extent, depending on the assay and the cell culture conditions, but it is always higher than the individual coefficients such as CVt and CVb and the, it can employed as a more comprehensive measure of the assays’ performance, even in the dynamic scaffold-based cultures, where the number of cells in the construct can be only predicted since the initial seeding through the regression models that use time in culture as an independent variable. Considering this, the ranking of the assays’ performance in different cell culture conditions can be performed by superimposing the linearity measure (R^2^ or aR^2^) and the overall CV values. In the current study, CV of different assays varied between 5.88% (DNA assay, 2D culture) and 45.22% (ATP assay, 3D dynamic Matrigel constructs), representing the best and worst performance, respectively. By the combination of the linearity and overall variation metrics, the best performance was demonstrated by DNA and Alamar Blue assays in 2D culture, MTT assay in the static and dynamic 3D Matrigel constructs, and the ATP assay in the 3D dynamic brain-specific GB TECs.

The diverse performance metrics (R^2^ and aR^2^, CVs, PIs) across different assays and platforms underscore the necessity for a comprehensive evaluation of assay suitability in 3D culture environments. This is particularly critical for applications in drug screening and tissue engineering, where assay accuracy and precision are paramount. Our study’s cross-assay and cross-platform analyses provide a framework for selecting appropriate assays based on specific research needs and culture conditions.

Our study focused on specific types of 3D cultures, a specific cell type, and a range of assays, which may limit the generalizability of our findings. Future studies could broaden the range of cell types and 3D culture models. Additionally, investigating the underlying reasons for the observed variability in assay performance across different platforms would provide deeper insights into the complexities of 3D cell culture. Therefore, our results rather emphasize the need for careful assay selection and interpretation in different culture models, especially when dealing with dynamic 3D systems. The variability in CVs and PIs across assays suggests that no single assay can be universally applied across all culture types without consideration of its specific limitations and the context of the experiment.

The observed variability in assay performance across different culture models highlights the complexity of 3D cell cultures and the challenges in obtaining consistent and reliable assay results. In 2D cultures, the DNA assay demonstrated remarkable precision, which is in contrast to its performance in 3D cultures, particularly in the static Matrigel. This finding aligns with the notion that 3D cultures present a more complex environment for cell proliferation and viability assessments ^[11b]^.

The high CVs in static and dynamic 3D cultures, especially for the DNA, Alamar Blue, and MTT assays, reflect the inherent challenges in standardizing assays in 3D environments.

Future research should focus on developing more standardized and reliable methods for assay application in 3D cultures. Investigating the underlying causes of variability in assay performance, particularly in dynamic systems like TECs 3D, will be crucial for enhancing assay reliability and precision in these complex environments.

## CONCLUSIONS

Overall, our study contributes significantly to the understanding of cell viability and counting assays in 3D cultures. The findings highlight the need for careful consideration of assay choice and interpretation in various 3D culture conditions, paving the way for more accurate and reliable research in the fields of bioengineering and cancer research.

## Author Contributions

Conceptualization, M.V., A.N., and A.G; methodology, M.V., A.N., A.D.I., B.H., and A.G; validation, M.V.; formal analysis, M.V. A.I., A.N., and A.G.; investigation, A.G.; resources, A.N. and A.G.; data curation, M.V. and A.G.; writing—original draft preparation, M.V., A.N., A.I., B.H., and A.G.; writing— review and editing, M.V., A.N., A.I., V. L., A. D.I., B.H. and A.G..; visualization, M.V., A.I., and V.L.; supervision, A.N., A. D.I., B.H. and A.G.; project administration, A.N., B.H.; funding acquisition, A.N., B.H., and A.G. All authors have read and agreed to the published version of the manuscript.

## Funding

This research was funded by Tour De Cure charity foundation (Australia),via Pioneering Cancer Research Grant awarded to A.N. and A.G. A.N. was supported by NHMRC Early Career Research Fellowship. The work of A.I. was supported by Russian travel scholarship for academic excellence. A.G. thanks Macquarie University for supporting her work via Macquarie University Research Fellowship. M.V. is grateful to Macquarie University for RTP PhD Scholarship.

Institutional Review Board Statement: Not applicable.

Informed Consent Statement: Not applicable

## Data Availability Statement

The raw research data used in this study is available from the corresponding authors via a reasonable request.

## Acknowledgments

Authors sincerely thank the PC2 Lab Operations team of the Macquarie University Faculty of Medicine, Human and Health Sciences for the exceptional everyday support of our experimental work over the years. The authors would like to acknowledge the Macquarie University Faculty of Science and Engineering Microscope Facility (MQFoSE MF) for access for its instrumentation and staff. M.V. thank Bavani Gunasegaran and Mina Ghanimi Fard for their assistance during the high labour load experiments. Authors thank Gilles J Guillemin for helpful discussions. A.G. acknowledge UNSW Microscopy Unit for the support in SEM microscopy.

