## Appendix A for "What Can We Count On? Performance of Microplate Cell Counting Assays in 2D Monolayer and 3D ECM-based In Vitro Tumour Models"

### *A1. Decellularization of sheep brain tissue*

Acellular brain ECM scaffolds were prepared from fresh sheep brain tissue by decellularization (DCL) as described elsewhere [1] with minor modifications [2]. Briefly, sheep (lamb) brain tissues obtained fresh-frozen from a local butchery, thawed at 4 °C, and washed with PBS. The hemispheres were thoroughly dissected and separated from corpus callosum, medulla, fornix, and spinal cord, and separated in sagittal plane. Then, the hemispheres were sliced parallel to the frontal plane using surgical blades into coronal section fragments of an average thickness of 1.5 - 2 cm. Eight to ten fragments were obtained per hemisphere. Each sliced brain hemisphere fragment was placed individually into a 50 mL Falcon conical centrifuge tube and filled with 0.1% v/v water solution of sodium dodecyl-sulphate (termed "decellularization solution") up to the total volume of 35 mL. The presence of the 15 mL of empty space in the tubes was essential for proper agitation of the samples. The tubes were secured on the orbital shaker and continuously shaken at 90 rpm for minimally one week until the tissue and the solution demonstrated characteristic changes indicating completion of the immersion-agitation DCL [1b] process. Specifically, the decellularized tissue acquired a felt-like texture and became white and semi-translucent, while the DCL solution changed its appearance from turbid to clear. The DCL solution was changed twice daily in aseptic conditions.

Next, the decellularized brain tissue fragments were aseptically transferred to new tubes and washed on the same shaker with 1% Antibiotic-Antimycotic (AA) (Sigma-Aldrich) solution in 1% PBS on sterile water (termed "washing solution") to remove the detergent residuals for 1-2 days. After a few times changing of the washing solution, when there were no more foam or cloudiness observed in the tubes, the decellularized brain tissue fragments were aseptically transferred into new 50 mL Falcon tubes, which were filled for the full volume with fresh washing solution, and then were transferred to a fridge (+4 °C) until use.

### *A2. Preparation, sterilization, and conditioning of acellular brain scaffolds*

Next, a two-step sterilization procedure of the decellularised brain tissue was applied. All the operations were performed aseptically in a biosafety cabinet. For chemical sterilization, the fragments of decellularized sheep brain hemispheres were soaked for 2 hours in a sterilizing solution containing 0.02% (v/v) peracetic acid (PAA) in 4% (v/v) ethanol in filter sterilized Milli-Q water, at room temperature. After the incubation, decellularised brain tissue fragments were washed 3 times with sterile PBS and placed into a new sterile Petri dish. Next, the scaffolds were made by cutting of the sterilized decellularized brain hemispheres' tissue fragments into pieces of approximately 2 mm × 2 mm × 2 mm using surgical blades. Ten to fifteen scaffolds were obtained from each section of the decellularized brain hemisphere slice.

For further sterilization, the scaffolds were placed on a thin sterile transparent polyethylene film which allows passing the ultraviolet (UV) light, and every scaffold was spread out as much as possible (the mean diameter of the spread scaffolds was 3.9±0.3 mm, and the mean thickness was ≈ 2 mm). After that, each scaffold was covered with a drop of sterilize PBS to avoid drying and sterilized with ultraviolet light for 30 minutes in the biosafety cabinet. Then, the scaffolds covered with another layer of sterile polyethylene film and the resulting "sandwich" was flipped to sterilize the other side of the scaffolds with UV light for a further 30 minutes.

Following that, the sterilized brain dECM scaffolds were aseptically placed one per well into 24 well-plates, and 1 mL of complete media was added into each well plate containing a scaffold. Then, the scaffolds were preconditioned in the tissue culture incubator at 37°C and 5% $CO_2$ for at least 12-15 hours or until use. The plates were regularly monitored for potential microbial contamination by observing the colour of media and inspecting the wells under microscope. The signatures of the acceptable level of the scaffolds' sterility were the absence of the media turbidity or colour change to orange, brown or yellow from red or pink, and absence of the conventional visible signs of the cell culture contamination.

The absence of cellular material in the obtained brain dECM scaffolds was controlled and confirmed by histological examination (see Section 2.8.). In addition, as a part of the main experiment, the DNA content was measured in a total of nine scaffolds using a Quant-iT PicoGreen dsDNA assay as described below in Section 2.6.1.

### *A3. Methods of the morphological characterisation of 3D TECs and Matrigel-based constructs*

#### *A3.1. Histological and immunohistochemical examination and confocal imaging of 3D TECs*

In the current study we examined histological features of 3D TECs obtained by growing of human glioblastoma cells U251 on acellular sheep brain scaffolds. After 14 days of in vitro culture, the TECs were fixed in 10% neutral buffered formalin for 24 h, and then frozen in O.C.T. media cryosectioned using Cryostat (model). Cryosections of the TECs (6 μm in thickness) were stained with hematoxylin and eosin (H&E) for general structure, by Masson's trichrome method (MTCh, for collagen) and with toluidine blue (for acid glycosaminoglycans) by routine protocols. Stained sections were embedded in DPX mounting medium and closed with coverslips. Stained histological samples were studied using an upright research microscope Axio Imager Z2 (Zeiss, Germany) equipped with dry-air EC Plan-Neofluar (5×/NA0.16; 10×/NA0.30; 20×/NA0.50 Ph) and oil-immersion α Plan Apochromat (100×/NA1.46 oil) objectives (Zeiss, Germany). Images were recorded using a preinstalled microscope digital video camera AxioCam (1388×1040, Zeiss, Germany) in a single-frame and stitching modes using Zen proprietary software (Zeiss).

The immunohistochemical staining was performed to compare and verify the phenotype of U251 cells seeded in 2D monolayers and in 3D TECs. The cells were stained for glial fibrillar acidic protein (GFAP), the structural protein of the intermediate filaments of astrocytes [3]. For GFAP staining, the BD Pharmingen™ Purified Mouse Anti-GFAP Cocktail (#556330, BD Biosciences) was applied in combination with the Donkey anti-Mouse IgG (H+L) Highly Cross-Adsorbed Secondary Antibody, Alexa Fluor™ 488 (#A-21202, Thermo Fisher Scientific).

The LIVE/DEAD Viability/Cytotoxicity Kit (Invitrogen, #L3224) was applied for the differential staining of the live and dead cells according to the manufacturer's protocol. Live cells stained with calcein-AM were visualized using a green false-palette, and the red colour depicted dead cells stained with ethidium homodimer-1.

An analysis of cellular morphology and viability was conducted using confocal microscopy. The study utilized a Zeiss LSM 880 IndiMo Axio Observer laser scanning confocal microscope (Zeiss, Germany) equipped with a Plan-Apochromat 40×/1.3 N.A. oil DIC UV-IR M27 lens for imaging procedures. The overview snapshots of the 3D TECs were obtained using Aperio XT Slide Scanner (Leica Biosystems, Wetzlar, Germany) equipped with 20×/0.75 Plan Apo objective, allowing the magnification 40x is obtained with a 2x magnification changes to 20×: 0.50 μm/pixel and 40×: 0.25 μm/pixel.

*A3.2. Scanning electron microscopy and image analysis in 2D and 3D cultures*

Three samples of each of the tested systems, including 2D cultured U251 cells, non-seeded Matrigel and brain dECM scaffolds, 3D Matrigel constructs and TECs fixed in 2.5% glutaraldehyde at 4°C overnight. Then, they were dehydrated through a graded ethanol series. After drying in LEICA EM CPD 300 Critical Point Dryer (Leica Microsystems GmbH, Wetzlar, Germany), the samples were mounted on aluminium specimen holders and coated with gold with Emitech K550 Gold Sputter Coater (Emitech Ltd., Ashford, Kent, UK). Scaffolds microstructures were visualized at magnifications of 500, 5000, 10000, 20000 using a JEOL JSM 7100F FESEM (Tokyo, Japan) at an accelerated voltage of 15 kV.

### *A4. Results of the morphological characterisation of the dECM scaffolds, 3D TECs and Matrigel-based constructs*

The results of histological examination of the fibro-porous brain dECM scaffolds and 3D TECs of U251cells cultured on the brain dECM scaffolds are shown in Figure A1.

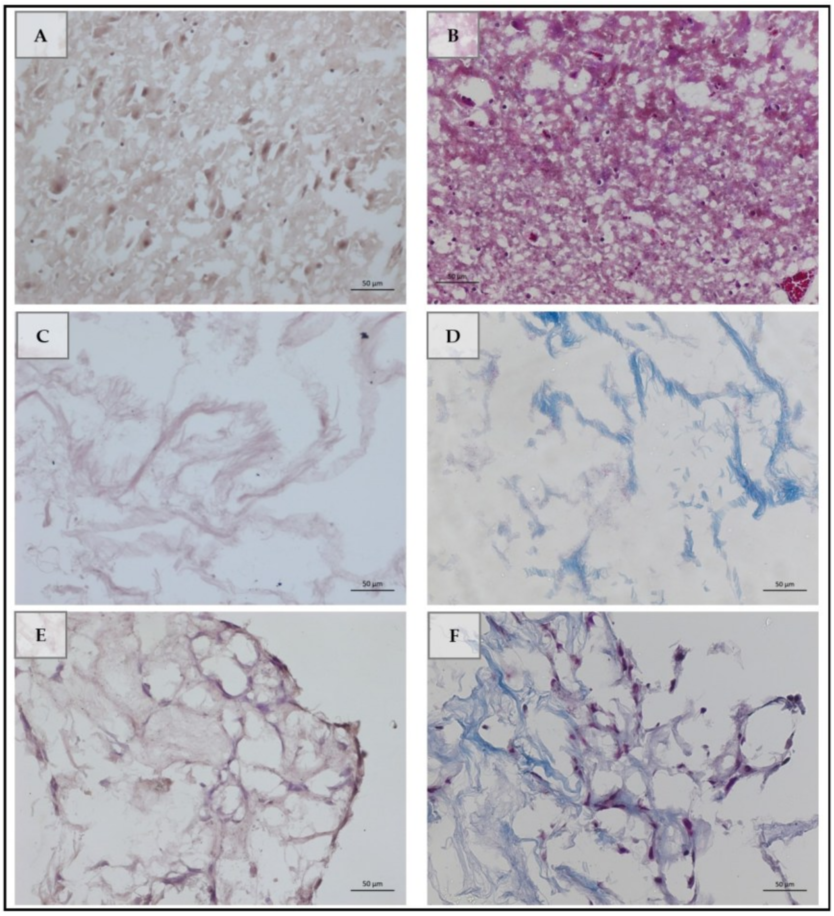


**Figure A1.** Histological examination of the native sheep brain tissue **(A, B)**, decellularised sheep brain hemispheres **(C, D)** and the 3D TECs of U251 cells cultured on the brain dECM scaffolds for 14 days. Samples were stained with hematoxylin and eosin **(A, C, E)** or Masson's trichrome method **(B, D, F)**. Scale bars 50 μm. Note the complete removal of cellular elements of the native brain tissue by decellularisation and the resulting fibro-porous structure of the dECM brain scaffolds **(C, D)**. Blue staining of the scaffolds' elements **(D, F)** reveals presence of fibrillar collagens. In 3D TECs, cells are attaching to the surface of the scaffold and gradually colonize it, penetrating to the deeper parts of the substrate and forming a network structure that to a notable extent follows the fibrous components of the scaffolds.

To demonstrate the gradual changes occurring in the 3D TECs depending on the time, we employed a histological scanner allowing to overview the structure of the whole TECs and see the extent of the scaffolds' colonization by U251 cells (Figure A2).

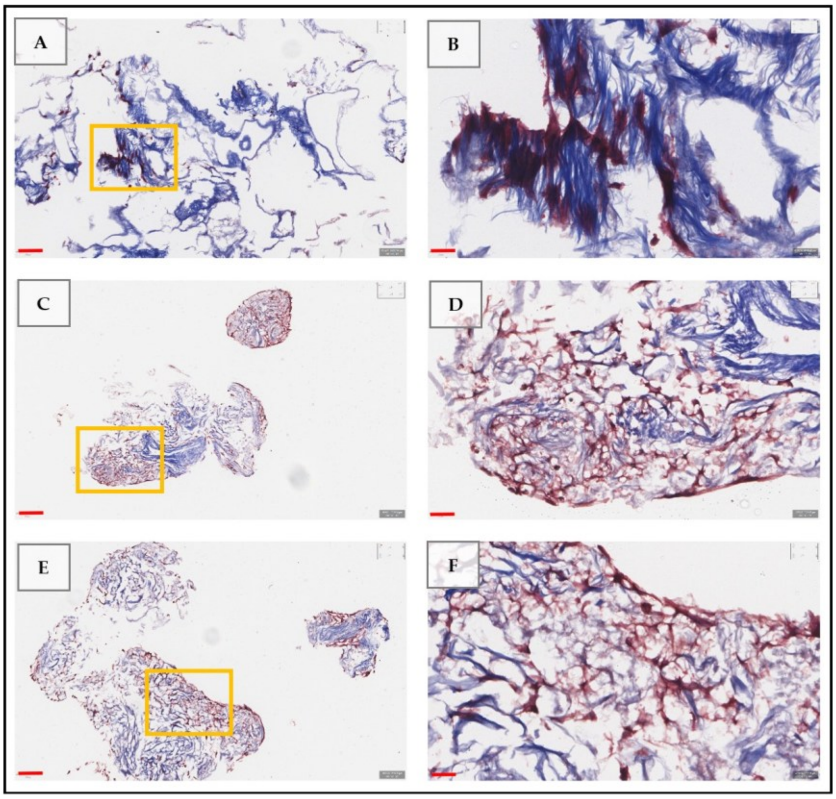


**Figure A2.** The in vitro development of 3D TECs combining sheep brain hemispheres' dECM and human glioblastoma U251 cells. The images demonstrate the sequential stages of the experimental tumour growth in vitro: 3 days **(A, B)**, 21 days **(C, D)**, and 28 days **(E, F)**. Staining by Masson's trichrome method shows cells in red colour tones and collagen matrix in blue colours. The images represent the screenshots obtained using the Aperio ScanScope XT scanner. The left column images **(A, C, E)** are taken at a lower magnification, the scale bards are 100 μm. The areas outlined by yellow frames in the left column images are shown at a higher magnification in the left column **(B, D, F)** with the scale bars of 20 μm. Note the efficient adhesion of the cells to the ECM of the scaffolds, the progressive tumour cells' invasion towards the deeper parts of the scaffold, the remodelling of the ECM and the formation of the cellular network, as well as the preservation of the highly porous structure of the 3D TECs even at the late stages of culturing.

Live/dead staining of 3D TECs showed high viability of U251 cells (Figure A3, A), while the staining of the f-actin cytoskeleton and nuclei of the cells allowed to demonstrate the formation of the cellular network penetrating throughout the scaffolds (Figure A3, B).

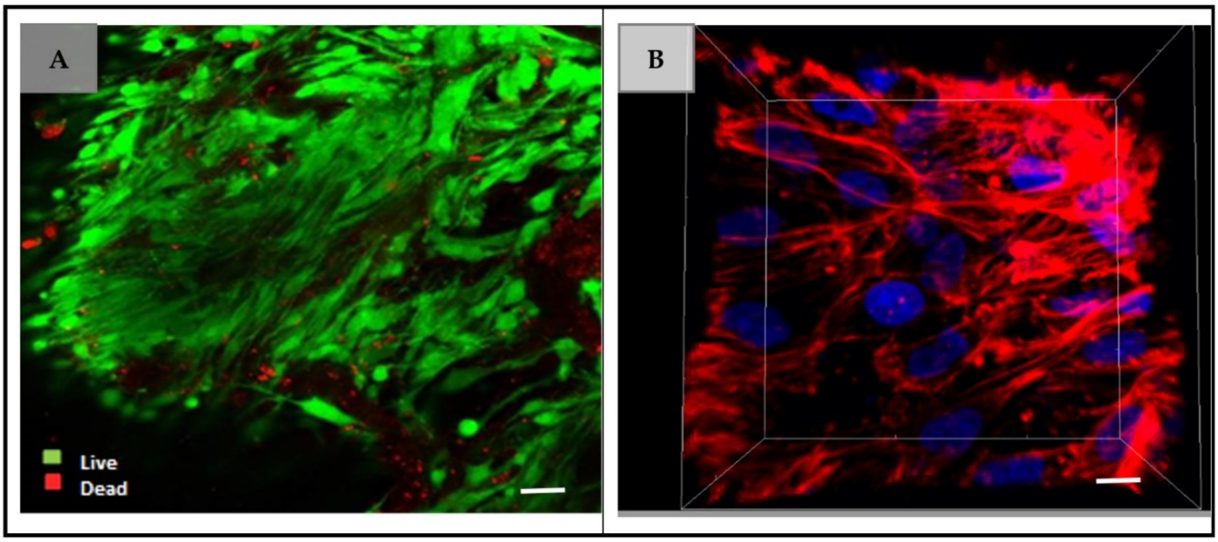


**Figure A3. (A)** Live/dead staining of 3D TECs combining sheep brain dECM scaffolds and human U251 glioblastoma cells cultured in vitro for 28 days. Green colour show live cells, red colour depicts dead cells. Scale bar 20 µm. (B) 3D reconstruction obtained from a confocal z-stack that reveals the spatial organization of the U251 cellular network within the 3D TECs cultured in vitro for 21 days. Cells were stained with phalloidin to show the f-actin cytoskeleton (red) and with DAPI to demonstrate cells' nuclei (blue). Scale bar 10 µm.

Immunohistochemical staining showed that both in 2D culture and in 3D TECs, U251 cells preserved the GFAP-positive phenotype (Figure A4).

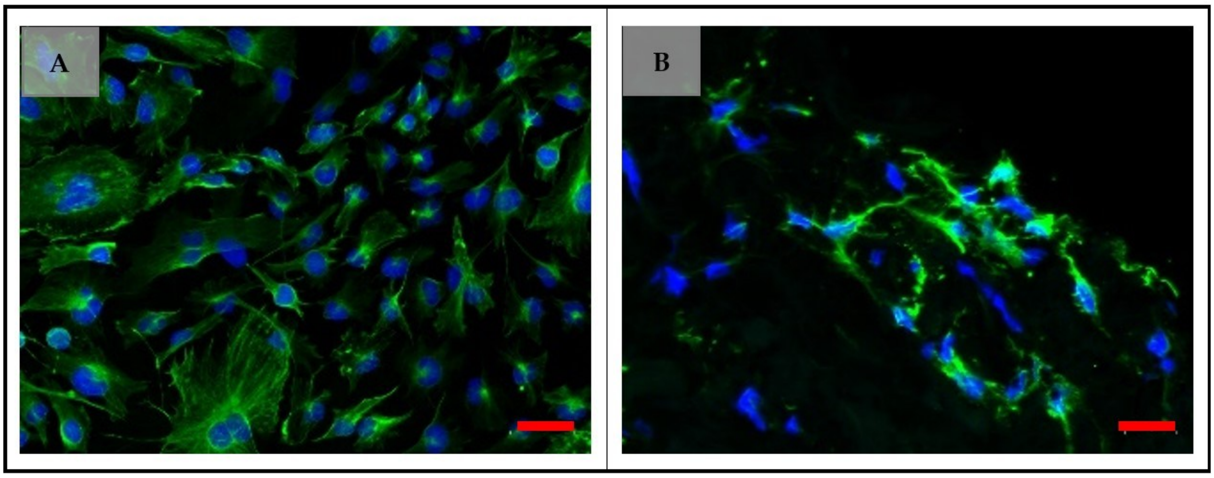


**Figure A4.** Immunohistochemical phenotype of the U251 cells cultured in a conventional 2D monolayer **(A)** and in 3D TECs based on sheep brain dECM **(B)**. Blue colour depicts cell nuclei stained with DAPI, and green colour shows GFAP, stained with anti-GFAP primary antibody and contrasted with Alexa Fluor488-conjugated secondary antibody. Note the complex cell shapes and the formation of the network of GFAP+ cells in 3D TECs. Scale bars 20 µm.

Scanning electron microscopy (SEM) was used to examine the spatial features of the sheep brain dECM fibro-porous scaffolds and U251 cells cultured on them to form the 3D TECs (Figures A5 – A). SEM study revealed that the fibro-porous scaffolds obtained via the decellularisation of sheep brain hemispheres contain 2 compartments with a notably different spatial structure. The first one is formed by membrane-like sheets of flattened fiber bundles, while the second one shows a more random fibrosus structure with the areas of high porosity interchanging with the more parallel aligned bundles of fibrillar elements (Figure A5). Such spatial heterogeneity is assumed to contribute to the enhancement of the heterogeneity of the cells' phenotypes.

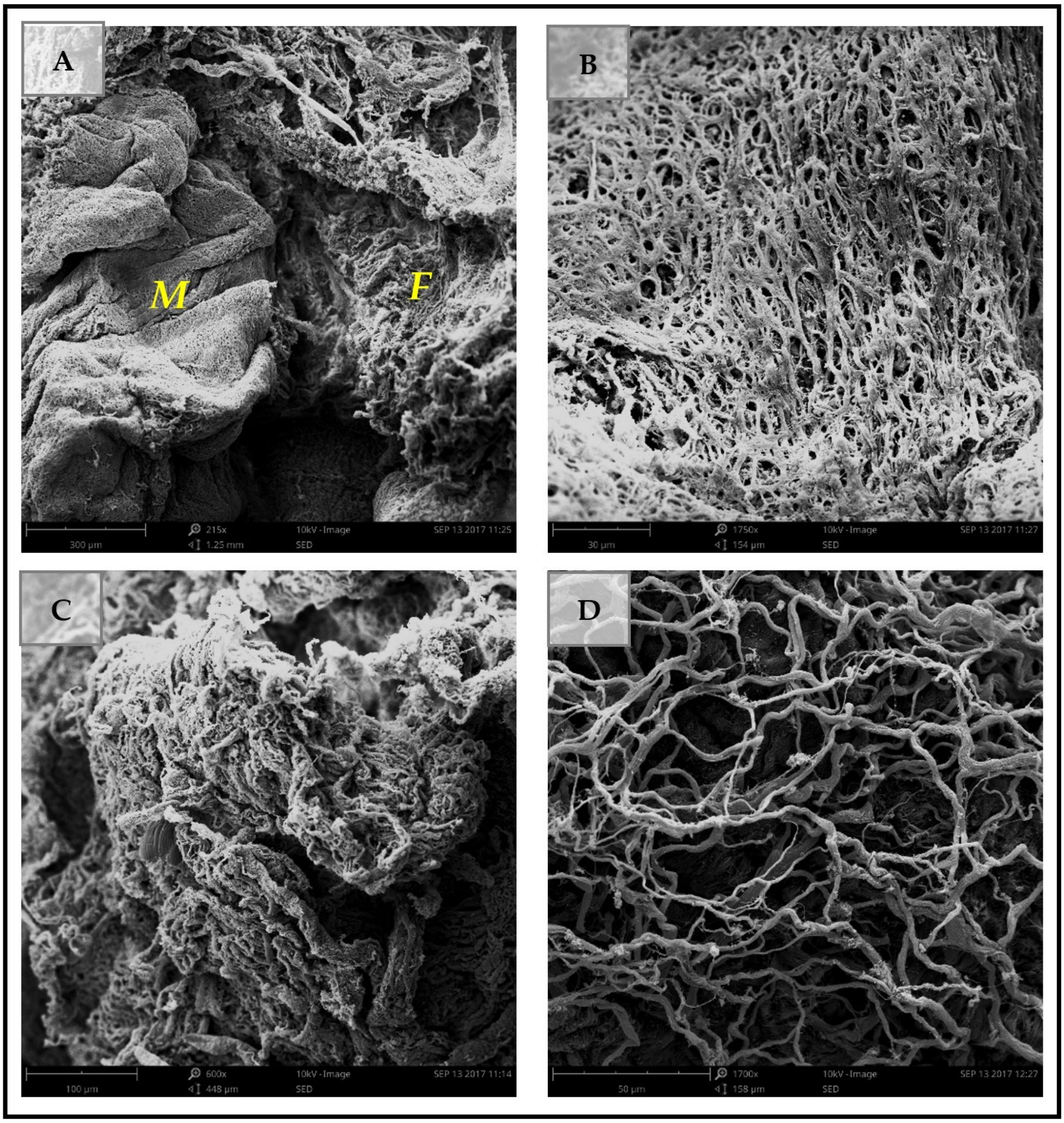


**Figure A5.** SEM images showing the features of the unseeded sheep brain dECM scaffolds. **(A)** An overview of the scaffold surface revealing the presence of two structurally different compartments, the membrane-like one (labelled *"M"*) and the fibrous one (labelled *"F"*). Magnification ×215; scale bar 100 µm. **(B)** The closer view of the membrane-like compartment of the brain dECM scaffold. Magnification ×1750; scale bar 30 µm. **(C, D)** The closer view of the fibrous compartment of the brain dECM scaffold. Magnification ×600 **(C)** and ×1700 **(D)**; scale bar 100 µm **(C)** and 50 µm **(D)**.

SEM imaging confirmed good attachment of the U251 cells to the cell culture plastic, forming a monolayer (Figure A6, A, B) and to the brain dECM scaffolds (Figure A6, C, D).

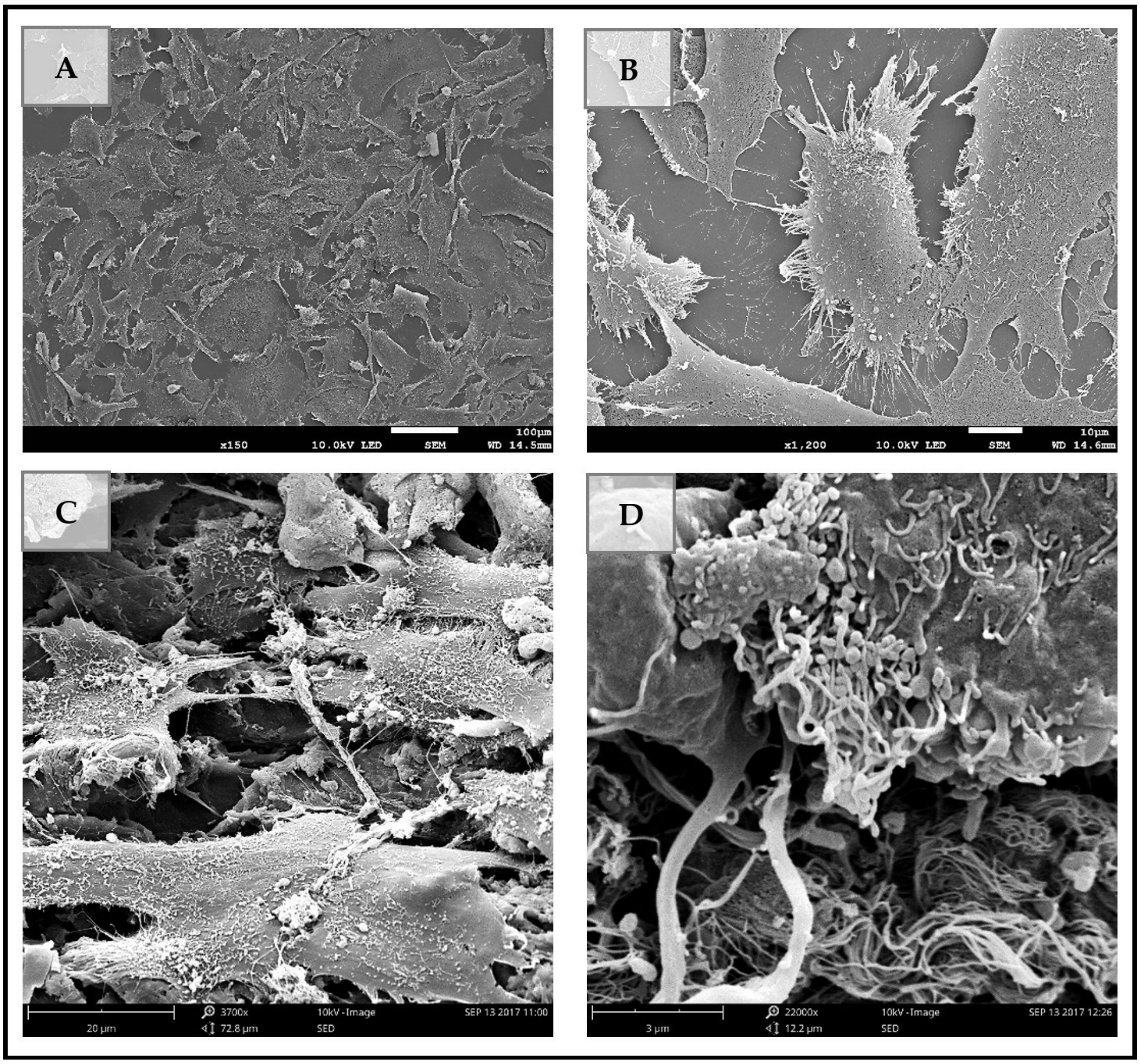


**Figure A6.** SEM imaging of U251 cells cultured for 1 day on the standard plastic vehicles (2D) **(A, B)** and on the 3D sheep brain dECM scaffolds **(C, D)**. The cellular monolayer is visible in 2D culture (A), while in 3D TECs the lining comprising 1-2 rows of cells is formed on the surface of the scaffold **(C)**. Note the wider spread of the cells in 3D TECs **(C)**, compared to 2D culture **(B)**. The shape of the cells is similar across 2D and 3D cultures, and the microvilli on the cell membrane surfaces are visible in both 2D and 3D TEC conditions. In both cell culture platforms, the cells demonstrate cell-cell contacts and communication via filopodia. Magnification ×150 **(A)**, ×1200 **(B)**, ×3700 **(C)**, and ×22000 **(D)**. Scale bars 100 µm **(A)**, 10 µm **(B)**, 20 µm **(C)** and 3 µm **(D)**.

Further stages of the development of the 3D TECs are shown in Figure A7.

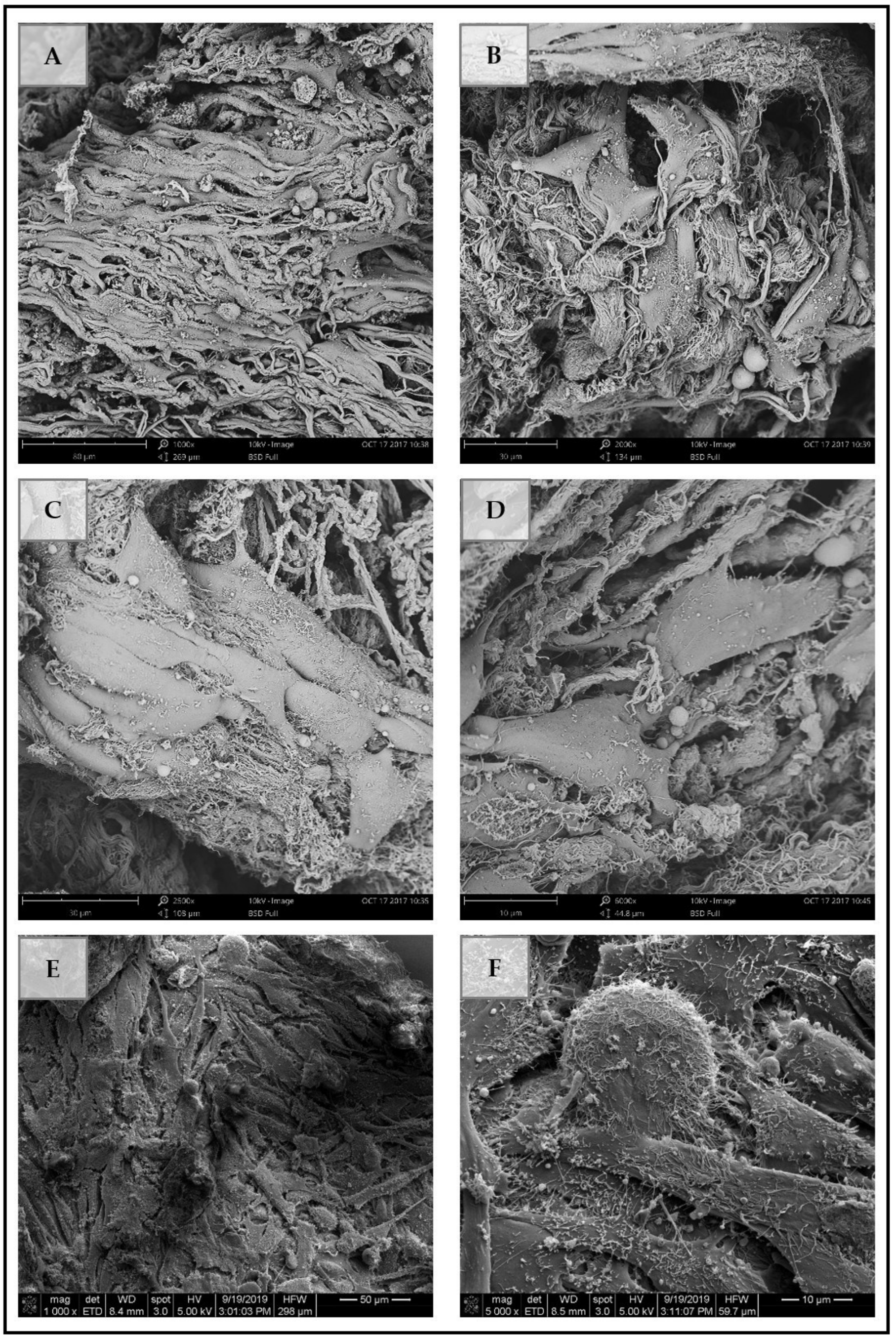


**Figure A7.** SEM imaging of 3D TECs combining human glioblastoma cells U251 and sheep brain dECM fibro-porous scaffolds cultured in vitro for 7 (A, B, C, D) or 30 (E, F) days. Note the alignment and elongation of the cells along with the fiber bundles of the scaffold (A, C). In the areas of the scaffold where the fibrillar elements are arranges more randomly, the cells preserve more polygonal shapes (B, D) and demonstrate invasive behavior (D). At this time point (7 days in vitro), the surfaces of the cells contain only a few microvilli and look smooth. Images (E and F) show the dense layer of the cells on the surface of the TECs, indicating strong cell-ECM and cell-cell contacts. There is notable heterogeneity of cell shapes and sizes, while the spherical cells are rare. The cell surfaces are covered by multiple microvilli, while no bubbling is found, confirming high viability of the cells. Magnification ×1000 **(A)**, ×2000 **(B)**, ×2500 **(C)**, ×5000 **(D, E, F)**. Scale bars 80 µm **(A)**, 30 µm **(B, C)**, 10 µm **(D)** 50 µm **(E),** and 10 µm **(F)**.

Figure A8 provides the overview of the alternative 3D ECM scaffold-based culture platform, the Matrigel-based constructs. In the SEM images, the difference between the Matrigel scaffolds (Figure A8, A-C) and the brain dECM scaffolds shown in Figures A5-A7 above is clearly visible. Matrigel has isotropic spatial organization. It is formed by fine fibers that form mesh-like structure. In contrast to the brain dECM, the adhesion of the U251 cells to the Matrigel was quite weak as it follows from the shapes of the cells and bubbling on their surfaces (Figure A8, D-I). Low adhesion results in detachment of the cells located on the surface of the gel scaffold (see Figure A8, C). The cells located near the surface of the Matrigel plugs have spherical shapes, indicating lack of adhesion to the ECM. Depending on the extent of the embedding into the Matrigel or exposure outside of the scaffold, the cells show abundant vesicles on their membranes, which is commonly considered as a sign of apoptosis. In addition, the formation of cellular clusters was observed (see Figure A8, I), however, within the clusters cells also kept the spherical shapes.

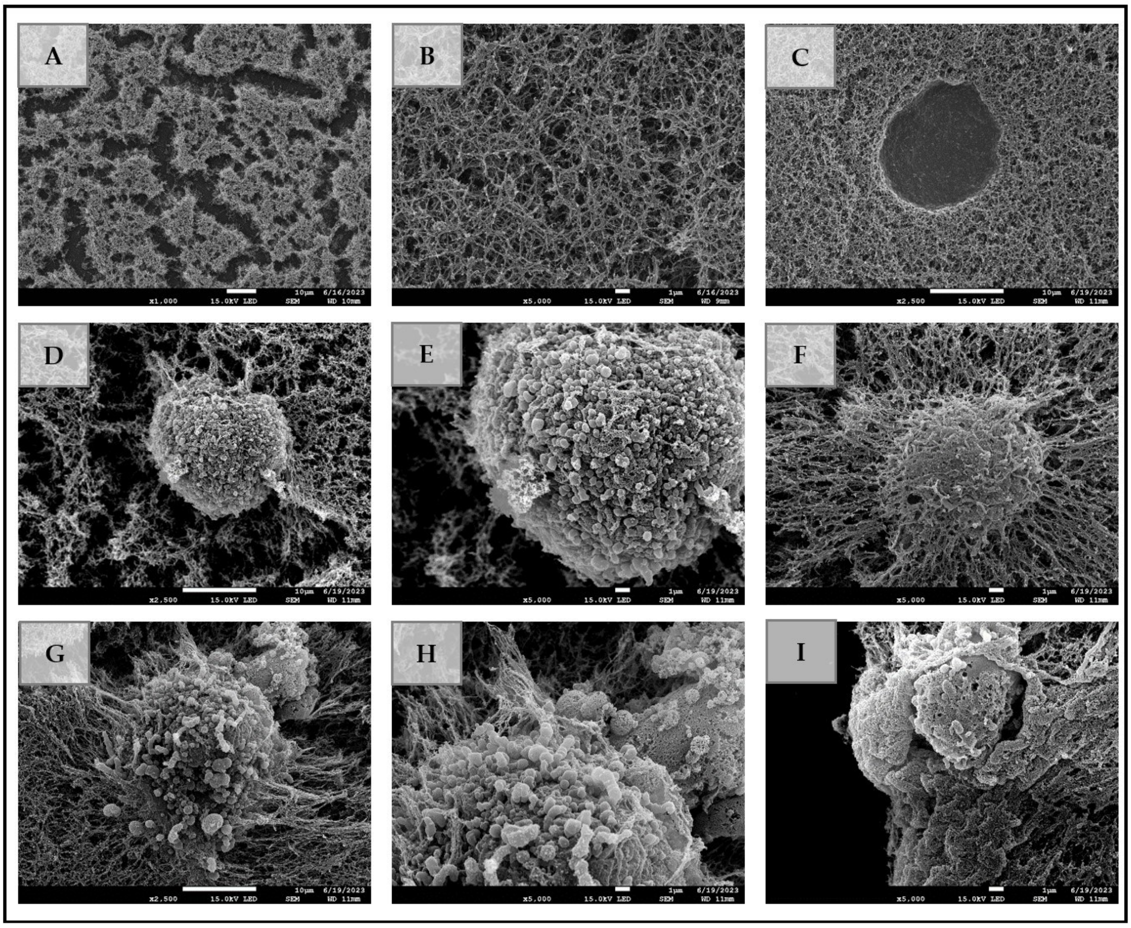


**Figure A8.** SEM imaging of Matrigel and Matrigel-based 3D constructs containing U251 cells. (A, B) View of the unseeded Matrigel scaffold at different magnifications. Note the isotropic structure of the gel. (C) Matrigel-based 3D construct, 1 day in vitro. Note the depression in the gel surface left following the detachment of the individual cell due to low adhesion. (D,F) Matrigel-based construct, 7 days in vitro. Note that the cells located on the surface of the gel have spherical shape and demonstrate multiple bubbling vesicles on the surface. (F-I) Matrigel-based constrcucts, 7 days in vitro. Note that the cells embedded into the gel (F) have less surface stress features than the cells exposed to a greater extent (G, H). (I) Formation of multicellular clusters in Matrigel-based 3D construct. Cells have limited cell-cell communication, filopodia and lamellipodia are absent, indicating the lack of cell motility and invasiveness. Magnification ×1000 **(A)**, ×5000 **(B, E, F, H, I)**, and ×2500 **(C, D, G)**. Scale bars 10 μm **(A, C, D, G)**, and 1 μm **(B, E, F, H, I)**.
