## Appendix B for "What Can We Count On? Performance of Microplate Cell Counting Assays in 2D Monolayer and 3D ECM-based In Vitro Tumour Models"

***Table B1. Calibration of DNA assay for experiments with 2D cell cultures.***

| DNA Concentration, ng | Fluorescence intensity, a.u. | | | | |
|---|---|---|---|---|---|
| | Run 1 | Run 2 | Run 3 | Mean | St. Dev. |
| 0.2 | 2945 | 2919 | 2936 | 2933.333 | 10.78064 |
| 1 | 5832 | 5768 | 5598 | 5732.667 | 98.74321 |
| 10 | 11492 | 11208 | 11181 | 11293.67 | 140.6754 |
| 20 | 22384 | 22857 | 22488 | 22576.33 | 202.9521 |
| 40 | 28361 | 28072 | 28613 | 28348.67 | 221.0344 |
| 80 | 137697 | 137315 | 137671 | 137561 | 174.2718 |
| 100 | 229354 | 232260 | 229403 | 230339 | 1358.499 |

* Gain: 812.

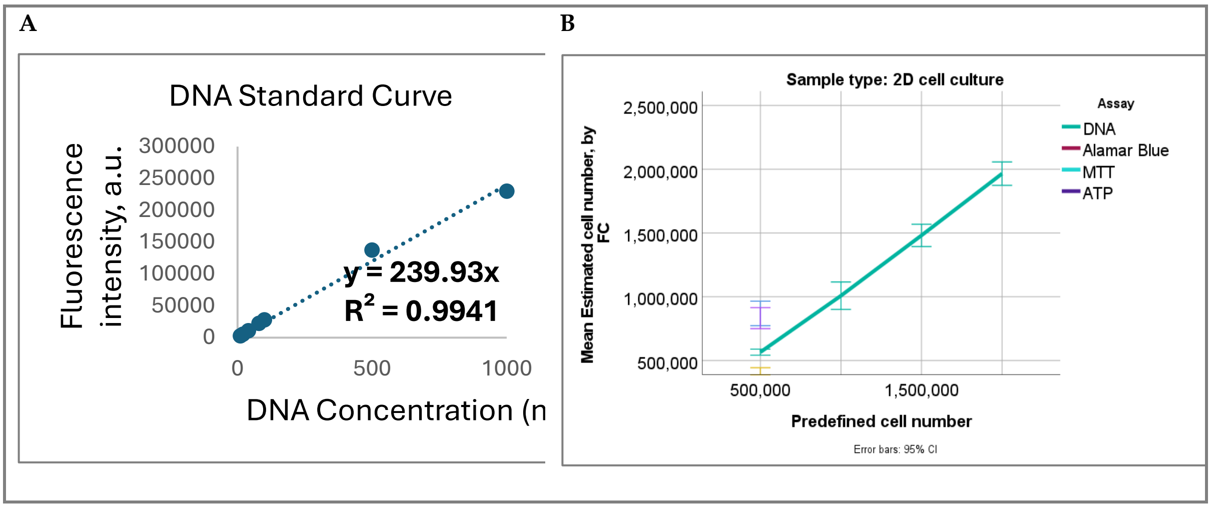


**Figure B1. (a)** Calibration curve for DNA assay for the experiments with 2D cell cultures. **(b)** Overview of the cell number estimation by DNA assay in 2D "static" (1-day) cell culture across the full range of the tested predefined cell numbers. The values of the estimated cell numbers provided for the 500,000 directly counted cells by other examined cell counting assays are shown for comparison.

**Table B2. Fold changes of the readings of the Alamar Blue, ATP and MTT assays in static (one-day) 2D and 3D Matrigel-based cell cultures of U251 cells.**

| Number of cells, counted directly | | Alamar Blue | | MTT | | ATP | |
|---|---|---|---|---|---|---|---|
| N | FC[a] | 2D | 3D | 2D | 3D | 2D | 3D |
| 5,200 | 1 | 1.00±0.43 | 1.00±0.79 | 1.00±0.31 | 0.55±1.30 | 1.00±0.06 | 1.00±0.57 |
| 15,600 | 3 | 4.80±2.26 | 1.55±1.12 | 4.82±1.26 | 2.20±0.93 | 2.77±0.19 | 2.23±0.36 |
| 31,000 | 5.96 | 11.54±3.66 | 2.20±0.53 | 9.24±2.35 | 6.73±0.94 | 6.15±0.19 | 4.79±0.11 |
| 62,000 | 11.92 | 23.00±3.78 | 3.86±0.73 | 20.19±5.57 | 10.83±1.99 | 11.63±0.38 | 12.65±1.99 |
| 125,000 | 24.04 | 41.68±5.19 | 6.86±0.85 | 42.83±9.42 | 20.33±1.89 | 18.03±5.53 | 26.40±6.95 |
| 187,000 | 35.96 | 62.35±8.88 | 10.88±3.29 | 62.49±11.03 | 27.62±2.56 | 27.74±8.46 | 38.12±13.72 |
| 250,000 | 48.08 | 85.10±11.98 | 16.22±1.35 | 84.88±15.89 | 34.08±1.49 | 36.67±11.20 | 52.97±19.49 |
| 375,000 | 72.12 | 121.53±13.82 | 25.29±2.87 | 123.15±19.24 | 42.1167±3.79 | 55.00±16.88 | 82.04±28.64 |
| 500,000 | 96.15 | 167.21±17.63 | 22.60±16.24 | 160.10±24.88 | 48.1167±3.46 | 73.27±18.95 | 113.32±30.67 |

Abbreviation: FC – fold change vs the sample with the smallest number of cells ($5.2\times10^3$). The data presented as Mean ± St. Dev.

As it follows from Table B2 (Appendix B), in the given dilution series, the numbers of the directly counted cells increased up to approximately 96-fold – from the sample with the smallest number of cells (5,200) to the largest samples (500,000 cells). However, in contrast to the DNA assay, metabolic assays' readings were not mirroring the increase in cell numbers in 2D cell culture. For example, while the predefined cell count increased by 96-fold, the Alamar Blue assay readings increased by approximately 167-fold, the MTT assay readings by about 160-fold, reflecting overestimation of the cell numbers in high cell density samples, while the ATP assay readings increased by around 73-fold (underestimation at high cell density).

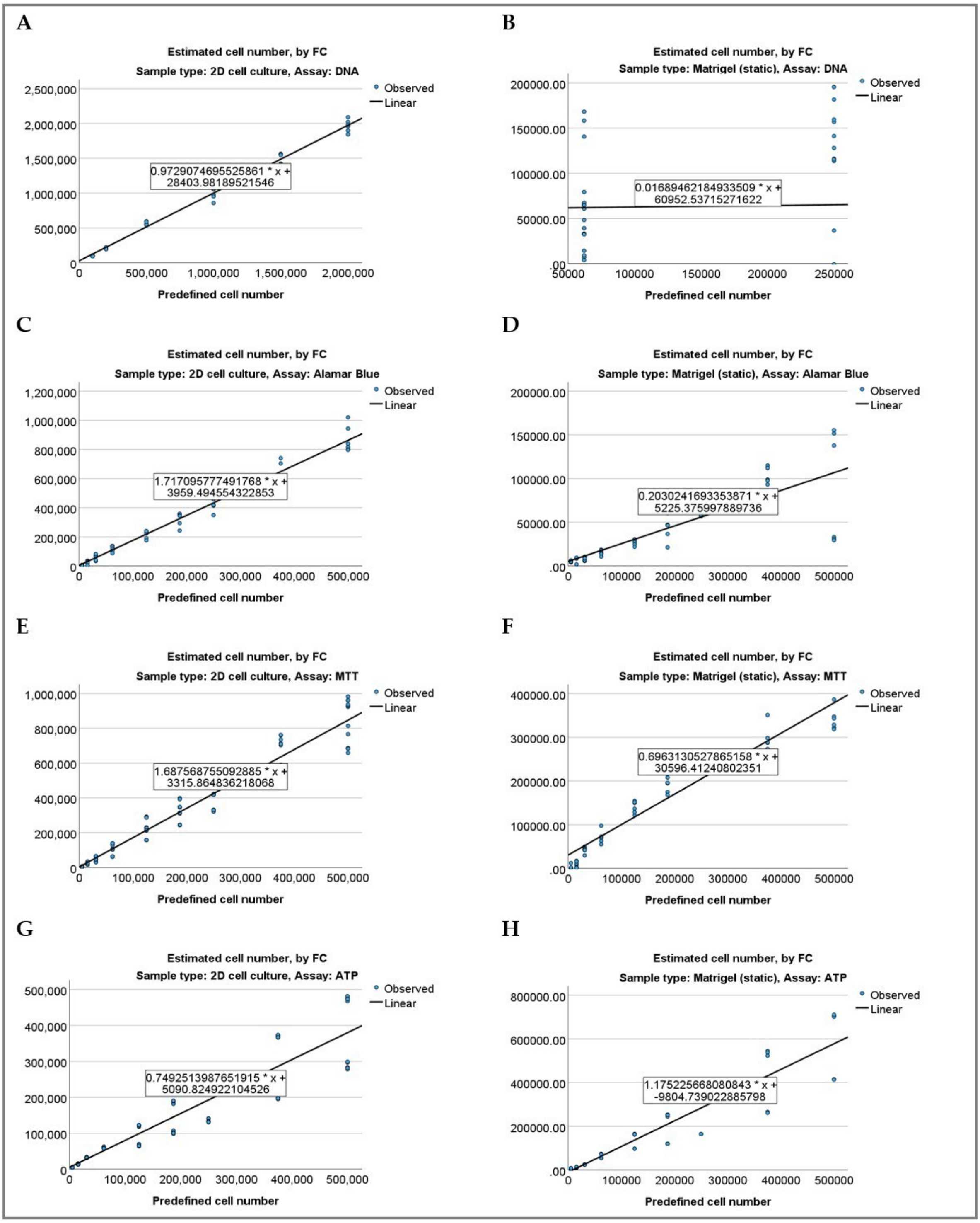


**Figure B2.** Linear regression plots for the assays testing in static 2D **(A, C, E, G)** and 3D Matrigel-based **(B, D, F, H)** cultures of U251 cells. Blue dots show the data point for the estimated cell numbers calculated by the fold change (FC) of the assay readings normalized to the reading in the smallest cell density sample in the series (shown on the vertical axis), while the predefined (directly counted) cell number stands for the independent variable (shown on the horizontal axis). The respective linear regression equations are shown in the inserts on the regression lines. **(A, B)** DNA assay; **(C, D)** Alamar Blue assay; **(E, F)** MTT assay; **(G, H)** ATP assay.

**Table B3. Pairwise comparison of the proportionality indexes of the Alamar Blue, ATP and MTT assays in static (one-day) 2D and 3D Matrigel-based cell cultures of U251 cells via the independent-samples Kruskal-Wallis test.**

| Cell culture platform | Sample 1-Sample 2 | Test Statistic | Std. Error | Std. Test Statistic | Sig. | Adj. Sig. [a] |
|---|---|---|---|---|---|---|
| 2D | ATP-DNA | 5.833 | 5.091 | 1.146 | .252 | 1.000 |
| | ATP-MTT | 16.944 | 4.554 | 3.721 | <.001 | .001 |
| | ATP-Alamar Blue | 19.500 | 4.554 | 4.282 | <.001 | .000 |
| | DNA-MTT | -11.111 | 5.091 | -2.182 | .029 | .174 |
| | DNA-Alamar Blue | -13.667 | 5.091 | -2.684 | .007 | .044 |
| | MTT-Alamar Blue | 2.556 | 4.554 | .561 | .575 | 1.000 |
| 3D Matrigel (static) | DNA-MTT | -5.361 | 6.646 | -.807 | .420 | 1.000 |
| | Alamar Blue-DNA | 7.306 | 6.646 | 1.099 | .272 | 1.000 |
| | Alamar Blue-MTT | -12.667 | 4.008 | -3.160 | .002 | .009 |
| | Alamar Blue-ATP | -14.889 | 4.008 | -3.715 | <.001 | .001 |
| | DNA-ATP | -7.583 | 6.646 | -1.141 | .254 | 1.000 |
| | MTT-ATP | -2.222 | 4.008 | -.554 | .579 | 1.000 |

Each row tests the null hypothesis that the Sample 1 and Sample 2 distributions are the same. Asymptotic significances (2-sided tests) are displayed. The significance level is .050. [a.] Significance values have been adjusted by the Bonferroni correction for multiple tests.

**Table B4. Technical (CVt), biological (CVb) and overall coefficients of variation (CV) of the DNA, Alamar Blue, ATP and MTT assays in static (one-day) 2D and 3D Matrigel-based cell cultures of U251 cells.**

| Cell culture platform | Assay | | Mean | N | Std. Deviation | Min. | Max. |
|---|---|---|---|---|---|---|---|
| 2D | DNA | CVt, % | 3.01 | 6 | 1.44 | 0.88 | 4.80 |
| | | CVb, % | 4.06 | 6 | 2.18 | 2.11 | 7.98 |
| | | CV, % | 5.88 | 6 | 2.21 | 3.98 | 10.18 |
| | Alamar Blue | CVt, % | 12.21 | 8 | 9.20 | 4.51 | 29.71 |
| | | CVb, % | 11.67 | 8 | 10.08 | 2.34 | 34.92 |
| | | CV, % | 22.31 | 9 | 14.35 | 10.54 | 47.05 |
| | MTT | CVt, % | 13.14 | 9 | 6.29 | 4.93 | 22.65 |
| | | CVb, % | 16.02 | 9 | 2.19 | 14.01 | 20.17 |
| | | CV, % | 22.17 | 9 | 5.57 | 15.54 | 30.68 |
| | ATP | CVt, % | 2.57 | 9 | .96 | 1.65 | 4.61 |
| | | CVb, % | 16.92 | 9 | 13.68 | 0.11 | 29.35 |
| | | CV, % | 18.59 | 9 | 13.26 | 3.02 | 30.70 |
| 3D Matrigel (static) | DNA | CVt, % | 59.50 | 2 | 27.93 | 39.75 | 79.25 |
| | | CVb, % | 21.96 | 2 | 3.27 | 19.64 | 24.27 |
| | | CV, % | 58.80 | 2 | 36.23 | 33.18 | 84.42 |
| | Alamar Blue | CVt, % | 12.59 | 9 | 9.84 | 4.92 | 35.08 |
| | | CVb, % | 17.25 | 9 | 19.01 | 3.65 | 65.30 |
| | | CV, % | 26.51 | 9 | 19.95 | 8.31 | 71.84 |
| | MTT | CVt, % | 21.36 | 9 | 31.80 | 2.67 | 98.82 |
| | | CVb, % | 10.98 | 8 | 19.31 | 0.00 | 58.07 |
| | | CV, % | 29.79 | 9 | 39.79 | 4.48 | 121.03 |
| | ATP | CVt, % | 1.33 | 9 | 1.64 | 0.30 | 5.62 |
| | | CVb, % | 26.04 | 9 | 13.82 | 2.02 | 47.05 |
| | | CV, % | 28.05 | 9 | 15.80 | 2.24 | 57.16 |

**Table B5. Pairwise comparison of the technical coefficients of variation (CVt) of the DNA, Alamar Blue, ATP and MTT assays in static (one-day) 2D and 3D Matrigel-based cell cultures of U251 cells via the independent-samples Kruskal-Wallis test. Statistically significant comparisons are highlighted.**

| Cell culture platform | Sample 1-Sample 2 | Test Statistic | Std. Error | Std. Test Statistic | Sig. | Adj. Sig. [a] |
|---|---|---|---|---|---|---|
| 2D | ATP-DNA | 2.000 | 4.944 | .405 | .686 | 1.000 |
| | ATP-Alamar Blue | 15.667 | 4.558 | 3.437 | <.001 | .004 |
| | ATP-MTT | 17.333 | 4.422 | 3.920 | <.001 | .001 |
| | DNA-Alamar Blue | -13.667 | 5.066 | -2.698 | .007 | .042 |
| | DNA-MTT | -15.333 | 4.944 | -3.101 | .002 | .012 |
| | Alamar Blue-MTT | -1.667 | 4.558 | -.366 | .715 | 1.000 |
| 3D Matrigel (static) | ATP-DNA | 21.444 | 6.656 | 3.222 | .001 | .008 |
| | ATP-Alamar Blue | 13.000 | 4.014 | 3.239 | .001 | .007 |
| | ATP-MTT | 12.667 | 4.014 | 3.156 | .002 | .010 |
| | MTT-Alamar Blue | .333 | 4.014 | .083 | .934 | 1.000 |
| | MTT-DNA | 8.778 | 6.656 | 1.319 | .187 | 1.000 |
| | Alamar Blue-DNA | 8.444 | 6.656 | 1.269 | .205 | 1.000 |

Each row tests the null hypothesis that the Sample 1 and Sample 2 distributions are the same. Asymptotic significances (2-sided tests) are displayed. The significance level is .050. [a.] Significance values have been adjusted by the Bonferroni correction for multiple tests.

**Table B6. Pairwise comparison of the biological coefficients of variation (CVb) of the DNA, Alamar Blue, ATP and MTT assays in static (one-day) 2D and 3D Matrigel-based cell cultures of U251 cells via the independent-samples Kruskal-Wallis test. Statistically significant comparisons are highlighted.**

| Cell culture platform | Sample 1-Sample 2 | Test Statistic | Std. Error | Std. Test Statistic | Sig. | Adj. Sig. [a] |
|---|---|---|---|---|---|---|
| 2D | DNA-Alamar Blue | -7.375 | 5.066 | -1.456 | .145 | .873 |
| | DNA-ATP | -10.944 | 4.944 | -2.214 | .027 | .161 |
| | DNA-MTT | -14.500 | 4.944 | -2.933 | .003 | .020 |
| | Alamar Blue-ATP | -3.569 | 4.558 | -.783 | .434 | 1.000 |
| | Alamar Blue-MTT | -7.125 | 4.558 | -1.563 | .118 | .708 |
| | ATP-MTT | 3.556 | 4.422 | .804 | .421 | 1.000 |
| 3D Matrigel (static) | DNA-ATP | -.333 | 6.430 | -.052 | .959 | 1.000 |
| | Alamar Blue-ATP | -5.278 | 3.877 | -1.361 | .173 | 1.000 |
| | MTT-Alamar Blue | 5.618 | 3.997 | 1.406 | .160 | .959 |
| | MTT-DNA | 10.563 | 6.502 | 1.624 | .104 | .626 |
| | MTT-ATP | -10.896 | 3.997 | -2.726 | .006 | .038 |
| | Alamar Blue-DNA | 4.944 | 6.430 | .769 | .442 | 1.000 |

Each row tests the null hypothesis that the Sample 1 and Sample 2 distributions are the same. Asymptotic significances (2-sided tests) are displayed. The significance level is .050. [a.] Significance values have been adjusted by the Bonferroni correction for multiple tests.

**Table B7. Pairwise comparison of the overall coefficients of variation (CV) of the DNA, Alamar Blue, ATP and MTT assays in static (one-day) 2D and 3D Matrigel-based cell cultures of U251 cells via the independent-samples Kruskal-Wallis test. Statistically significant comparisons are highlighted.**

| Cell culture platform | Sample 1-Sample 2 | Test Statistic | Std. Error | Std. Test Statistic | Sig. | Adj. Sig. [a] |
|---|---|---|---|---|---|---|
| 2D | DNA-ATP | -11.333 | 5.096 | -2.224 | .026 | .157 |
| | DNA-Alamar Blue | -14.056 | 5.096 | -2.758 | .006 | .035 |
| | DNA-MTT | -15.556 | 5.096 | -3.053 | .002 | .014 |
| | ATP-Alamar Blue | 2.722 | 4.558 | .597 | .550 | 1.000 |
| | ATP-MTT | 4.222 | 4.558 | .926 | .354 | 1.000 |
| | Alamar Blue-MTT | -1.500 | 4.558 | -.329 | .742 | 1.000 |
| 3D Matrigel (static) | MTT-Alamar Blue | 3.333 | 4.014 | .830 | .406 | 1.000 |
| | MTT-ATP | -5.000 | 4.014 | -1.246 | .213 | 1.000 |
| | MTT-DNA | 12.444 | 6.656 | 1.870 | .062 | .369 |
| | Alamar Blue-ATP | -1.667 | 4.014 | -.415 | .678 | 1.000 |
| | Alamar Blue-DNA | 9.111 | 6.656 | 1.369 | .171 | 1.000 |
| | ATP-DNA | 7.444 | 6.656 | 1.118 | .263 | 1.000 |

Each row tests the null hypothesis that the Sample 1 and Sample 2 distributions are the same. Asymptotic significances (2-sided tests) are displayed. The significance level is .050. [a.] Significance values have been adjusted by the Bonferroni correction for multiple tests.

**Table B8. Hypothesis test summary on the differences of the technical, biological and overall coefficients of variation (CVt, CVb, and CV) of the DNA, Alamar Blue, ATP and MTT assays between the static (one-day) 2D and 3D Matrigel-based cell cultures of U251 cells assessed via the independent-samples Kruskal-Wallis test. Statistically significant comparisons are highlighted.**

| Assay | | Null Hypothesis | Test | Sig. [a,b] |
|---|---|---|---|---|
| DNA | 1 | The distribution of CVt, % is the same across categories of Cell culture platform. | Independent-Samples Kruskal-Wallis Test | .046 |
| | 2 | The distribution of CVb, % is the same across categories of Cell culture platform. | Independent-Samples Kruskal-Wallis Test | .046 |
| | 3 | The distribution of CV, % is the same across categories of Cell culture platform. | Independent-Samples Kruskal-Wallis Test | .046 |
| Alamar Blue | 1 | The distribution of CVt, % is the same across categories of Cell culture platform. | Independent-Samples Kruskal-Wallis Test | .923 |
| | 2 | The distribution of CVb, % is the same across categories of Cell culture platform. | Independent-Samples Kruskal-Wallis Test | .441 |
| | 3 | The distribution of CV, % is the same across categories of Cell culture platform. | Independent-Samples Kruskal-Wallis Test | .757 |
| MTT | 1 | The distribution of CVt, % is the same across categories of Cell culture platform. | Independent-Samples Kruskal-Wallis Test | .508 |
| | 2 | The distribution of CVb, % is the same across categories of Cell culture platform. | Independent-Samples Kruskal-Wallis Test | .009 |
| | 3 | The distribution of CV, % is the same across categories of Cell culture platform. | Independent-Samples Kruskal-Wallis Test | .145 |
| ATP | 1 | The distribution of CVt, % is the same across categories of Cell culture platform. | Independent-Samples Kruskal-Wallis Test | .005 |
| | 2 | The distribution of CVb, % is the same across categories of Cell culture platform. | Independent-Samples Kruskal-Wallis Test | .200 |
| | 3 | The distribution of CV, % is the same across categories of Cell culture platform. | Independent-Samples Kruskal-Wallis Test | .233 |

[a.] The significance level is .050. [b.] Asymptotic significance is displayed.

**Table B9. Analysis of correlations (Spearman's correlation coefficient, Rs) between the coefficients of variation of different types and predefined cell number in the static 2D and 3D (Matrigel-based) cultures of U251 cells. Statistically significant correlations are highlighted.**

| Assay | Metric | Statistic | Cell culture platform | | | | | | | |
|---|---|---|---|---|---|---|---|---|---|---|
| | | | 2D | | | | 3D Matrigel (static) | | | |
| | | | Cell number | CVt, % | CVb, % | CV, % | Cell number | CVt, % | CVb, % | CV, % |
| DNA | Cell number | Rs | 1.000 | -.600 | .371 | .029 | 1.000 | -1.000 | 1.000 | -1.000 |
| | | p | . | .208 | .468 | .957 | . | . | . | . |
| | | N | 6 | 6 | 6 | 6 | 2 | 2 | 2 | 2 |
| | CVt, % | Rs | -.600 | 1.000 | .143 | .371 | -1.000** | 1.000 | -1.000 | 1.000 |
| | | p | .208 | . | .787 | .468 | . | . | . | . |
| | | N | 6 | 6 | 6 | 6 | 2 | 2 | 2 | 2 |
| | CVb, % | Rs | .371 | .143 | 1.000 | .829* | 1.000** | -1.000** | 1.000 | -1.000 |
| | | p | .468 | .787 | . | .042 | . | . | . | . |
| | | N | 6 | 6 | 6 | 6 | 2 | 2 | 2 | 2 |
| | CV, % | Rs | .029 | .371 | .829* | 1.000 | -1.000** | 1.000** | -1.000** | 1.000 |
| | | p | .957 | .468 | .042 | . | . | . | . | . |
| | | N | 6 | 6 | 6 | 6 | 2 | 2 | 2 | 2 |
| Alamar Blue | Cell number | Rs | 1.000 | -.786* | -.786* | -.933** | 1.000 | -.733* | -.150 | -.150 |
| | | p | . | .021 | .021 | <.001 | . | .025 | .700 | .700 |
| | | N | 9 | 8 | 8 | 9 | 9 | 9 | 9 | 9 |
| | CVt, % | Rs | -.786* | 1.000 | .429 | .786* | -.733* | 1.000 | .067 | .367 |
| | | p | .021 | . | .289 | .021 | .025 | . | .865 | .332 |
| | | N | 8 | 8 | 8 | 8 | 9 | 9 | 9 | 9 |
| | CVb, % | Rs | -.786* | .429 | 1.000 | .786* | -.150 | .067 | 1.000 | .800** |
| | | p | .021 | .289 | . | .021 | .700 | .865 | . | .010 |
| | | N | 8 | 8 | 8 | 8 | 9 | 9 | 9 | 9 |
| | CV, % | Rs | -.933** | .786* | .786* | 1.000 | -.150 | .367 | .800** | 1.000 |
| | | p | <.001 | .021 | .021 | . | .700 | .332 | .010 | . |
| | | N | 9 | 8 | 8 | 9 | 9 | 9 | 9 | 9 |
| MTT | Cell number | Rs | 1.000 | -.950** | -.650 | -.933** | 1.000 | -.717* | -.643 | -.933** |
| | | p | . | <.001 | .058 | <.001 | . | .030 | .086 | <.001 |
| | | N | 9 | 9 | 9 | 9 | 9 | 9 | 8 | 9 |
| | CVt, % | Rs | -.950** | 1.000 | .683* | .967** | -.717* | 1.000 | .119 | .850** |
| | | p | <.001 | . | .042 | <.001 | .030 | . | .779 | .004 |
| | | N | 9 | 9 | 9 | 9 | 9 | 9 | 8 | 9 |
| | CVb, % | Rs | -.650 | .683* | 1.000 | .733* | -.643 | .119 | 1.000 | .429 |
| | | p | .058 | .042 | . | .025 | .086 | .779 | . | .289 |
| | | N | 9 | 9 | 9 | 9 | 8 | 8 | 8 | 8 |
| | CV, % | Rs | -.933** | .967** | .733* | 1.000 | -.933** | .850** | .429 | 1.000 |
| | | p | <.001 | <.001 | .025 | . | <.001 | .004 | .289 | . |
| | | N | 9 | 9 | 9 | 9 | 9 | 9 | 8 | 9 |
| ATP | Cell number | Rs | 1.000 | -.717* | .650 | .650 | 1.000 | -.233 | .233 | .200 |
| | | p | . | .030 | .058 | .058 | . | .546 | .546 | .606 |
| | | N | 9 | 9 | 9 | 9 | 9 | 9 | 9 | 9 |
| | CVt, % | Rs | -.717* | 1.000 | -.333 | -.333 | -.233 | 1.000 | .600 | .650 |
| | | p | .030 | . | .381 | .381 | .546 | . | .088 | .058 |

| | | | | | | | | | | |
|---|---|---|---|---|---|---|---|---|---|---|
| | | N | 9 | 9 | 9 | 9 | 9 | 9 | 9 | 9 |
| | CVb, % | Rs | .650 | -.333 | 1.000 | 1.000** | .233 | .600 | 1.000 | .983** |
| | | p | .058 | .381 | . | . | .546 | .088 | . | <.001 |
| | | N | 9 | 9 | 9 | 9 | 9 | 9 | 9 | 9 |
| | CV, % | Rs | .650 | -.333 | 1.000** | 1.000 | .200 | .650 | .983** | 1.000 |
| | | p | .058 | .381 | . | . | .606 | .058 | <.001 | . |
| | | N | 9 | 9 | 9 | 9 | 9 | 9 | 9 | 9 |

* Correlation is significant at the 0.05 level (2-tailed). ** Correlation is significant at the 0.01 level (2-tailed).

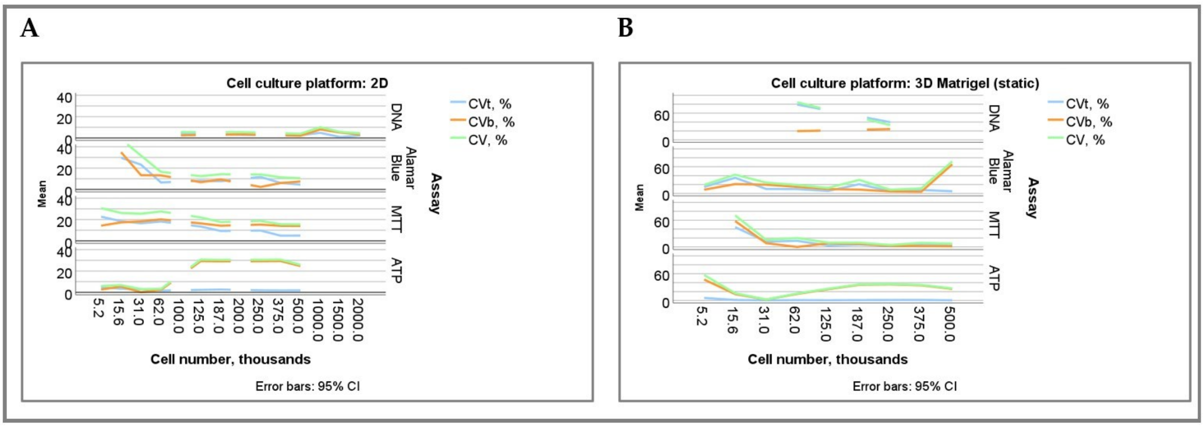


**Figure B3.** Dependence of the technical, biological, and overall coefficients of variation (CVt, CVb, and CV, respectively) of different assays on the predefined cell number in static (1-day) 2D and 3D (Matrigel-bases) cultures of U251 cells. **(A)** 2D cultures; **(B)** 3D Matrigel constructs.

**Table B10. Descriptive statistics for estimated cell number (ECN) detected by different assays in dynamic (1-21 day) 3D Matrigel-based cultures of U251 cells and calculated by fold changes compared to the each assay average reading on day 1.**

| Time, days in vitro | Assay | N | Mean | Std. Deviation | Minimum | Maximum |
|---|---|---|---|---|---|---|
| 1 | DNA | 26 | 10000.00 | 1299.57 | 7633.50 | 12581.30 |
| | Alamar Blue | 9 | 10000.00 | 4799.21 | 519.51 | 16554.46 |
| | MTT | 36 | 10000.00 | 1231.03 | 7725.86 | 11428.48 |
| | ATP | 27 | 10013.92 | 2409.44 | 7872.62 | 14550.78 |
| 3 | DNA | 27 | 10756.96 | 3410.10 | 7172.10 | 16706.40 |
| | Alamar Blue | 27 | 18213.38 | 3933.53 | 10252.32 | 27228.63 |
| | MTT | 36 | 13178.57 | 3387.77 | 6432.88 | 16600.39 |
| | ATP | 27 | 17670.31 | 2902.30 | 11420.96 | 20824.17 |
| 7 | DNA | 24 | 36709.28 | 10508.88 | 18492.80 | 51173.40 |
| | Alamar Blue | 27 | 85168.85 | 38938.61 | 26466.12 | 133377.86 |
| | MTT | 36 | 33255.65 | 10237.71 | 18187.23 | 43870.48 |
| | ATP | 27 | 43827.04 | 23692.71 | 14801.42 | 68333.28 |
| 10 | DNA | 27 | 75819.69 | 24453.83 | 34818.50 | 119311.80 |
| | Alamar Blue | 27 | 167576.63 | 11872.70 | 142252.81 | 188821.46 |
| | MTT | 36 | 52903.37 | 10038.93 | 36288.93 | 65145.85 |
| | ATP | 24 | 112287.96 | 52856.84 | 15446.41 | 163436.10 |
| 14 | DNA | 27 | 306637.40 | 90536.27 | 147381.80 | 443427.00 |
| | Alamar Blue | 27 | 190681.33 | 25189.57 | 149647.45 | 249706.07 |
| | MTT | 36 | 105381.64 | 18795.11 | 72668.63 | 141784.20 |
| | ATP | 27 | 196758.13 | 128131.96 | 15577.44 | 294440.22 |
| 17 | DNA | 18 | 395541.57 | 34595.61 | 334568.00 | 456038.30 |
| | Alamar Blue | 27 | 221103.58 | 49652.40 | 128413.79 | 307782.04 |
| | MTT | 36 | 147857.28 | 15528.20 | 116159.72 | 171992.88 |
| | ATP | 27 | 255890.97 | 118261.32 | 14248.19 | 353065.23 |
| 21 | DNA | 27 | 502760.09 | 86388.81 | 381195.50 | 691782.90 |
| | Alamar Blue | 27 | 233103.66 | 45518.30 | 197765.29 | 377317.51 |
| | MTT | 36 | 217220.99 | 24261.91 | 171287.61 | 250629.46 |
| | ATP | 27 | 519525.59 | 329972.89 | 11360.08 | 831837.90 |

**Table B11. Descriptive statistics for estimated cell number (ECN) detected by different assays in dynamic (1-21 day) 3D brain dECM-based TECs with U251 cells and calculated by fold changes compared to the each assay average reading on day 1.**

| Time, days in vitro | Assay | N | Mean | Std. Deviation | Minimum | Maximum |
|---|---|---|---|---|---|---|
| 1 | DNA | 21 | 10000.00 | 5830.13 | 3195.95 | 21715.36 |
| | Alamar Blue | 16 | 10000.00 | 9232.33 | 840.26 | 27748.91 |
| | MTT | 27 | 10000.00 | 8385.74 | 2361.99 | 43450.06 |
| | ATP | 27 | 10000.00 | 4800.73 | 5252.53 | 19010.39 |
| 3 | DNA | 24 | 23094.81 | 6781.44 | 14775.91 | 41927.30 |
| | Alamar Blue | 14 | 39341.16 | 29079.53 | 748.45 | 75797.64 |
| | MTT | 27 | 10914.73 | 6367.83 | 3147.83 | 21222.09 |
| | ATP | 27 | 38453.45 | 10161.94 | 17768.40 | 57232.72 |
| 7 | DNA | 23 | 34738.35 | 10524.10 | 10379.76 | 55284.29 |
| | Alamar Blue | 27 | 110749.34 | 39002.32 | 37529.51 | 157216.33 |
| | MTT | 27 | 33953.48 | 9963.29 | 22007.93 | 50410.33 |
| | ATP | 27 | 114836.49 | 38912.55 | 44952.62 | 173169.89 |
| 10 | DNA | 24 | 63951.84 | 10703.25 | 39981.60 | 83620.20 |
| | Alamar Blue | 27 | 119599.20 | 41065.23 | 48838.90 | 174410.19 |
| | MTT | 27 | 50834.43 | 7144.77 | 39745.40 | 59840.38 |
| | ATP | 27 | 127270.11 | 24574.40 | 92221.26 | 169934.68 |
| 14 | DNA | 24 | 76971.54 | 15770.87 | 49808.63 | 105761.83 |
| | Alamar Blue | 27 | 232795.96 | 43777.31 | 174863.33 | 324074.57 |
| | MTT | 27 | 71004.26 | 11401.28 | 59391.33 | 98683.19 |
| | ATP | 27 | 174638.81 | 29208.73 | 130971.23 | 221210.99 |
| 17 | DNA | 8 | 92264.91 | 9842.55 | 80433.99 | 107407.66 |
| | Alamar Blue | 27 | 256791.90 | 48841.34 | 168863.99 | 343706.45 |
| | MTT | 27 | 102346.27 | 37618.70 | 68709.11 | 186360.18 |
| | ATP | 27 | 219763.48 | 22906.44 | 184659.42 | 256999.84 |
| 21 | DNA | 24 | 96387.44 | 24025.73 | 58838.77 | 152354.37 |
| | Alamar Blue | 27 | 232293.65 | 31341.87 | 181220.08 | 297811.50 |
| | MTT | 27 | 111476.95 | 15242.83 | 80608.93 | 130228.94 |
| | ATP | 27 | 225845.90 | 15983.66 | 196866.76 | 254638.60 |

.

**Table B12. Results of the independent-samples Kruskal-Wallis test for the differences in estimated cell number (ECN) detected by different assays in dynamic 3D culture between the Matrigel-based constructs and the brain dECM-based TECs with U251 cells. The statistically significant differences are highlighted.**

| Time, days in vitro | | Assay | | | |
|---|---|---|---|---|---|
| | | DNA | Alamar Blue | MTT | ATP |
| 1 | Total N | 47 | 25 | 63 | 54 |
| | Test Statistic | 1.145 [a,b] | .157 [a,b] | .563 [a,b] | 1.573 [a,b] |
| | Degree Of Freedom | 1 | 1 | 1 | 1 |
| | Asymptotic Sig.(2-sided test) | .285 | .692 | .453 | .210 |
| 3 | Total N | 51 | 41 | 63 | 54 |
| | Test Statistic | 35.113 [a,b] | 7.112 [a,b] | 2.168 [a,b] | 29.603 [a,b] |
| | Degree Of Freedom | 1 | 1 | 1 | 1 |
| | Asymptotic Sig.(2-sided test) | <.001 | .008 | .141 | <.001 |
| 7 | Total N | 47 | 54 | 63 | 54 |
| | Test Statistic | .799 [a,b] | 6.689 [a,b] | .766 [a,b] | 30.552 [a,b] |
| | Degree Of Freedom | 1 | 1 | 1 | 1 |
| | Asymptotic Sig.(2-sided test) | .371 | .010 | .382 | <.001 |
| 10 | Total N | 51 | 54 | 63 | 51 |
| | Test Statistic | 2.057 [a,b] | 26.846 [a,b] | 1.528 [a,b] | .051 [a,b] |
| | Degree Of Freedom | 1 | 1 | 1 | 1 |
| | Asymptotic Sig.(2-sided test) | .152 | <.001 | .216 | .821 |
| 14 | Total N | 51 | 54 | 63 | 54 |
| | Test Statistic | 37.385 [a,b] | 16.599 [a,b] | 35.177 [a,b] | 4.418 [a,b] |
| | Degree Of Freedom | 1 | 1 | 1 | 1 |
| | Asymptotic Sig.(2-sided test) | <.001 | <.001 | <.001 | .036 |
| 17 | Total N | 26 | 54 | 63 | 54 |
| | Test Statistic | 16.000 [a,b] | 6.869 [a,b] | 21.650 [a,b] | 4.418 [a,b] |
| | Degree Of Freedom | 1 | 1 | 1 | 1 |
| | Asymptotic Sig.(2-sided test) | <.001 | .009 | <.001 | .036 |
| 21 | Total N | 51 | 54 | 63 | 54 |
| | Test Statistic | 37.385 [a,b] | .356 [a,b] | 45.568 [a,b] | 4.418 [a,b] |
| | Degree Of Freedom | 1 | 1 | 1 | 1 |
| | Asymptotic Sig.(2-sided test) | <.001 | .551 | <.001 | .036 |

[a.] The test statistic is adjusted for ties. [b.] Multiple comparisons are not performed because there are less than three test fields.

**Table B13. Ranks of the estimated cell numbers obtained by different assays in 3D dynamic Matrigel-based constructs and bran dECM-based TECs of U251 cells.**

| Time, days in vitro | | Assay | Sample type | | | |
|---|---|---|---|---|---|---|
| | | | Matrigel (dynamic) | | TEC | |
| | | | N | Mean Rank | N | Mean Rank |
| 1 | Estimated cell number, by FC | DNA | 26 | 50.50 | 21 | 47.90 |
| | | Alamar Blue | 9 | 55.22 | 16 | 39.38 |
| | | MTT | 36 | 51.03 | 27 | 42.74 |
| | | ATP | 27 | 44.59 | 27 | 51.70 |
| | | Total | 98 | | 91 | |
| 3 | Estimated cell number, by FC | DNA | 27 | 29.30 | 24 | 45.75 |
| | | Alamar Blue | 27 | 83.56 | 14 | 57.00 |
| | | MTT | 36 | 44.14 | 27 | 19.63 |
| | | ATP | 27 | 83.96 | 27 | 68.59 |
| | | Total | 117 | | 92 | |
| 7 | Estimated cell number, by FC | DNA | 24 | 50.79 | 23 | 27.39 |
| | | Alamar Blue | 27 | 84.52 | 27 | 74.93 |
| | | MTT | 36 | 44.17 | 27 | 26.67 |
| | | ATP | 27 | 54.22 | 27 | 77.30 |
| | | Total | 114 | | 104 | |
| 10 | Estimated cell number, by FC | DNA | 27 | 47.37 | 24 | 39.58 |
| | | Alamar Blue | 27 | 99.00 | 27 | 73.59 |
| | | MTT | 36 | 30.25 | 27 | 18.78 |
| | | ATP | 24 | 63.08 | 27 | 78.56 |
| | | Total | 114 | | 105 | |
| 14 | Estimated cell number, by FC | DNA | 27 | 92.37 | 24 | 28.83 |
| | | Alamar Blue | 27 | 62.63 | 27 | 88.44 |
| | | MTT | 36 | 27.50 | 27 | 23.48 |
| | | ATP | 27 | 64.00 | 27 | 68.56 |
| | | Total | 117 | | 105 | |
| 17 | Estimated cell number, by FC | DNA | 18 | 99.17 | 8 | 20.00 |
| | | Alamar Blue | 27 | 55.33 | 27 | 69.00 |
| | | MTT | 36 | 26.92 | 27 | 17.78 |
| | | ATP | 27 | 60.67 | 27 | 55.63 |
| | | Total | 108 | | 89 | |
| 21 | Estimated cell number, by FC | DNA | 27 | 86.85 | 24 | 20.29 |
| | | Alamar Blue | 27 | 41.26 | 27 | 79.11 |
| | | MTT | 36 | 40.81 | 27 | 31.07 |
| | | ATP | 27 | 73.15 | 27 | 77.89 |
| | | Total | 117 | | 105 | |

**Table B14. Pairwise comparison of the ECN obtained by the DNA, Alamar Blue, ATP and MTT assays in dynamic 3D Matrigel-based cell cultures of U251 cells via the independent-samples Kruskal-Wallis test. Statistically significant comparisons are highlighted.**

| Time, days in vitro | Sample 1-Sample 2 | Test Statistic | Std. Error | Std. Test Statistic | Sig. | Adj. Sig. [a] |
|---|---|---|---|---|---|---|
| 1 | ATP-DNA | 5.907 | 7.812 | .756 | .450 | 1.000 |
| | ATP-MTT | 6.435 | 7.238 | .889 | .374 | 1.000 |
| | ATP-Alamar Blue | 10.630 | 10.943 | .971 | .331 | 1.000 |
| | DNA-MTT | -.528 | 7.318 | -.072 | .943 | 1.000 |
| | DNA-Alamar Blue | -4.722 | 10.996 | -.429 | .668 | 1.000 |
| | MTT-Alamar Blue | 4.194 | 10.596 | .396 | .692 | 1.000 |
| 3 | DNA-MTT | -14.843 | 8.635 | -1.719 | .086 | .514 |
| | DNA-Alamar Blue | -54.259 | 9.231 | -5.878 | <.001 | .000 |
| | MTT-Alamar Blue | 39.417 | 8.635 | 4.565 | <.001 | .000 |
| | DNA-ATP | -54.667 | 9.231 | -5.922 | <.001 | .000 |
| | MTT-ATP | -39.824 | 8.635 | -4.612 | <.001 | .000 |
| | Alamar Blue-ATP | -.407 | 9.231 | -.044 | .965 | 1.000 |
| 7 | ATP-Alamar Blue | 30.296 | 8.996 | 3.368 | <.001 | .005 |
| | DNA-Alamar Blue | -33.727 | 9.273 | -3.637 | <.001 | .002 |
| | MTT-Alamar Blue | 40.352 | 8.415 | 4.795 | <.001 | .000 |
| | DNA-ATP | -3.431 | 9.273 | -.370 | .711 | 1.000 |
| | MTT-ATP | -10.056 | 8.415 | -1.195 | .232 | 1.000 |
| | MTT-DNA | 6.625 | 8.710 | .761 | .447 | 1.000 |
| 10 | ATP-Alamar Blue | 35.917 | 9.273 | 3.873 | <.001 | .001 |
| | DNA-Alamar Blue | -51.630 | 8.996 | -5.739 | <.001 | .000 |
| | MTT-Alamar Blue | 68.750 | 8.415 | 8.170 | <.001 | .000 |
| | DNA-ATP | -15.713 | 9.273 | -1.695 | .090 | .541 |
| | MTT-ATP | -32.833 | 8.710 | -3.770 | <.001 | .001 |
| | MTT-DNA | 17.120 | 8.415 | 2.035 | .042 | .251 |
| 14 | ATP-DNA | 28.370 | 9.232 | 3.073 | .002 | .013 |
| | MTT-Alamar Blue | 35.130 | 8.635 | 4.068 | <.001 | .000 |
| | MTT-ATP | -36.500 | 8.635 | -4.227 | <.001 | .000 |
| | Alamar Blue-ATP | -1.370 | 9.232 | -.148 | .882 | 1.000 |
| | MTT-DNA | 64.870 | 8.635 | 7.512 | <.001 | .000 |
| | Alamar Blue-DNA | 29.741 | 9.232 | 3.222 | .001 | .008 |
| 17 | ATP-DNA | 38.500 | 9.531 | 4.040 | <.001 | .000 |
| | MTT-Alamar Blue | 28.417 | 7.974 | 3.564 | <.001 | .002 |
| | MTT-ATP | -33.750 | 7.974 | -4.233 | <.001 | .000 |
| | Alamar Blue-ATP | -5.333 | 8.524 | -.626 | .532 | 1.000 |
| | MTT-DNA | 72.250 | 9.042 | 7.991 | <.001 | .000 |
| | Alamar Blue-DNA | 43.833 | 9.531 | 4.599 | <.001 | .000 |
| 21 | ATP-DNA | 13.704 | 9.232 | 1.484 | .138 | .826 |
| | MTT-Alamar Blue | .454 | 8.635 | .053 | .958 | 1.000 |
| | MTT-ATP | -32.343 | 8.635 | -3.745 | <.001 | .001 |
| | Alamar Blue-ATP | -31.889 | 9.232 | -3.454 | <.001 | .003 |
| | MTT-DNA | 46.046 | 8.635 | 5.332 | <.001 | .000 |
| | Alamar Blue-DNA | 45.593 | 9.232 | 4.939 | <.001 | .000 |

Each row tests the null hypothesis that the Sample 1 and Sample 2 distributions are the same. Asymptotic significances (2-sided tests) are displayed. The significance level is .050. [a.] Significance values have been adjusted by the Bonferroni correction for multiple tests.

**Table B15. Pairwise comparison of the ECN obtained by the DNA, Alamar Blue, ATP and MTT assays in dynamic 3D TECs of U251 cells via the independent-samples Kruskal-Wallis test. Statistically significant comparisons are highlighted.**

| Time, days in vitro | Sample 1-Sample 2 | Test Statistic | Std. Error | Std. Test Statistic | Sig. | Adj. Sig. [a] |
|---|---|---|---|---|---|---|
| 1 | DNA-ATP | -3.799 | 7.685 | -.494 | .621 | 1.000 |
| | MTT-ATP | -8.963 | 7.189 | -1.247 | .212 | 1.000 |
| | Alamar Blue-ATP | -12.329 | 8.333 | -1.480 | .139 | .834 |
| | MTT-DNA | 5.164 | 7.685 | .672 | .502 | 1.000 |
| | Alamar Blue-DNA | 8.530 | 8.765 | .973 | .330 | 1.000 |
| | Alamar Blue-MTT | -3.366 | 8.333 | -.404 | .686 | 1.000 |
| 3 | DNA-Alamar Blue | -11.250 | 8.980 | -1.253 | .210 | 1.000 |
| | MTT-Alamar Blue | 37.370 | 8.794 | 4.250 | <.001 | .000 |
| | DNA-ATP | -22.843 | 7.491 | -3.049 | .002 | .014 |
| | MTT-ATP | -48.963 | 7.267 | -6.737 | <.001 | .000 |
| | Alamar Blue-ATP | -11.593 | 8.794 | -1.318 | .187 | 1.000 |
| | MTT-DNA | 26.120 | 7.491 | 3.487 | <.001 | .003 |
| 7 | DNA-Alamar Blue | -47.535 | 8.560 | -5.553 | <.001 | .000 |
| | MTT-Alamar Blue | 48.259 | 8.210 | 5.878 | <.001 | .000 |
| | DNA-ATP | -49.905 | 8.560 | -5.830 | <.001 | .000 |
| | MTT-ATP | -50.630 | 8.210 | -6.167 | <.001 | .000 |
| | Alamar Blue-ATP | -2.370 | 8.210 | -.289 | .773 | 1.000 |
| | MTT-DNA | .725 | 8.560 | .085 | .933 | 1.000 |
| 10 | DNA-Alamar Blue | -34.009 | 8.544 | -3.981 | <.001 | .000 |
| | MTT-Alamar Blue | 54.815 | 8.289 | 6.613 | <.001 | .000 |
| | DNA-ATP | -38.972 | 8.544 | -4.561 | <.001 | .000 |
| | MTT-ATP | -59.778 | 8.289 | -7.212 | <.001 | .000 |
| | Alamar Blue-ATP | -4.963 | 8.289 | -.599 | .549 | 1.000 |
| | MTT-DNA | 20.806 | 8.544 | 2.435 | .015 | .089 |
| 14 | ATP-Alamar Blue | 19.889 | 8.289 | 2.400 | .016 | .098 |
| | DNA-Alamar Blue | -59.611 | 8.544 | -6.977 | <.001 | .000 |
| | MTT-Alamar Blue | 64.963 | 8.289 | 7.838 | <.001 | .000 |
| | DNA-ATP | -39.722 | 8.544 | -4.649 | <.001 | .000 |
| | MTT-ATP | -45.074 | 8.289 | -5.438 | <.001 | .000 |
| | MTT-DNA | 5.352 | 8.544 | .626 | .531 | 1.000 |
| 17 | ATP-Alamar Blue | 13.370 | 7.032 | 1.901 | .057 | .343 |
| | DNA-Alamar Blue | -49.000 | 10.400 | -4.712 | <.001 | .000 |
| | MTT-Alamar Blue | 51.222 | 7.032 | 7.285 | <.001 | .000 |
| | DNA-ATP | -35.630 | 10.400 | -3.426 | <.001 | .004 |
| | MTT-ATP | -37.852 | 7.032 | -5.383 | <.001 | .000 |
| | MTT-DNA | 2.222 | 10.400 | .214 | .831 | 1.000 |
| 21 | ATP-Alamar Blue | 1.222 | 8.289 | .147 | .883 | 1.000 |
| | DNA-MTT | -10.782 | 8.544 | -1.262 | .207 | 1.000 |
| | DNA-Alamar Blue | -58.819 | 8.544 | -6.884 | <.001 | .000 |
| | MTT-Alamar Blue | 48.037 | 8.289 | 5.795 | <.001 | .000 |
| | DNA-ATP | -57.597 | 8.544 | -6.741 | <.001 | .000 |
| | MTT-ATP | -46.815 | 8.289 | -5.648 | <.001 | .000 |

Each row tests the null hypothesis that the Sample 1 and Sample 2 distributions are the same. Asymptotic significances (2-sided tests) are displayed. The significance level is .050. [a.] Significance values have been adjusted by the Bonferroni correction for multiple tests.

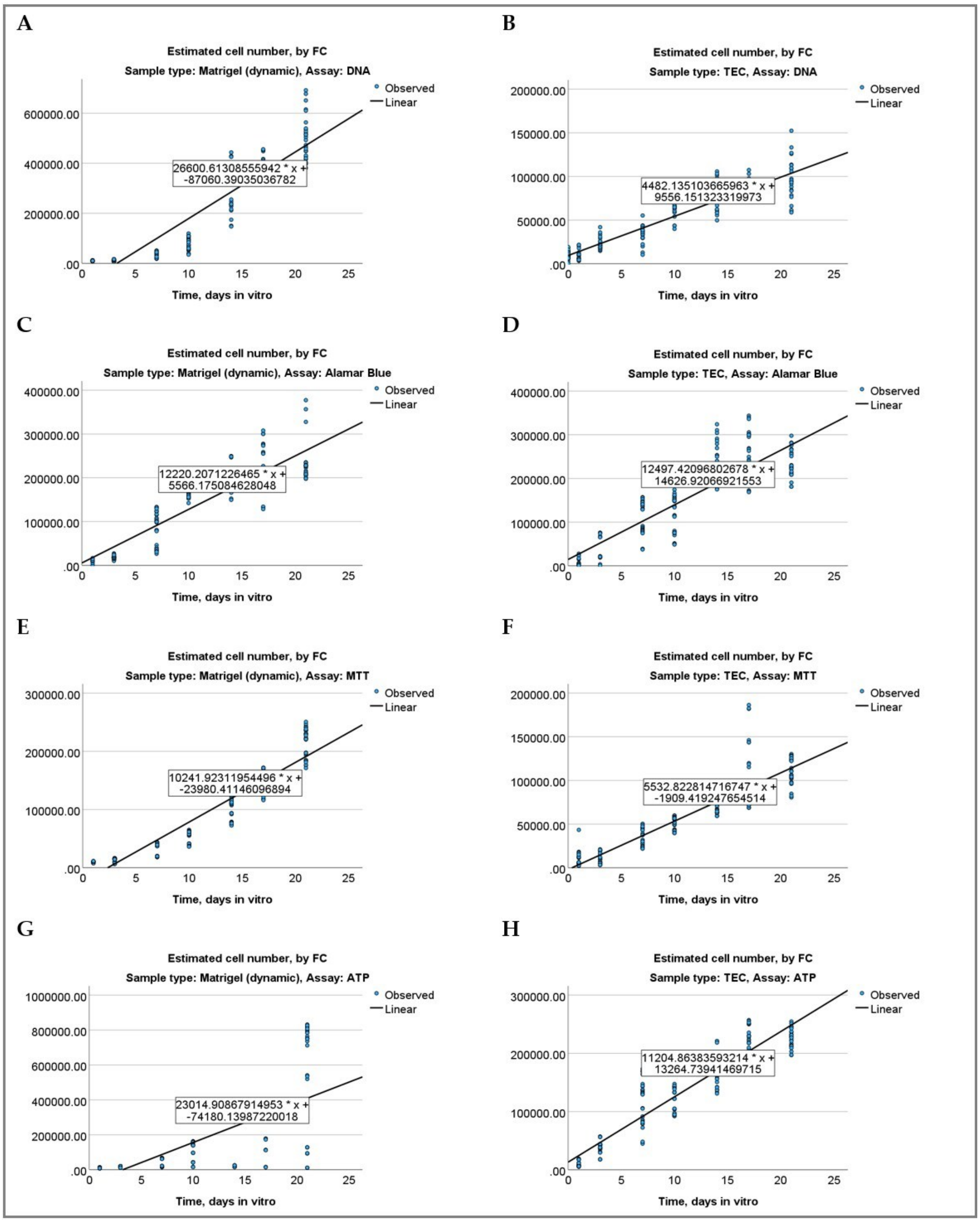


**Figure B2.** Linear regression plots for the assays testing in 3D dynamic Matrigel-based constructs **(A, C, E, G)** and 3D brain dECM-based TECs **(B, D, F, H)** of U251 cells. Blue dots show the data point for the estimated cell numbers calculated by the fold change (FC) of the assay readings normalized to the reading in the smallest cell density sample in the series (shown on the vertical axis), while the time of in vitro culture (measured in days) stands for the independent variable (shown on the horizontal axis). The respective linear regression equations are shown in the inserts on the regression lines. **(A, B)** DNA assay; **(C, D)** Alamar Blue assay; **(E, F)** MTT assay; **(G, H)** ATP assay.

**Table B16. Technical (CVt), biological (CVb) and overall coefficients of variation (CV) of the DNA, Alamar Blue, ATP and MTT assays in dynamic (1-21-day) 3D Matrigel-based constructs and brain dECM-based TECs of U251 cells.**

| Cell culture platform | Assay | | Mean | N | Std. Deviation | Min. | Max. |
|---|---|---|---|---|---|---|---|
| 3D Matrigel (dynamic) | DNA | CVt, % | 12.89 | 7 | 5.38 | 7.01 | 22.91 |
| | | CVb, % | 17.48 | 7 | 10.48 | 1.66 | 28.55 |
| | | CV, % | 23.01 | 7 | 9.77 | 8.75 | 32.25 |
| | Alamar Blue | CVt, % | 16.31 | 7 | 13.047 | 6.40 | 45.25 |
| | | CVb, % | 13.48 | 7 | 14.157 | .00 | 41.71 |
| | | CV, % | 25.37 | 7 | 15.626 | 7.08 | 47.99 |
| | MTT | CVt, % | 5.37 | 7 | 2.19 | 3.19 | 8.71 |
| | | CVb, % | 16.98 | 7 | 7.78 | 8.86 | 30.11 |
| | | CV, % | 18.18 | 7 | 7.73 | 10.50 | 30.78 |
| | ATP | CVt, % | 18.11 | 7 | 10.84 | 3.34 | 34.84 |
| | | CVb, % | 38.94 | 7 | 21.37 | 3.24 | 63.85 |
| | | CV, % | 45.22 | 7 | 18.66 | 16.44 | 65.13 |
| 3D TECs | DNA | CVt, % | 19.41 | 7 | 6.21 | 9.98 | 27.12 |
| | | CVb, % | 12.94 | 7 | 16.34 | .00 | 49.00 |
| | | CV, % | 27.26 | 7 | 15.35 | 10.67 | 58.30 |
| | Alamar Blue | CVt, % | 26.21 | 7 | 15.22 | 10.09 | 54.52 |
| | | CVb, % | 27.86 | 7 | 35.47 | .60 | 93.39 |
| | | CV, % | 41.02 | 7 | 30.35 | 13.49 | 92.32 |
| | MTT | CVt, % | 25.04 | 7 | 16.20 | 8.37 | 54.20 |
| | | CVb, % | 20.21 | 7 | 20.56 | 2.91 | 51.93 |
| | | CV, % | 36.01 | 7 | 26.50 | 13.67 | 83.86 |
| | ATP | CVt, % | 17.45 | 7 | 11.13 | 3.77 | 37.65 |
| | | CVb, % | 12.82 | 7 | 7.40 | 5.49 | 23.59 |
| | | CV, % | 23.12 | 7 | 14.26 | 7.08 | 48.01 |

**Table B17. Pairwise comparison of the technical coefficients of variation (CVt) of the DNA, Alamar Blue, ATP and MTT assays in in dynamic (1-21-day) 3D Matrigel-based constructs and brain dECM-based TECs of U251 cells via the independent-samples Kruskal-Wallis test. Statistically significant comparisons are highlighted.**

| Cell culture platform | Sample 1-Sample 2 | Test Statistic | Std. Error | Std. Test Statistic | Sig. | Adj. Sig. [a] |
|---|---|---|---|---|---|---|
| 3D Matrigel (dynamic) | MTT-DNA | 10.429 | 4.397 | 2.372 | .018 | .106 |
| | MTT-Alamar Blue | 11.429 | 4.397 | 2.599 | .009 | .056 |
| | MTT-ATP | -13.286 | 4.397 | -3.022 | .003 | .015 |
| | DNA-Alamar Blue | -1.000 | 4.397 | -.227 | .820 | 1.000 |
| | DNA-ATP | -2.857 | 4.397 | -.650 | .516 | 1.000 |
| | Alamar Blue-ATP | -1.857 | 4.397 | -.422 | .673 | 1.000 |
| 3D TECs | ATP-DNA | 2.000 | 4.397 | .455 | .649 | 1.000 |
| | ATP-MTT | 4.000 | 4.397 | .910 | .363 | 1.000 |
| | ATP-Alamar Blue | 5.143 | 4.397 | 1.170 | .242 | 1.000 |
| | DNA-MTT | -2.000 | 4.397 | -.455 | .649 | 1.000 |
| | DNA-Alamar Blue | -3.143 | 4.397 | -.715 | .475 | 1.000 |
| | MTT-Alamar Blue | 1.143 | 4.397 | .260 | .795 | 1.000 |

Each row tests the null hypothesis that the Sample 1 and Sample 2 distributions are the same. Asymptotic significances (2-sided tests) are displayed. The significance level is .050. [a.] Significance values have been adjusted by the Bonferroni correction for multiple tests.

**Table B18. Pairwise comparison of the biological coefficients of variation (CVb) of the DNA, Alamar Blue, ATP and MTT assays in in dynamic (1-21-day) 3D Matrigel-based constructs and brain dECM-based TECs of U251 cells via the independent-samples Kruskal-Wallis test. Statistically significant comparisons are highlighted.**

| Cell culture platform | Sample 1-Sample 2 | Test Statistic | Std. Error | Std. Test Statistic | Sig. | Adj. Sig. [a] |
|---|---|---|---|---|---|---|
| 3D Matrigel (dynamic) | Alamar Blue-DNA | 3.714 | 4.397 | .845 | .398 | 1.000 |
| | Alamar Blue-MTT | -3.714 | 4.397 | -.845 | .398 | 1.000 |
| | Alamar Blue-ATP | -11.714 | 4.397 | -2.664 | .008 | .046 |
| | DNA-MTT | .000 | 4.397 | .000 | 1.000 | 1.000 |
| | DNA-ATP | -8.000 | 4.397 | -1.819 | .069 | .413 |
| | MTT-ATP | -8.000 | 4.397 | -1.819 | .069 | .413 |
| 3D TECs | DNA-ATP | -2.571 | 4.396 | -.585 | .559 | 1.000 |
| | DNA-Alamar Blue | -4.071 | 4.396 | -.926 | .354 | 1.000 |
| | DNA-MTT | -4.500 | 4.396 | -1.024 | .306 | 1.000 |
| | ATP-Alamar Blue | 1.500 | 4.396 | .341 | .733 | 1.000 |
| | ATP-MTT | 1.929 | 4.396 | .439 | .661 | 1.000 |
| | Alamar Blue-MTT | -.429 | 4.396 | -.097 | .922 | 1.000 |

Each row tests the null hypothesis that the Sample 1 and Sample 2 distributions are the same. Asymptotic significances (2-sided tests) are displayed. The significance level is .050. [a.] Significance values have been adjusted by the Bonferroni correction for multiple tests.

**Table B19. Pairwise comparison of the overall coefficients of variation (CV) of the DNA, Alamar Blue, ATP and MTT assays in in dynamic (1-21-day) 3D Matrigel-based constructs and brain dECM-based TECs of U251 cells via the independent-samples Kruskal-Wallis test. Statistically significant comparisons are highlighted.**

| Cell culture platform | Sample 1-Sample 2 | Test Statistic | Std. Error | Std. Test Statistic | Sig. | Adj. Sig. [a] |
|---|---|---|---|---|---|---|
| 3D Matrigel (dynamic) | MTT-DNA | 3.571 | 4.397 | .812 | .417 | 1.000 |
| | MTT-Alamar Blue | 3.714 | 4.397 | .845 | .398 | 1.000 |
| | MTT-ATP | -11.857 | 4.397 | -2.697 | .007 | .042 |
| | DNA-Alamar Blue | -.143 | 4.397 | -.032 | .974 | 1.000 |
| | DNA-ATP | -8.286 | 4.397 | -1.884 | .060 | .357 |
| | Alamar Blue-ATP | -8.143 | 4.397 | -1.852 | .064 | .384 |
| 3D TECs | ATP-DNA | 2.571 | 4.397 | .585 | .559 | 1.000 |
| | ATP-MTT | 4.000 | 4.397 | .910 | .363 | 1.000 |
| | ATP-Alamar Blue | 5.714 | 4.397 | 1.300 | .194 | 1.000 |
| | DNA-MTT | -1.429 | 4.397 | -.325 | .745 | 1.000 |
| | DNA-Alamar Blue | -3.143 | 4.397 | -.715 | .475 | 1.000 |
| | MTT-Alamar Blue | 1.714 | 4.397 | .390 | .697 | 1.000 |

Each row tests the null hypothesis that the Sample 1 and Sample 2 distributions are the same. Asymptotic significances (2-sided tests) are displayed. The significance level is .050. [a.] Significance values have been adjusted by the Bonferroni correction for multiple tests.
